# CyteArc: in-place analysis and reuse of remote atlases at 100-million-cell scale

**DOI:** 10.64898/2026.11.24.768888

**Authors:** Gautam Ahuja, Saatvik Viniak, Yi Su, Parashar Dhapola

## Abstract

Public repositories hold hundreds of millions of single-cell profiles, yet analyzing them still requires downloading full files. We present CyteArc, a framework that analyzes remote atlases in place, scaling to nearly 100 million cells on standard CPUs faster than benchmarked workflows, with the lowest memory from 10 million cells upward. Without copying counts, CyteArc integrates millions of cells across studies using 1% to 4% of the disk space of other tools. CyteArc saves every step with full provenance, keeping analyses reproducible and letting researchers branch without rerunning upstream stages. Because those saved steps stay linked, completed cloud atlases act as live references, annotating new datasets without exporting reference files. Using CyteBase, an open cloud repository of over 160 million single-cell and spatial profiles in CyteArc format, we screen over 30 million blood cells across cohorts in minutes and identify rare circulating progenitors whose expansion tracks COVID-19 severity.

---

Curated single-cell atlases serve as quantitative references for annotating cell types, resolving rare states, and testing whether molecular responses generalize across cohorts and diseases [1]. Public repositories such as CZ CELLxGENE Discover [2] now hold more than 90 million unique cells across over 1,500 datasets, and individual perturbation screens approach 100 million cells [3]. Yet most datasets are still shared as single H5AD [4] or RDS files built for local download and in-memory analysis in tools such as Scanpy [5] and Seurat [6]. That file model leaves cloud access, large-scale computation, reference reuse, and analysis tracking split across separate tools (Table 1). Portals such as CELLxGENE Census ease retrieval by letting researchers fetch selected cells and genes. Even so, large queries can still exhaust memory when loaded into external tools for clustering or marker testing, and combining portal studies with local datasets still requires exporting files. Reference mapping adds another divide: Seurat [7] requires exporting separate reference files that can reach tens of gigabytes for large atlases, while platforms such as scvi-hub [8] and ArchMap [9] depend on separately trained machine-learning models. Tracking how results were produced is equally fragmented, because standard tools overwrite saved objects during updates and workflow managers such as Snakemake [10] track scripts and files rather than interactive steps such as cell filters and clustering. As a result, combining studies, annotating cells, and keeping analyses traceable still requires exporting files locally and stitching together separate tools.

**Table 1.**
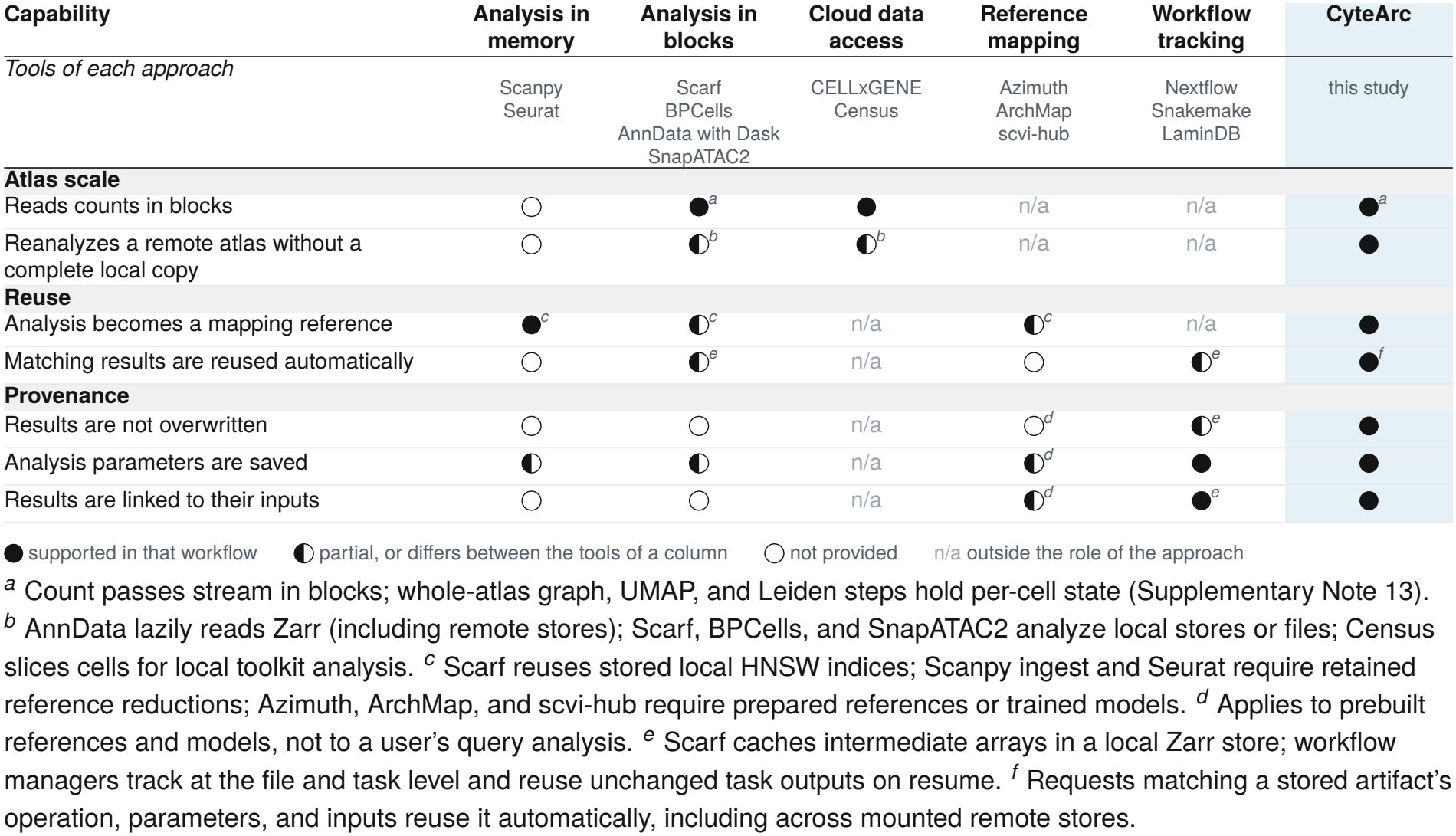
Capabilities for scaling, reusing, and tracing single-cell atlas analyses. Documented workflows of representative tools in each category [2, 4–16].

| Capability | Analysis in memory | Analysis in blocks | Cloud data access | Reference mapping | Workflow tracking | CyteArc |
|---|---|---|---|---|---|---|
| *Tools of each approach* | Scanpy<br>Seurat | Scarf<br>BPCells<br>AnnData with Dask<br>SnapATAC2 | CELLxGENE<br>Census | Azimuth<br>ArchMap<br>scvi-hub | Nextflow<br>Snakemake<br>LaminDB | this study |
| **Atlas scale** |  |  |  |  |  |  |
| Reads counts in blocks | ○ | ●<sup>*a*</sup> | ● | n/a | n/a | ●<sup>*a*</sup> |
| Reanalyzes a remote atlas without a complete local copy | ○ | ◐<sup>*b*</sup> | ◐<sup>*b*</sup> | n/a | n/a | ● |
| **Reuse** |  |  |  |  |  |  |
| Analysis becomes a mapping reference | ●<sup>*c*</sup> | ◐<sup>*c*</sup> | n/a | ◐<sup>*c*</sup> | n/a | ● |
| Matching results are reused automatically | ○ | ◐<sup>*e*</sup> | n/a | ○ | ◐<sup>*e*</sup> | ●<sup>*f*</sup> |
| **Provenance** |  |  |  |  |  |  |
| Results are not overwritten | ○ | ○ | n/a | ○<sup>*d*</sup> | ◐<sup>*e*</sup> | ● |
| Analysis parameters are saved | ◐ | ◐ | n/a | ◐<sup>*d*</sup> | ● | ● |
| Results are linked to their inputs | ○ | ○ | n/a | ◐<sup>*d*</sup> | ●<sup>*e*</sup> | ● |

● supported in that workflow ◐ partial, or differs between the tools of a column ○ not provided n/a outside the role of the approach
<sup>*a*</sup> Count passes stream in blocks; whole-atlas graph, UMAP, and Leiden steps hold per-cell state (Supplementary Note 13).
<sup>*b*</sup> AnnData lazily reads Zarr (including remote stores); Scarf, BPCells, and SnapATAC2 analyze local stores or files; Census slices cells for local toolkit analysis. <sup>*c*</sup> Scarf reuses stored local HNSW indices; Scanpy ingest and Seurat require retained reference reductions; Azimuth, ArchMap, and scvi-hub require prepared references or trained models. <sup>*d*</sup> Applies to prebuilt references and models, not to a user's query analysis. <sup>*e*</sup> Scarf caches intermediate arrays in a local Zarr store; workflow managers track at the file and task level and reuse unchanged task outputs on resume. <sup>*f*</sup> Requests matching a stored artifact's operation, parameters, and inputs reuse it automatically, including across mounted remote stores.

Even after datasets are downloaded locally, scaling analysis to millions of cells forces difficult trade-offs across biological resolution, hardware, and disk layout. Subsampling approaches such as geometric sketching [17] and Seurat v5 [6] keep runtimes manageable by clustering a smaller representative subset of cells and assigning the remaining cells afterward, rather than building the similarity graph across every cell. GPU-based toolkits such as rapids-singlecell [18] analyze every cell at high speed, but require specialized GPU hardware. On CPUs, batched tools such as BPCells [12], Scanpy with Dask [13], and our earlier toolkit Scarf [11] retain every cell by reading expression counts from disk in batches that fit in memory.

However, single-cell workflows scan counts in two directions, by cell for quality control and dimensionality reduction and by gene for marker testing and targeted gene queries. When a tool stores counts in only one direction, gene-level steps must read extra data, transpose matrices in memory, or rely on a manually maintained second copy on disk. Taken together, these access and scaling barriers still force researchers to copy tens of gigabytes locally and depend on high-memory or GPU servers, keeping full-resolution atlas analysis out of reach for most groups.

Here we present CyteArc, an open-source framework that brings remote cloud access, large-scale CPU analysis, reference reuse, and step-by-step history tracking into a single workflow. CyteArc stores datasets in the open Zarr directory format, similar to AnnData, but handles results differently: instead of modifying one object in place, it writes every analysis output as a separate read-only record linked to the exact inputs and settings that produced it. To make public atlases immediately usable in this format, we also built CyteBase, an open cloud collection of 1,372 datasets spanning over 160 million primary single-cell and spatial profiles. Researchers can open a remote CyteBase store by fetching only its lightweight cell and gene tables, pull only the counts each step needs, and join cells across multiple cloud or local stores without copying raw counts. In human blood stores, this remote querying enabled screening across dozens of cohorts to isolate rare circulating hematopoietic progenitors and test their links to disease and severity without downloading the underlying datasets. To scale full-resolution analysis to nearly 100 million cells on standard CPUs without GPUs or subsampling, each store keeps counts ordered by both cell and gene with block sizes configurable at import for dataset scale. A memory planner then schedules batchwise reads and computation within a configured RAM budget and checks larger graph stages upfront before allocating memory.

Saving each step as a read-only record linked to its inputs also changes how researchers explore and reuse completed analyses. During exploratory work, analysts can test new cell filters, clustering resolutions, or parameter choices in separate branches and return to earlier states without repeating unchanged upstream steps. Completed stores also keep their dimensionality reductions, integration parameters, and neighbor indices alongside their counts. As a result, any finished atlas can act directly as a live reference, annotating tens of millions of query cells without exporting separate reference files or rereading reference counts. The same design extends across multiple assays per cell, such as RNA and surface proteins, scaling weighted nearest-neighbor (WNN) integration [7] past a million cells and allowing protein-defined T-cell states to be transferred onto cohorts profiled by RNA alone. Below, we demonstrate CyteArc and CyteBase across remote cross-study discovery, 100-million-cell CPU scaling, iterative reanalysis, reference mapping, and multimodal integration.

## Results

### In-place cloud analysis and virtual composites enable cross-study discovery across public atlases

Although cloud-friendly array formats such as Zarr [19] and TileDB [20] support fetching parts of a dataset over the network, analyzing public single-cell atlases directly in cloud storage remains difficult. Conventional single-file archives such as H5AD [4] and Seurat RDS files [6] bundle raw counts together with metadata and editable results into one local file, while disk-backed engines such as BPCells [12] operate on local storage. A CyteArc store organizes a Zarr directory into three distinct layers: lightweight cell and gene metadata tables, read-only expression count arrays stored both by cell and by gene, and separate read-only analysis outputs (Fig. 1a; Methods). This layout enables an overlay mount, which opens a read-only cloud store in a local workspace by copying only the lightweight metadata tables (1.2% of the 10-million-cell mouse embryo store [21]). Expression counts and stored results stay in the cloud, are streamed on demand, and remain untouched as new outputs are written locally (Supplementary Note 1). To compare cloud and local storage directly on one million mouse embryo cells (8 CPUs; 24 GiB memory budget), opening the cloud store directly ran the full workflow (from quality control through clustering and marker testing) in 10.3 minutes, 2.3-fold faster than cloud Scanpy [5] on Dask [13] and within a minute of local-only BPCells (9.4 minutes, from the multi-tool benchmark below). On local disk, CyteArc finished in 7.1 minutes, 1.3-fold faster than BPCells and 3.9-fold faster than Scanpy on Dask (Fig. 1b; Supplementary Note 1). Researchers can therefore analyze remote cloud atlases at speeds competitive with local-disk tools without storing local copies of the raw counts, making it practical to query public repositories directly in place.

**Figure 1.**
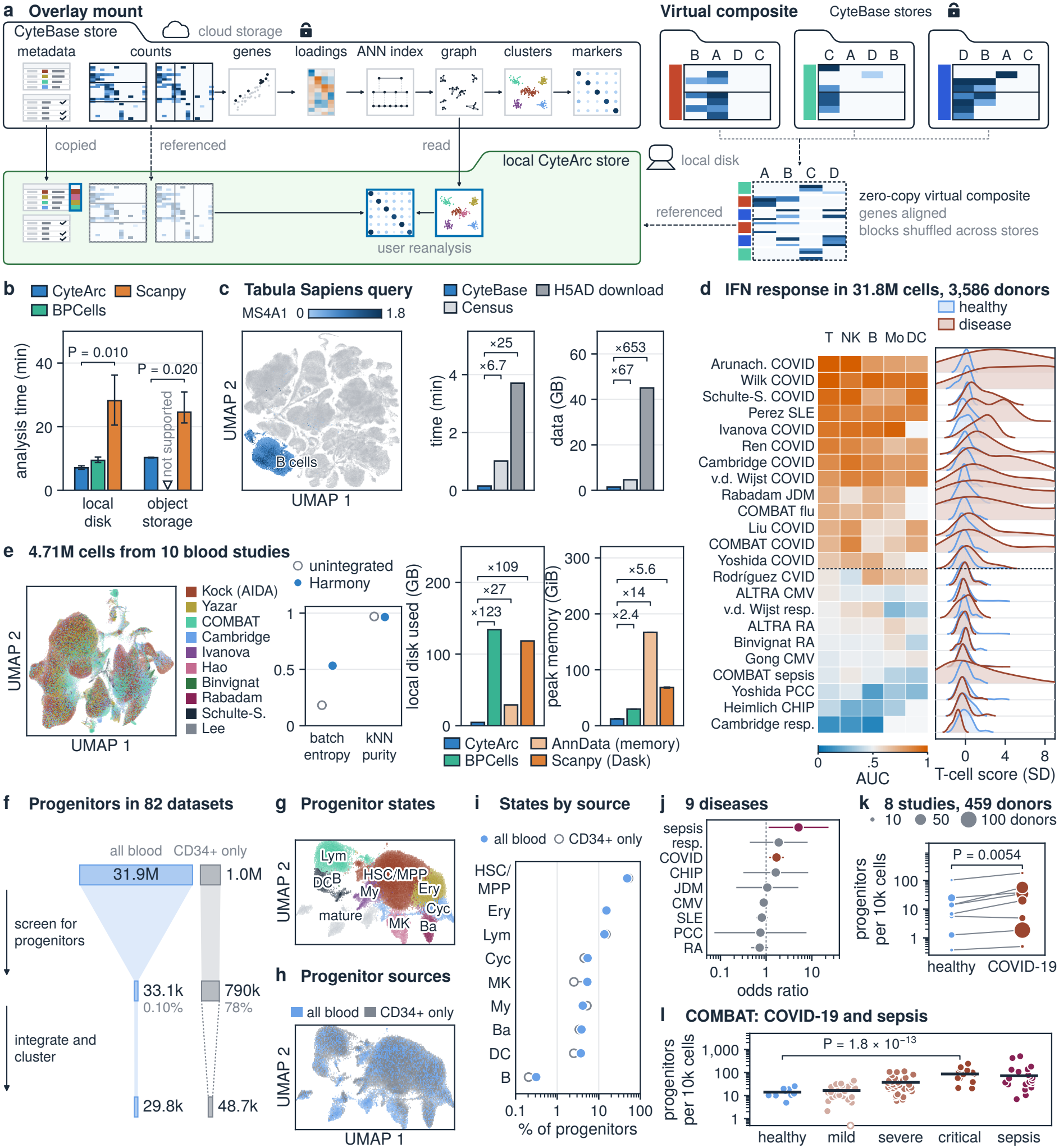
Zero-copy remote mounting, CyteBase queries, and multi-study composites. (**a**) Overlay mount of a read-only cloud store into a local workspace (metadata copied, counts referenced, new outputs local); right, zero-copy composite aligning genes and shuffling cell blocks. (**b**) Runtime on 1M embryo cells (8 CPUs) from local disk or cloud storage (bars, means of n = 3; error bars, ranges; brackets, Welch P; CyteArc versus BPCells on local disk, P = 0.017; open triangle, local-only). (**c**) Cloud Tabula Sapiens UMAP with 114,495 B cells colored by MS4A1; right, query time and bytes versus CELLxGENE Census and H5AD download. (**d**) Donor interferon scores across 23 disease-versus-healthy contrasts from 18 of the 71 scored CyteBase blood stores: per-lineage AUROC (left) and T-cell score densities (right). (**e**) Harmony UMAP of the 10-study blood composite (4.71M cells) colored by study; middle, batch entropy and kNN purity; right, local disk for counts (the composite’s index against the baselines’ count copies) and peak memory of four routes (memory: means and ranges, n = 3). (**f**) In-place screen of 82 CyteBase blood stores for circulating HSPCs (band width *∝* cells^1*/*2^). (**g**) UMAP of 83,086 integrated cells from 64 stores colored by progenitor state (mature leukocytes, grey). (**h**) Same UMAP colored by source: unselected blood (blue) versus CD34+-enriched stores (grey). (**i**) Progenitor state shares in unselected blood (filled) versus CD34+-enriched stores (open; log axis). (**j**) Odds ratios (95% CIs) of blood progenitors in nine diseases versus healthy controls: random-effects pools for COVID-19, cytomegalovirus infection, and rheumatoid arthritis, single-study estimates otherwise (dashed, OR = 1). (**k**) Progenitors per 10,000 blood cells in healthy and COVID-19 donors of eight studies (area *∝* donors; bracket, P value). (**l**) Progenitors per 10,000 cells per COMBAT donor by cohort (bars, pooled rates; bracket, trend P value).

To make public single-cell atlases directly analyzable in the cloud, we converted 1,372 RNA datasets (including 639 spatial datasets) from CZ CELLxGENE Discover [2] into cloud-hosted CyteArc stores. Released openly as CyteBase on Hugging Face, this resource spans 163.6 million primary observations across nine species in 1.09 TB of cloud storage (Supplementary Fig. 1a; Supplementary Note 2; Methods). Because CyteBase stores maintain both cell-ordered and gene-ordered arrays, targeted queries fetch only the requested genes and cells rather than transferring gigabytes over the network. On a cloud-hosted Tabula Sapiens CyteArc store [22, 23], retrieving UMAP [24] coordinates, cell-type labels, and B-cell MS4A1 counts took 9.0 seconds on first access and 2.6 seconds on repeated queries. That query ran 6.7-fold faster with 67-fold less network traffic than running the same gene query in CELLxGENE Census (even without UMAP coordinates), and transferred hundreds of times less data than downloading the source H5AD file (Fig. 1c; Supplementary Note 2).

Because selective gene reads transfer only a small fraction of the combined archive, researchers can screen multi-gene biological programs across entire public repositories in minutes. A 10-gene screen across all 82 human blood-containing stores in CyteBase transferred only 4.7% of their combined cell- and gene-ordered count arrays (Supplementary Fig. 1b). To test this capability on a biological signature, we scored an 18-gene type I interferon program across all unselected blood stores (excluding sorted, enriched, or stimulated datasets; Methods), covering 31.8 million primary cells from 3,586 donors in 18.1 minutes of combined runtime. Across all 13 viral and autoimmune interferon-driven contrasts (10 COVID-19 cohorts plus systemic lupus erythematosus, juvenile dermatomyositis, and influenza, each versus within-study healthy controls), donor interferon scores ranked disease donors above healthy controls in every evaluated immune lineage (AUROC 0.55 to 1.00). In T cells and dendritic cells, every interferon-driven disease contrast achieved a higher AUROC than every non-interferon control condition evaluated in that lineage (AUROC 0.64 to 1.00 versus 0.12 to 0.54 in T cells; 0.74 to 0.94 versus 0.29 to 0.63 in dendritic cells; Fig. 1d; Supplementary Note 3). Selective cloud reading thus turns cross-study signature validation from downloading full archives across dozens of studies into a routine, minutes-long check across tens of millions of cells.

Combining datasets across studies normally requires downloading each study, matching their gene lists onto a shared set of genes, and concatenating their raw count matrices into a new copy in memory or on disk before integration can begin. CyteArc avoids copying raw counts through a virtual composite (Fig. 1a; Methods): because independent studies often profile different genes in different orders, the virtual composite builds a small local index that maps shared genes across read-only source stores into one common order and reads cell batches directly from those stores on demand. To verify that a virtual composite matches a combined local store, we tested a four-study COVID-19 cohort and confirmed that it reproduced the exact Leiden [25] clusters (adjusted Rand index, ARI = 1.0) and principal component analysis (PCA) subspace of that local store (Supplementary Note 4). Scaling to 10 human blood studies totaling 4.71 million primary cells and 18,446 shared genes, CyteArc built the virtual composite as a 1.09 GB local index at creation (15.2 GB after clustering) without copying counts and clustered the cohort at a mean peak memory of 12.2 GiB. In a Harmony2 [26, 27] integration of the same 10-study composite, mixing across studies within each cell type nearly tripled (0.18 to 0.53) while local cell-type purity was preserved (0.970 versus 0.965; Fig. 1e; Supplementary Note 4). We benchmarked this virtual composite against in-memory AnnData [4] concatenation, on-disk AnnData concatenation with Scanpy [5] on Dask [13], and BPCells [12], giving every comparator local H5AD files that were already filtered to the same cells and aligned to the shared genes. Even with that head start, the virtual composite’s index required 27- to 123-fold less local disk than the comparators’ count copies, and peak memory was 2.4- to 14-fold lower (Fig. 1e; Supplementary Note 4).

Combining in-place repository screening with virtual composites makes it possible to resolve rare cell populations that individual blood cohorts capture only sparsely, such as circulating hematopoietic stem and progenitor cells (HSPCs; Supplementary Note 6). Scanning all 82 human blood-containing stores in CyteBase (31.9 million unselected peripheral-blood cells plus CD34+-enriched reference stores) in 8.1 minutes flagged candidate cells carrying author progenitor labels or co-expressing at least two of four canonical progenitor markers (CD34, SPINK2, PROM1, and CRHBP), accounting for 0.10% of unselected peripheral blood (Fig. 1f). To avoid repeatedly streaming such sparse 0.10% subsets over the network, we extracted the blood candidates together with sampled CD34+-enriched reference cells into compact local subsets (across eight parallel jobs, the longest taking 34.6 minutes), then joined and analyzed the 83,086 extracted cells across 41 qualifying studies in a virtual composite using Harmony2 integration and Leiden clustering in 7.5 minutes at 7.0 GiB peak memory. Clustering separated residual mature leukocytes from 18 progenitor clusters across nine marker-defined states spanning multipotent, lineage-primed, and cycling compartments (Fig. 1g). Across all nine states, circulating progenitors from unselected peripheral blood differed from CD34+-enriched reference shares by at most 3.5 percentage points, with the three largest shares in hematopoietic stem and multipotent progenitor (HSC/MPP), erythroid, and lymphoid-primed states (Fig. 1h; Fig. 1i). Thus, progenitors circulating in unselected human blood span the full multipotent, lineage-primed, and cycling hematopoietic hierarchy rather than representing a single mobilized subset.

Because screening CyteBase in place extracts uniform donor-level measurements across dozens of public cohorts without downloading them, it allowed us to test whether circulating progenitor expansion reported in single COVID-19 studies [28, 29] generalizes across cohorts and distinguishes COVID-19 from chronic inflammatory conditions across 1,234 donors, nine diseases, and 15 studies with internal healthy controls (Fig. 1j; Supplementary Note 7; Methods). In a random-effects meta-analysis across eight COVID-19 studies, circulating progenitor odds were elevated 1.69-fold relative to healthy controls (P = 0.0054, Benjamini-Hochberg q = 0.049), rising in seven of eight cohorts with consistent effect sizes and holding after adjusting for lymphocyte sequencing depth or jointly for depth and donor sex (Fig. 1k; Supplementary Note 7). A single sepsis cohort also showed elevated progenitor odds (odds ratio 5.12, P = 0.034, q = 0.15), whereas progenitor odds were slightly lower rather than elevated across six chronic viral, autoimmune, post-infectious, and clonal conditions (pooled odds ratio 0.86, 95% CI 0.77 to 0.97, weighted mainly by three cytomegalovirus cohorts; COVID-19 versus chronic pool 1.95-fold, P = 0.00068; Fig. 1j). Across five multi-severity COVID-19 cohorts, progenitor odds rose on average 1.85-fold per severity tier (healthy through severe), and within the uniformly graded COMBAT cohort they climbed monotonically at 2.05-fold per grade from healthy volunteers to critical disease (Fig. 1l; Supplementary Note 7). Together, these remote queries, cross-study screens, and virtual composites depend on CyteArc’s core analysis engine; we next examine how its cell- and gene-ordered storage layout and memory planner scale full-resolution analysis from 1 million to nearly 100 million cells on standard CPUs.

### Bounded-memory batchwise processing scales single-cell analysis to 100 million cells

Scaling full-resolution analysis from 1 million to nearly 100 million cells requires keeping memory within a fixed budget while ensuring that read requests and runtime grow at most linearly with cell count. Because standard workflows read expression counts both by cell (for quality control and PCA) and by gene across all cells (for variable-gene selection, scaling, and marker testing; Fig. 2a), scaling them exposes three distinct bottlenecks. First, in-memory tools such as Scanpy [5] and Seurat [6] exhaust RAM or hit R’s 32-bit sparse-matrix index limits, while single-orientation disk-backed workflows must either unpack unneeded entries during gene-wise scans or create a transposed copy in RAM (as in Scanpy on Dask [13]). Second, even when counts are stored in gene-ordered chunks, keeping fixed file-shard and chunk sizes causes read requests to grow quadratically as datasets expand: under a fixed memory budget, doubling the cell count halves how many genes fit in memory per batch (doubling the number of gene batches) while doubling how many cell chunks each gene spans, multiplying total read requests fourfold. Third, even after batchwise count processing is bounded, downstream neighbor-graph construction, embedding, clustering, and marker testing can still time out, crash, or run out of memory at multi-million-cell scale when individual stages scale poorly or create unbounded intermediate arrays in memory.

**Figure 2.**
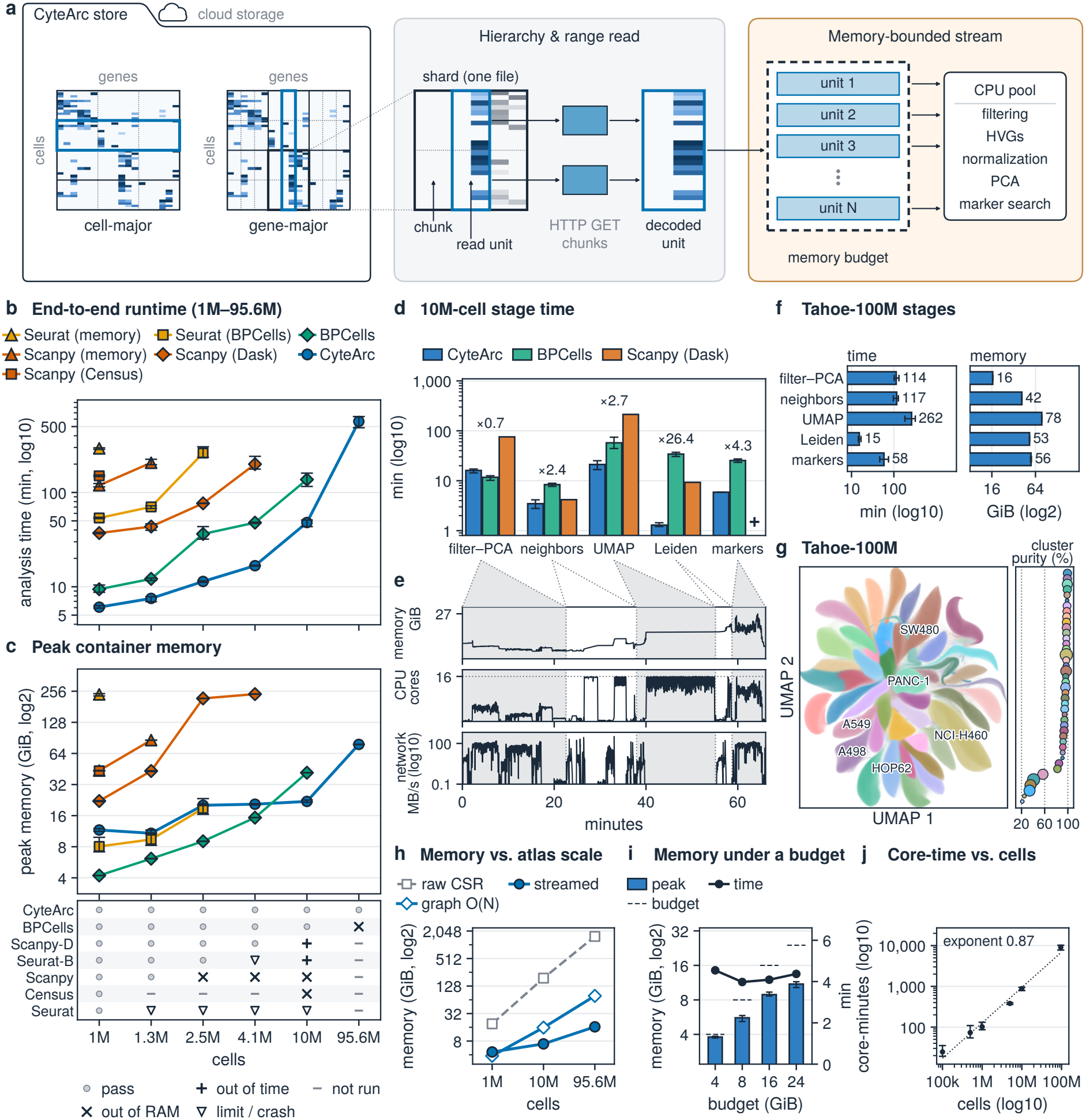
Bounded-memory analysis scaling from 0.1M to 95.6M cells. (**a**) CyteArc store holding cell-major and gene-major compressed chunks in shards, processed batchwise by the memory planner within a memory budget. (**b**) End-to-end analysis runtime across five 1M-10M-cell atlases and Tahoe-100M (95.6M cells; Tabula Sapiens in Extended Data Table 1; points, means; error bars, ranges). (**c**) Peak container memory of the runs in b (CyteArc at 1M-10M cells, whole runs including store import; on Tahoe-100M, analysis only, with store preparation peaking at 128.5-129.6 GiB), with run outcomes below; BPCells on Tahoe-100M ran out of memory in Leiden clustering. (**d**) Stage runtimes on 10M embryo cells for CyteArc, BPCells, and Scanpy on Dask (numbers, BPCells/CyteArc time ratio; +, timed out; bars, means; error bars, ranges; Scanpy on Dask, mean of its two incomplete attempts). (**e**) One-second container memory, active CPUs, and network receive rate during a separate 10M-cell cloud-storage run (shading, stages). (**f**) Stage runtime (left) and peak container memory (right) on Tahoe-100M on 16 CPUs (bars, means of n = 3; error bars, ranges). (**g**) UMAP of 83.9M filtered Tahoe-100M cells, each pixel colored by its commonest cell line (six largest named); right, dominant cell-line purity per Leiden cluster (area, cluster size). (**h**) Uncompressed sparse-matrix size (dashed) versus peak memory of the batchwise variable-gene, normalization, and PCA passes and of per-cell graph stages at 1M and 10M cells (means of n = 3) and 95.6M cells (first run). (**i**) Peak container memory (bars, means of n = 3; error bars, ranges) and runtime (line, right axis) from import through PCA on 1M cells under 4-24 GiB memory budgets (dashed). (**j**) Analysis core-time (minutes × CPUs) versus cells across 0.1M-10M embryo cells and Tahoe-100M (n = 3 per size; error bars, ranges; dotted, power-law fit).

CyteArc addresses all three bottlenecks by pairing cell-ordered (cell-major) and gene-ordered (gene-major) Zarr arrays with batchwise readers, a memory planner, and multi-threaded downstream graph, embedding, clustering, and marker stages (Fig. 2a; Methods). Saving both orientations at import adds a median of 75% to cell-major storage across CyteBase rather than decoding 91 bytes (90 of them unneeded) for every byte used during gene-wise passes at 10 million mouse embryo cells (Supplementary Note 8).

To prevent quadratic growth in read requests at larger scales, gene-major file shards can be configured at import so each shard spans more cells across fewer genes; this restores linear scaling in both read requests and pass times, cutting variable-gene read requests 7.8-fold on the 10-million-cell embryo store (Supplementary Fig. 2a; Supplementary Fig. 2b) and 7.9-fold on Tahoe-100M, where the variable-gene pass dropped from 196 to 43 minutes with identical outputs (Supplementary Note 8). To turn this layout into sustained throughput, especially when reading from cloud storage, CyteArc uses batchwise readers guided by the memory planner: they size active read buffers to stay within the configured memory budget and fetch upcoming chunks ahead of time so disk and network transfers overlap with multi-core CPU computation. Together, these layout and execution choices let CyteArc analyze every cell on standard CPUs without relying on GPUs (as in rapids-singlecell [18]) or subsampling (as in geometric sketching [17] and Seurat v5 [6]).

To test how this architecture performs in practice, we benchmarked CyteArc against six widely used single-cell workflows end to end, from quality control and PCA through neighbor-graph construction, UMAP [24], Leiden clustering [25], and all-cluster Wilcoxon marker testing across six atlases ranging from 1 to 10 million cells (Fig. 2b; Fig. 2c; Extended Data Table 1; Supplementary Note 9). Five of the seven workflows failed to complete this range even when retried on cloud instances with up to 336 GiB of RAM: in-memory Seurat [6] hit R’s 32-bit sparse-matrix index limit above 1 million cells, in-memory Scanpy [5] and Scanpy on CELLxGENE Census [2] ran out of memory while loading or filtering cells, Scanpy on Dask [13] ran batchwise through PCA but timed out or ran out of memory after creating a transposed marker matrix in RAM, and Seurat on BPCells [12] counts crashed during UMAP initialization or timed out in Leiden clustering (Fig. 2c; Extended Data Table 1; Supplementary Note 9). Only native BPCells and CyteArc completed all six atlases. Across the five atlases shown in Fig. 2b, CyteArc ran 1.5- to 3.2-fold faster than BPCells (and 1.3-fold faster on Tabula Sapiens), cutting total runtime including one-time store preparation by 31% to 57% across those five atlases (10% to 57% across all six). On the 10-million-cell mouse embryo atlas, CyteArc finished in 48.0 minutes compared with 2.3 hours for BPCells, while staying on 32 GiB cloud instances (24 GiB memory budget) as BPCells peaked at 41.6 GiB (Fig. 2c; Supplementary Note 9). This speed advantage held when replacing the benchmark’s Gaussian quality filter (which keeps 5% to 10% fewer cells) with the comparators’ exact cell filter and marker gene set, where CyteArc remained 1.2- to 2.7-fold faster across the four atlases tested (Supplementary Note 9).

Breaking down the 10-million-cell mouse embryo run by stage showed that fast batchwise count processing alone does not determine end-to-end speed: BPCells completed cell filtering through PCA faster than CyteArc (11.7 versus 16.1 minutes). Instead, the majority of total runtime was spent in downstream neighbor-graph construction, UMAP embedding, clustering, and all-cluster marker discovery. In those stages, CyteArc’s parallel neighbor search (2.4-fold faster), UMAP (2.7-fold), deterministic multi-threaded Leiden clustering (26.4-fold), and batchwise Wilcoxon marker tests (4.3-fold) compressed more than two hours of downstream computation into roughly half an hour (Fig. 2d). Furthermore, one-second resource monitoring from a separate 10-million-cell run read directly from cloud storage showed how data transfer and compute trade off across stages: count passes fetched remote batches concurrently with multi-core computation (reaching 152 MB/s during filtering through PCA and 144 MB/s during marker search), downstream graph and embedding stages used up to 15.9 of 16 CPU cores (with brief network bursts up to 327 MB/s during neighbor search), and peak memory stayed well below budget (27.2 GiB under a 48 GiB budget; Fig. 2e; Supplementary Note 11).

Scaling an order of magnitude further, CyteArc streamed the complete workflow, including all-cluster Wilcoxon marker discovery, directly from cloud storage on the 95.6-million-cell Tahoe-100M chemical perturbation atlas [3] (1.52 TiB uncompressed sparse counts). Running each workflow stage as a job on 16 standard CPUs with 256 GiB of RAM (192 GiB budget), end-to-end analysis averaged 9.4 hours across three replicates (peaking at a mean of 78.3 GiB during UMAP) following an average 4.0-hour store preparation, whereas native BPCells ran out of memory during Leiden clustering on a 336 GiB cloud instance (Fig. 2b; Fig. 2c; Fig. 2f; Supplementary Note 12). Retaining all 83.9 million filtered cells at full resolution, multi-threaded Leiden clustering finished in a mean of 15.3 minutes. In the resulting 43 clusters (first replicate), 35 drew at least 90% of their cells from a single perturbed cancer cell line, with 96% median single-line purity across the 43 clusters (Fig. 2g). Full-resolution analysis of perturbation screens approaching 100 million cells therefore runs end to end on 16-CPU, 256 GiB cloud instances directly from cloud storage, without multi-GPU hardware or cell subsampling.

Across this nearly hundredfold jump from 1 million to 95.6 million cells, sizing cell and gene batches to a configured memory budget (24 GiB on the embryo atlases and 192 GiB on Tahoe-100M) decoupled count-pass memory from matrix size. Even as uncompressed sparse counts grew more than 80-fold (from 19.1 GiB at 1 million embryo cells to 1.52 TiB on Tahoe-100M), batchwise variable-gene selection, normalization, and PCA stayed at or below 16.5 GiB (Fig. 2h). Analysis peak memory was instead governed either by stages that deliberately use the configured budget to maximize throughput (such as all-cluster marker search, peaking at a mean of 22.0 GiB under a 24 GiB budget at 10 million cells) or by stages whose internal data structures scale with cell count (such as UMAP, peaking at a mean of 78.3 GiB on Tahoe-100M). Lowering the configured budget (and scaling down batch and chunk targets at 8 and 4 GiB) across 24, 16, 8, and 4 GiB on 1 million embryo cells kept peak memory strictly below every limit while import-to-PCA runtime remained stable at 4.0 to 4.5 minutes (Fig. 2i; Supplementary Note 13). Before starting each step, CyteArc also estimates whether its required memory fits within the budget, stopping cleanly with the required memory estimate rather than crashing mid-run: under an 8 GiB budget at 10 million cells, when each stage was run separately, count and neighbor-graph stages completed while Leiden clustering stopped upfront before allocating memory; in a continuous end-to-end run, UMAP similarly stopped upfront (Supplementary Note 13). Across the nearly thousandfold span from 100,000 embryo cells to 95.6 million Tahoe-100M cells, total analysis core-time (runtime multiplied by CPUs) scaled with a scaling exponent of 0.87 overall and 1.00 (linear) across the 1-million-cell embryo to 95.6-million-cell Tahoe-100M range (Fig. 2j; Supplementary Note 13). Once full-resolution analysis scales linearly on standard CPUs, the next bottleneck shifts from running a pipeline once to iterating on analysis parameters, cell subsets, and reference versions, reusing finished atlases as mapping references, and integrating additional modalities without redoing upstream work.

### Provenance-linked artifacts support iterative reanalysis, million-cell reference mapping, and multimodal integration

Exploratory single-cell analysis is iterative: researchers repeatedly adjust clustering resolutions, neighbor graphs, cell subsets, or batch-correction settings [30]. In standard in-memory tools, modifying an object in place overwrites prior results or forces researchers to manage separate file copies and manually track which upstream steps must be rerun [5]. CyteArc instead records each stage output as a read-only artifact linked to its operation, parameters, and upstream input fingerprints, forming a provenance graph inside the store (Fig. 3a; Methods). Researchers can therefore rerun the same workflow with modified settings without manually skipping steps or juggling separate copies: CyteArc automatically reuses any previously computed local or cloud artifact whose parameters and inputs are unchanged, runs only stages downstream of a change, and keeps every earlier branch accessible in the same store (Supplementary Note 14). On a mounted 10-million-cell mouse embryo store [21], running the workflow with the initial parameters retrieved the stored clustering directly in 19.3 seconds without recomputing a single step or duplicating any data locally. Changing only the Leiden clustering resolution on 8 CPUs automatically reused the remote neighbor graph, finishing in 2.3 to 3.3 minutes and writing only 6.0 to 7.1 MB of new cluster labels locally (versus 29.7 minutes from cell filtering through Leiden in the 16-CPU initial run; Fig. 3a; Supplementary Fig. 3a). Similarly, changing a parameter earlier in the pipeline (increasing the neighbor count to 21) automatically skipped count preprocessing, PCA, and neighbor-index construction, recomputing only the steps downstream of the stored index (neighbor search, graph construction, and clustering) in 11.7 minutes and writing 2.79 GB of local neighbor, graph, and cluster artifacts rather than copying the 40.2 GB source H5AD. Even when carrying a neighbor-count change (11 to 15 neighbors) all the way through UMAP and all-gene marker discovery on a 10-million-cell embryo run, automatic reuse of preprocessing, PCA, and the neighbor index cut full-pipeline runtime by 27% (Supplementary Fig. 3b). The same automatic reuse extends to lineage subsetting, adding batch correction, and updating cell-type annotations as reference atlases grow. Isolating and clustering 272,930 endothelial cells from the mounted 10-million-cell embryo atlas across six parameter combinations used only 1.72 GB in the local user store (before the whole-atlas branches above), 23-fold smaller than the 40.2 GB source H5AD (Supplementary Fig. 3c). Adding Harmony2 batch correction [26, 27] to an already-analyzed four-study COVID-19 virtual composite reused the saved PCA to cut runtime by 32% while avoiding 2.56 GB of remote count transfers (Supplementary Fig. 3d; Supplementary Fig. 3e). Likewise, when the healthy Human Lung Cell Atlas core (584,944 cells [31]) was updated into a new reference version by adding 239,707 interstitial lung disease and control cells to include disease-associated aberrant basaloid annotations [32], CyteArc updated cell-type assignments on an independent pulmonary fibrosis cohort [33] in place (adding 61 MB versus 157 MB for rerunning both versions in a fresh store) while keeping the earlier version’s assignments recoverable in 31 seconds without re-reading the reference (Supplementary Fig. 3f; Supplementary Note 14).

**Figure 3.**
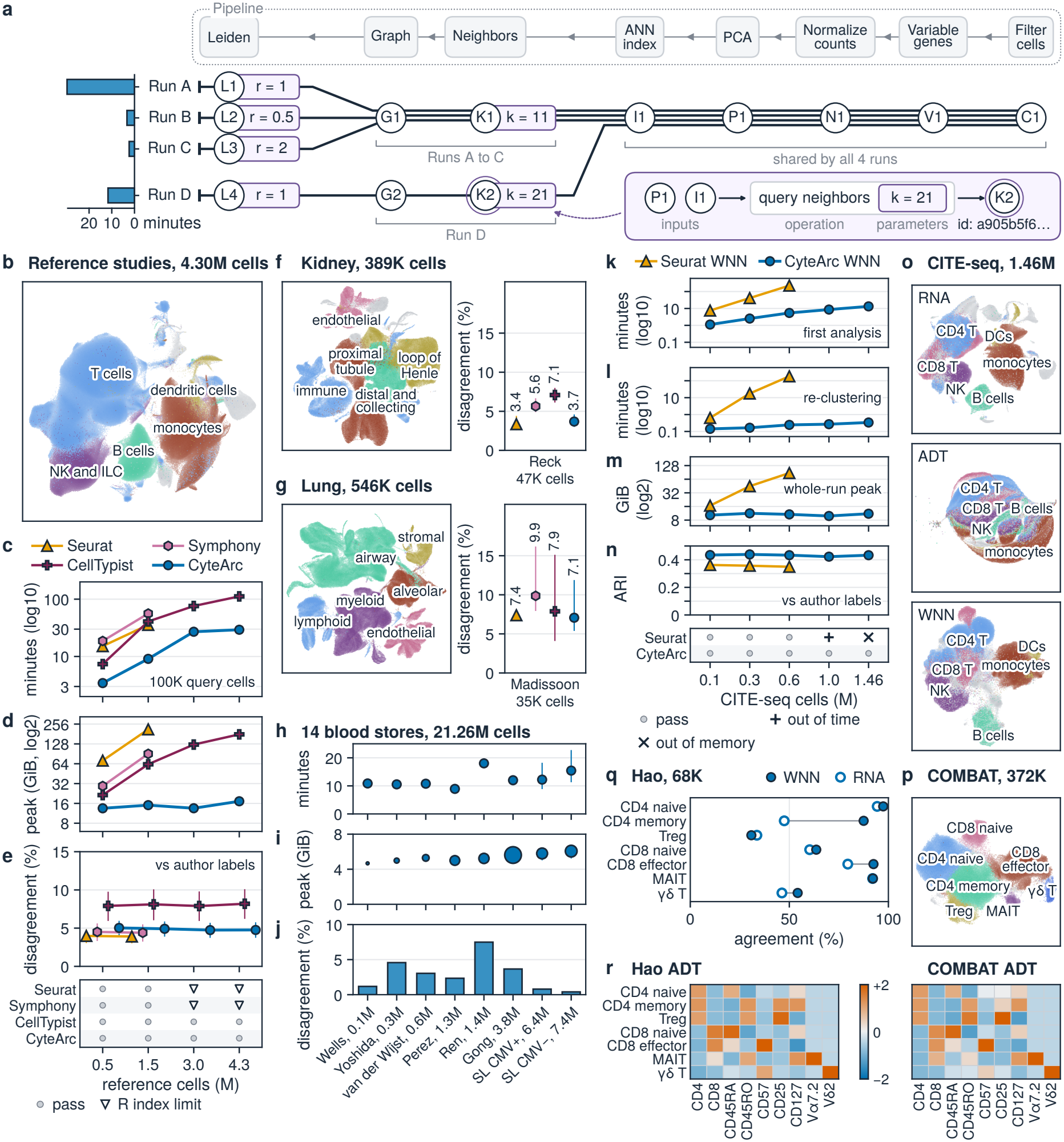
Provenance-tracked branching, reference mapping, and multimodal integration. (**a**) Four Leiden runs on a mounted 10M-cell embryo store sharing upstream artifacts; bars, newly computed stage minutes; inset, provenance record of K2 with its artifact ID. (**b**) Cells of the seven reference studies in the Harmony2 UMAP of the ten-study blood composite, colored by cell type. (**c**) Time to build 0.50M-4.29M-cell blood references and map 100,002 query cells (n = 1 per point; CellTypist with 2,000 variable genes in c-e). (**d**) Peak container memory of the runs in c. (**e**) Label disagreement on the ladder query (bars, donor-bootstrap 95% CIs; bottom, outcomes). (**f**) Kidney reference UMAP (388,878 cells) and disagreement on Reck et al. (donor-bootstrap 95% CIs; CellTypist with all genes in f and g). (**g**) Human Lung Cell Atlas core UMAP (546,358 cells) and disagreement on Madissoon et al. (donor-bootstrap 95% CIs). (**h**) Runtime per job to map 14 CyteBase blood stores (21.3M cells, six studies in seven CELLxGENE collections) onto the stored 4.29M-cell reference. (**i**) Peak container memory per store in h (area ∝ cells). (**j**) Label disagreement per job across cells in the nine reference types. (**k**) Initial workflow runtime on 0.10M-1.46M CITE-seq blood cells for CyteArc WNN and Seurat (n = 1 per point). (**l**) Runtime to recluster each stored WNN graph at resolution 1.2. (**m**) Whole-run peak container memory across the CITE-seq series. (**n**) Cluster agreement (ARI) with author labels at 19 clusters, the number of author cell types (points, means of 3 Leiden seeds; bars, ranges; bottom, outcomes). (**o**) RNA, ADT, and WNN UMAPs of all 1.46M CITE-seq cells after Harmony2 study correction, colored by cell type. (**p**) WNN UMAP of 371,747 COMBAT T cells across seven subtypes. (**q**) Subtype agreement of Hao T cells mapped by RNA onto COMBAT WNN (filled) versus RNA (open) clusters. (**r**) Scaled median surface-protein CLR per subtype in RNA-mapped Hao T cells (left) and COMBAT T cells (right).

Reusing an analyzed atlas for nearest-neighbor label transfer is fragile in standard workflows: matching a query to a reference requires keeping the exact gene-scaling statistics, PCA loadings, batch-correction centroids, and neighbor index synchronized, which typically forces teams to export multi-gigabyte standalone reference files [7] or train a separate classifier [34]. Because CyteArc’s provenance graph already stores and links these stage artifacts (including Harmony2 centroids [26, 27]), any stored atlas analysis can be used directly as a mapping reference without risking mismatched preprocessing or duplicating data (Methods). Marking an existing analysis as a mapping reference writes only a 0.5 to 0.6 MB index file linking those stored artifacts in seconds, rather than exporting a separate reference file (up to 76.5 GB for Seurat and 2.10 GB for R Symphony at 1.50 million blood cells; Supplementary Fig. 4a; Supplementary Fig. 4b; Supplementary Note 15). Across seven-study human blood references spanning 0.50 to 4.29 million cells (Fig. 3b), CyteArc built each reference from raw counts and mapped 100,002 held-out query cells in 3.4 to 29.3 minutes (Fig. 3c) while keeping peak memory near constant at 13.5 to 17.2 GiB (Fig. 3d). At 1.50 million cells, CyteArc finished in 9.1 minutes and 15.1 GiB, versus 35.6 minutes and 212.7 GiB for Seurat and 56.1 minutes and 90.5 GiB for R Symphony; neither R tool could run at 3.00 or 4.29 million cells because their in-memory sparse matrices exceeded R’s 32-bit entry limit. CellTypist similarly ran out of memory at 1.50 million cells on all genes and required up to 178.6 GiB at 4.29 million cells even when restricted to 2,000 variable genes. Once a reference is already built, annotating a new 100,002-cell query in CyteArc took only 31 to 68 seconds across all four reference sizes, running 13- to 22-fold faster than querying a saved Seurat reference and 29- to 79-fold faster than R Symphony at matched sizes (Supplementary Fig. 4c). This speed and memory efficiency preserved nearest-neighbor label quality: CyteArc stayed within 0.5 to 1.0 percentage points of Seurat and R Symphony in label disagreement (4.7% to 5.0% versus 3.9% to 4.0% and 4.4% to 4.5%, and below variable-gene CellTypist at 7.9% to 8.2%; Fig. 3e) and achieved the highest recall on rare cell types below 1% of the reference pool (60.8% to 61.5%, versus 58.4% to 58.8% for Seurat and 45.8% to 46.2% for R Symphony; Supplementary Fig. 4d). Enabling Harmony2 batch correction with Symphony query projection [35] kept the same bounded memory, fast stored-reference mapping, and comparable label agreement (Supplementary Note 15).

We next tested whether this mapping performance extends beyond blood to solid-organ atlases. When mapping an independent kidney study onto a two-study kidney reference [36–38], CyteArc matched Seurat within 0.4 percentage points and improved on R Symphony and CellTypist (3.7% disagreement versus 3.4%, 5.6%, and 7.1%; Fig. 3f). When mapping an independent three-donor lung study onto the remaining Human Lung Cell Atlas core [31, 39], CyteArc had 7.1% disagreement, comparable to Seurat (7.4%), CellTypist (7.9%), and R Symphony (9.9%) with overlapping donor-level confidence intervals (Fig. 3g). To demonstrate large-scale annotation against a remotely hosted atlas without copying reference or query datasets locally, we used a single cloud-hosted 4.29-million-cell blood reference to annotate 21.3 million primary cells across 14 remote CyteBase blood stores in place. Streaming query counts from remote storage while writing only the new annotations locally took 6.9 to 20.7 minutes and 4.7 to 6.5 GiB of peak memory per store (Fig. 3h; Fig. 3i). Across the 21.1 million cells carrying labels in the nine reference types, transferred labels matched author annotations on 98.2% of cells overall, with 0.4% to 7.5% disagreement across the eight cohort runs and 95.9% pooled agreement across the six smaller cohorts (Fig. 3j). Finally, CyteArc scores every transferred label by neighbor-vote confidence, flagging uncertain or perturbation-shifted cells rather than forcing low-confidence labels onto uncertain or shifted cell states. When mapping interferon-beta-stimulated PBMCs onto unstimulated controls [40], stricter confidence filtering abstained on perturbation-shifted cells (increasing the unassigned share from 2.9% at 0.5 confidence to 59.6% at 0.9) while raising label agreement among assigned cells from 85.1% to 98.1% (Supplementary Note 16).

Single-cell studies increasingly pair RNA with additional modalities from the same cells, including surface proteins (CITE-seq) [41], chromatin accessibility [42], and sample hashtags [43]. Multimodal containers such as MuData [44] and Seurat [7] align multiple assays to a shared cell table, yet running multimodal analyses on them still relies on in-memory processing without a cross-assay memory budget and overwrites per-assay outputs in place without recording which upstream modality settings produced a fused graph. Weighted nearest-neighbor (WNN) integration [7] fits naturally into CyteArc’s architecture because it fuses compact per-modality embeddings and neighbor indices rather than raw counts. CyteArc stores every modality inside one shared store, processes each assay batchwise one at a time under the same memory budget into read-only artifacts, and fuses those stored embeddings and neighbor indices into a WNN graph linked within the same provenance graph (Methods). Across human blood CITE-seq cohorts spanning 0.10 to 1.46 million cells profiled for RNA and 188 shared surface proteins [28, 29], CyteArc completed the two-assay WNN workflow in 1.1 to 13.3 minutes (Fig. 3k), running 41-fold faster than Seurat v5 WNN [6] at 600,000 cells (5.4 minutes versus 3.8 hours). When changing the clustering resolution, CyteArc’s provenance graph automatically reused all upstream RNA, protein, and WNN graph artifacts, reclustering in 8.5 to 20.4 seconds across all sizes, up to 40-fold faster than recomputing from raw counts (20.4 seconds versus 13.6 minutes at 1.46 million cells, with identical cluster labels) and 748-fold faster than Seurat reclustering its stored WNN graph at 600,000 cells (14.7 seconds versus 3.1 hours; Fig. 3l; Supplementary Fig. 3g). Processing assays batchwise kept CyteArc’s peak memory within 11.3 GiB up to 1.46 million cells, whereas Seurat reached 87.6 GiB at 600,000 cells, timed out after 10 hours at 1.0 million cells (mostly in WNN clustering), and ran out of memory during RNA normalization on the full 1.46-million-cell cohort (Fig. 3m). When clustered to match the 19 harmonized author cell types, CyteArc WNN agreed more closely with author annotations than Seurat WNN at every size Seurat completed (mean ARI 0.43 to 0.44 versus 0.35 to 0.36; Fig. 3n) and improved recovery of the ten surface-protein-defined immune states over both Seurat WNN and RNA alone (mean F1 0.44 to 0.45 versus 0.30 to 0.33; Supplementary Fig. 5a). Without batch correction, CyteArc WNN substantially outperformed RNA alone within COMBAT (mean ARI 0.77 to 0.78 versus 0.60 to 0.62), whereas lower surface-protein agreement in Stephenson left pooled WNN (ARI 0.43 to 0.44) slightly below RNA alone (ARI 0.46). Once Harmony2 corrected for study across all 1.46 million cells, CyteArc WNN surpassed both single modalities overall (ARI 0.44 versus 0.41 for RNA and 0.20 for surface proteins; Fig. 3o; Supplementary Note 17).

Most single-cell reference mapping today relies on unimodal RNA atlases, yet the same direct reference reuse used above extends to multimodal CyteArc stores. Because a completed WNN analysis keeps its RNA embedding, RNA neighbor index, and protein-resolved cluster labels linked inside one store, a multimodal atlas store can transfer its sharper cell-state definitions onto new studies that profile RNA alone. Clustering 371,747 T cells from a CITE-seq blood atlas with WNN resolved seven named T-cell subtypes (Fig. 3p). When we projected an independent CITE-seq cohort [7] onto this reference’s stored RNA neighbor index using RNA counts alone (68,085 query T cells carrying matching subtype labels), transferring WNN-defined cluster labels raised subtype agreement from 68.3% (using RNA-only reference clusters) to 83.4%. Transfer accuracy improved in five of the seven subtypes, with the largest gains in memory CD4 T cells (47.4% to 87.5%) and effector CD8 T cells (79.5% to 92.5%). Agreement remained essentially unchanged in mucosal-associated invariant T (MAIT) cells (92.0% to 92.2%) and dipped slightly in regulatory T cells (33.6% to 30.8%), where the resolution-0.5 WNN reference cluster was only half regulatory T cells and both references showed low RNA-only transfer recall (Fig. 3q). Because the query cohort also measured surface proteins that were withheld during mapping, we used those independent protein profiles across all 73,034 RNA-mapped T cells to validate the transferred labels; every transferred subtype reproduced its canonical surface markers, including TCR V*δ* 2 on *γδ* T cells, TCR V*α*7.2 on MAIT cells, CD57 on effector CD8 T cells, and CD25 on regulatory T cells (Fig. 3r; Supplementary Note 18). By keeping multimodal embeddings, neighbor indices, and WNN annotations linked inside a single queryable store, CyteArc lets routine RNA-only studies inherit protein-resolved cell states directly from multimodal atlas stores without exporting standalone reference files or training a separate classifier.

## Discussion

CyteArc turns public single-cell datasets from static download archives into live, shared cloud workspaces where researchers can analyze, combine, and extend atlases directly in cloud storage. Each cloud-hosted store, such as those in CyteBase, pairs lightweight local access with dual-orientation counts processed under a memory budget and a built-in read-only record of every step. Because upstream results stay stored alongside the data, new analyses can build directly on prior work without copying raw counts. Together, CyteArc and CyteBase lower the hardware barrier to atlas-scale genomics, allowing groups without GPUs to analyze nearly 100-million-cell atlases at full single-cell resolution on standard CPU instances.

Working directly on a cloud repository shifts cross-study biology from downloading and rebuilding entire datasets to querying public archives as a shared resource. Researchers can screen gene programs across dozens of hosted studies, pull out rare cell subsets, and combine multi-million-cell cohorts without duplicating raw counts. In CyteBase blood cohorts, that approach resolved rare circulating hematopoietic progenitors across a nine-state differentiation hierarchy and distinguished their severity-linked expansion in acute COVID-19 from chronic conditions. Keeping every analysis step linked to its inputs extends the same reuse model to iterative reanalysis and cell-type annotation. Completed Harmony2-integrated [26, 27] and multimodal CITE-seq stores act directly as mapping references without separate reference exports, dedicated classifiers [34], or GPU-trained latent models [8, 9]. Because each reference retains its embeddings and neighbor indices, researchers can annotate tens of millions of cloud cells in place and project protein-defined immune states onto routine RNA-only studies.

A scalable store prevents workflow fragmentation only when downstream methods run natively on the same data without exporting matrices across incompatible R and Python packages [45, 46]. CyteArc therefore implements core single-cell methods directly from published specifications under permissive licenses atop its memory planner and provenance graph, including deterministic multi-threaded Leiden clustering [25, 47] (Methods). Across preprocessing, integration, chromatin and regulatory inference, gene-set scoring, and trajectory analysis, these implementations closely reproduce established reference tools under matched settings (Table 2), with all intentional methodological departures documented alongside matched benchmarks (Supplementary Notes 19 to 25).

**Table 2.** Numerical agreement with reference implementations and direct computations under matched inputs and settings. Protocols, software versions, and the four tolerances set in advance are detailed in Methods and Supplementary Notes 19 to 25. |Δ|, absolute difference; ARI, adjusted Rand index; AUROC, area under the receiver operating characteristic curve; PCs, principal components; *r*, Pearson correlation; *ρ*, Spearman correlation.

| Comparison | Reference | Data | Result |
|---|---|---|---|
| **Numerical agreement with matched settings** |  |  |  |
| PCA (Gram and Krylov) | Dense float64 SVD | 3,696 cells; 2,000–5,000 genes; 21 PCs | Max angle 0.0143°; relative variance error 2.8 × 10<sup>−8</sup> |
| Native Leiden | igraph; thread repetitions | 18,597 cells; 1, 4, 8, 16 threads | Partitions identical across threads; modularity 0.86294/0.86278 |
| Complete RNA pipeline | Fresh runs without reuse | 18,597 cells; 3 deterministic runs | Sequential arrays identical; ARI 1.00 |
| Batchwise LSI | Dense SVD; first component omitted | 6,194 cells; 25,000 peaks; 30 components | 80 iterations, 100 oversamples; max angle 0.000935° |
| WNN | Seurat 5.5.1; shared candidate pools and bandwidths | 7,299 CITE-seq cells; 6,194 TEA-seq cells | Neighbors identical; Max \|Δ\| 3.0 × 10<sup>−8</sup> |
| Harmony2 | harmonypy 2.0.2; R harmony 2.0.5; shared PCs and states | 6,000 Kang and 6,000 Binvignat PBMCs; 10 seeds per tool | Assignment step max \|Δ\| 4.5 × 10<sup>−7</sup> (harmonypy) and 1.4 × 10<sup>−6</sup> (R), penalty off; full fits with each tool's stopping rule within harmonypy's seed spread on 2 of 2 datasets |
| Symphony query correction | R Symphony 0.1.3; unclipped shared scaling | 3,000 reference and 500 query HLCA cells; 2,000 genes | Max \|Δ\| 8.0 × 10<sup>−8</sup>; largest correction 7.90 |
| GeneScores | Dense interval overlap and TF-IDF aggregation | 400 cells; 240,122 peaks; 256 genes | Exact; 25 nonoverlapping genes retained |
| chromVAR | R chromVAR 1.34.1; shared motif/background peaks | 2,612 cells; 98,316 peaks; 879 motifs | Max \|Δ\|: deviations 1.7 × 10<sup>−14</sup>; z-scores 2.8 × 10<sup>−12</sup> |
| EmptyDrops | DropletUtils 1.32.1; shared model precision | 1,191/17,781 tested barcodes | Cell-call Jaccard 1.000/0.983; ambient Max \|Δ\| 10<sup>−16</sup> |
| Ambient RNA correction | SoupX 1.6.2; matched pool and contamination fraction | 996/11,769 filtered cells | Contamination Max \|Δ\| 4.0 × 10<sup>−14</sup>; corrected-count difference < 1 |
| HTO demultiplexing | Seurat 5.5.1; k-means | 21,522 cells, 8 tags; 8,438 cells, 10 tags | Global classes identical; 1 singlet identity differs |
| AUCell; WAGGR mean | decoupler 2.2.0 | 4,331 cells; 18 gene sets | Max \|Δ\| 3.0 × 10<sup>−8</sup>; 9.2 × 10<sup>−7</sup>, respectively |
| GRNBoost2 | arboreto 0.1.6; scikit-learn 1.9.1 | 2,612 cells; 14,211 genes; 1,249 factors | Consecutive stopping: max \|Δ\| 0.0; *r* = 1 |
| cisTarget recovery | pySCENIC 0.12.1; shared ranks and weights | 18 regulators; 3 seeds; 54 modules | AUC: Max \|Δ\| 8.3 × 10<sup>−17</sup>; curves: Exact |
| **Markers, trajectories and supporting scores** |  |  |  |
| Marker statistics | presto; Scanpy; BPCells | Three datasets of about one million cells | AUROC: Max \|Δ\| 1.1 × 10<sup>−6</sup>; log<sub>10</sub> *P* versus BPCells: Max \|Δ\| 2.9 × 10<sup>−5</sup> |
| Fate probabilities | CellRank 2; matched graphs | 3,696 pancreatic cells; 1,110 LARRY progenitors | 1 − *r*: 2.1 × 10<sup>−9</sup>; 5.3 × 10<sup>−11</sup>, respectively |
| LISI; cell-cycle scores | scib-metrics 0.6.1; Scanpy 1.12.4 | Shared neighbors; shared control genes and gene lists | Max \|Δ\| 1.5 × 10<sup>−14</sup>; Max \|Δ\| 3.9 × 10<sup>−16</sup>, respectively |
| MAGIC imputation | magic-impute 3.0.0; shared graph and diffusion time | 4,331 PBMCs; 200 genes | Max \|Δ\| 6.7 × 10<sup>−6</sup> |
| Population-balance pseudotime | Dense truncated potential (self-check) | 3,546 pancreatic cells; same 29 modes | Spearman *ρ* = 1.00000; max \|Δ\| 3.187 × 10<sup>−7</sup> |

Two practical considerations guide deployment across workloads. First, while CyteArc keeps batchwise count operations within a configured memory budget, downstream graph and embedding stages still hold per-cell state that grows with dataset size, so CyteArc checks memory requirements upfront and halts cleanly before allocating when a stage exceeds the budget. Second, specialized training loaders such as annbatch [48] and scDataset [49] optimize multi-epoch deep-learning workloads through specialized block or shuffled disk loading, whereas CyteArc prioritizes zero-download remote access; when a workflow requires many repeated passes over a sparse multi-store subset, saving a compact local store once avoids redundant network transfers while preserving the same store format.

As single-cell atlases grow beyond what reviewers and collaborators can practically rerun from raw files and scripts [50, 51], and as computational biology adopts autonomous AI agents [52, 53], in-place edits and unlinked output files break the link between published claims and underlying data. Embedding a read-only provenance graph inside each CyteArc store keeps every cell selection, reduction, neighbor graph, cluster label, and transferred annotation traceable to exact counts and parameters. Consortia, reviewers, and automated agents can therefore verify how any published result was derived, retrieve cloud artifacts in under a minute, and launch new analyses across hosted stores in CyteBase without altering shared data or rerunning upstream steps.

## Online Methods

### Dual-orientation store layout and import

A CyteArc store is a Zarr version 3 hierarchy [54] containing a shared cell metadata table, per-assay feature metadata tables, count arrays, precomputed count summaries and immutable analysis artifacts (Fig. 2a). Count matrices are stored as dense two-dimensional arrays whose inner chunks are packed into shards with the Zarr sharding codec [55] and compressed at import with Blosc LZ4 (level 5, bit shuffle) when targeting local storage or Zstandard (level 3) when targeting cloud object storage (stores imported locally and later uploaded keep Blosc LZ4), read through a Hugging Face fsspec store for CyteBase (hf://) URIs and through obstore [56] for other remote object stores. Every chunk in the declared grid, including all-zero chunks, is written at import so that a missing shard object raises a read error instead of silently returning zeros (an inner chunk missing inside a stored shard still reads as zeros). Throughout this work, MB and GB denote decimal bytes (10^6^ and 10^9^ bytes, layout targets included), whereas MiB and GiB denote binary bytes (2^20^ and 2^30^ bytes).

Cell-wise operations (quality control, PCA and LSI) process sequential batches of complete cell rows, whereas gene-wise operations (variable-gene selection, subset-renormalized log-normalization and marker discovery) process feature bands batchwise across all cells; storing both matrix orientations eliminates unused-feature decoding during gene-wise passes. Store geometry is governed by two import targets: an uncompressed read-unit target *U* (default 1 GB), which bounds one sequential cell-major shard or all-cell feature band, and an inner-chunk target *C* ≤ *U* (default 100 MB). For an assay of *N* cells and *G* features stored at *b* bytes per entry, the cell-major array (counts, *N* × *G*) is partitioned into shards of *h* complete cell rows, each subdivided into inner chunks of *h* rows by 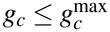 features:

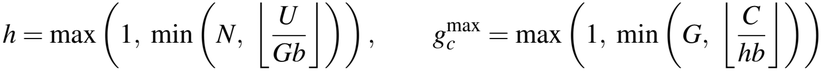

while for RNA assays a paired gene-major copy (countsT, *G* × *N*) processes feature bands of height *w* ≤ *w*^max^ = max(1, min(*G*, ⌊*U/*(*Nb*)⌋)) across all *N* cells without decoding cell-major chunks, where (*g*_*c*_, *w*) are chosen jointly within 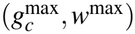 so that one divides the other. Each countsT shard spans *s*_*g*_ = max(*g*_*c*_, *w*) features by *s*_*c*_ = *h* · min(⌈*N/h*⌉, max(1, ⌊*U/*(*hs*_*g*_ *b*)⌋)) cells, a multiple of *h* bounded by *U*, and each inner countsT chunk spans *w* features by *c* cells, where *c* is the largest divisor of *s*_*c*_ not exceeding *c*^max^ = min(*s*_*c*_, max(1, ⌊*C/*(*wb*)⌋)); when that divisor is below *c*^max^*/*2, CyteArc instead aligns *c* and *h* so that one is a multiple of the other, trimming *h* if needed, and recomputes *s*_*c*_. A full-transcriptome gene-major pass issues *n*_chunks_ = ⌈*G/w*⌉⌈*N/c*⌉ chunk reads. When *N* grows large enough that *w < g*_*c*_, the shard height locks to *s*_*g*_ = *g*_*c*_ ≈ *CG/U*, so *s*_*c*_ and *c* saturate near *U/*(*g*_*c*_*b*) ≈ *U* ^2^*/*(*CGb*) cells regardless of *N*, while the band height *w* ≈ *U/*(*Nb*) narrows inversely with *N*. Under fixed (*U,C*), *n*_chunks_ therefore grows as *N*^2^. The targets are set at import and stay at their defaults unless the user changes them; raising *U* (as in the 5-million-cell scaled layout, *U* = 2 GB, *C* = 200 MB) or raising *U* relative to *C* to shrink *g*_*c*_ (as in the 10-million-cell scaled layout, *U* = 2 GB, *C* = 100 MB) widens *s*_*c*_ and *c*, keeping request counts linear in *N* across the 5- and 10-million-cell embryo stores.

At import, a matrix whose values are all non-negative integers, regardless of its source dtype, is stored in the narrowest unsigned integer dtype (uint8 to uint64) that holds its largest count; matrices with non-integral or negative values keep their source dtype, with float16 widened to float32. The importer processes input rows batchwise once to write counts while computing per-row nonzero-entry digests, cell totals, detected-feature counts and per-feature detected-cell counts, then builds countsT by destination shard, several shards at a time: each shard’s buffer is reserved in the memory ledger, filled by reading each overlapping counts chunk once and committed in a single write. Before writing any data, the importer verifies that the planned buffers fit within the memory budget and otherwise stops, reporting the largest pair of halved targets (*U,C*) that fits.

### Memory-budgeted batchwise execution and admission control

Each datastore resolves an operation memory budget 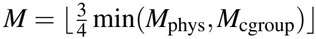, equal by default to three quarters of the smaller of the host’s physical memory *M*_phys_ and any active Linux cgroup limit *M*_cgroup_, which users may override in bytes or as a fraction in (0, 1); a compute-worker limit *W* defaults to the smallest of the logical CPU count, the process’s CPU affinity and the cgroup CPU quota. *M* governs the buffers each operation allocates to read, decode, compute and write; it is not an operating-system cap on process memory, and memory that caller code holds outside the operation lies outside *M*.

For a batchwise pass over *N*_*u*_ work units, an operation that retains *R* resident bytes and *S* scratch bytes throughout leaves *B* = *M* − *R* − *S* bytes for units in flight. Each unit in flight needs *u* bytes for decoded arrays, compressed payloads and outputs, *d* bytes of codec workspace and *m*_in_ ≥ 1 concurrent inner chunk reads of *r*_in_ bytes each, with *m*_in_ capped by the inner chunks per shard (or an explicit per-unit read cap). Keeping a floor on *L* at the active compute or write width (bounded by ⌊*B/*(*u* + *d* + *r*_in_)⌋), the execution planner chooses the number of units in flight *L* together with *m*_in_, maximizing *Lm*_in_ subject to

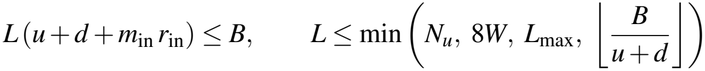

so that roughly eight reads per worker overlap with multi-threaded computation (read-group passes such as marker search can hold slightly more, up to (*W* + 1)⌈8*W/*(*W* + 1)⌉, or 136 at *W* = 16). The in-flight unit cap *L*_max_ is *W* + 1 for unordered all-cell read-group passes, such as marker search, 8*W* for gene-major cell-band passes, such as variable-gene statistics and normalization, and otherwise follows the pass’s ordering or prefetch depth. If *B* ≤ 0, or a single unit (including one inner read when *r*_in_ *>* 0, so *u* + *d* + *r*_in_ *> B*) exceeds *B*, the operation raises a MemoryError before reading data.

Asynchronous gene-major passes (countsT construction, variable-gene statistics, subset-renormalized normalization and marker discovery) and virtual composite reads also enforce *B* at run time through a reservation ledger. Each buffer request waits until enough budget is free, fails at once if it exceeds *B*, and bytes still held when the pass completes raise an error. Cell-major batchwise passes (import, quality-control summaries, PCA and LSI) size their block heights and prefetch queues with the same planner. Steps with bounded feature-space state (exact Gram or randomized Krylov PCA and batchwise LSI) and steps whose state grows as *O*(*N*) with the number of cells (k-means embedding initialization, Harmony2, HNSW index construction and search, connectivity graph construction, multimodal graph fusion, Leiden clustering and UMAP) estimate their peak memory from the shapes of their inputs and, when the estimate exceeds *M*, stop before allocating and report the memory required. BLAS, OpenMP and Numba thread pools are limited to max(1, ⌊*W/W*_active_⌋) threads per active worker, where *W*_active_ is the operation’s number of concurrent compute or write workers, and each planner-managed count pass logs its planned and observed worker widths, its I/O and compute waits and its peak held bytes.

### Immutable artifacts and provenance

Every computed result (cell or feature selections, normalized matrices, feature scalings, reductions, batch corrections, neighbor indices, graphs, embeddings, clusterings, marker tables, projections and label transfers) is persisted inside the store as an immutable Zarr group with a random 256-bit hexadecimal identifier (`artifact_id`) and a canonical provenance record *π*(*a*) = (*o, θ, I, r*) comprising the operation name *o*, scientific parameters *θ*, input artifacts *I* and operation revision *r*, a positive integer omitted when *r* = 1. The inputs link all outputs into a directed acyclic graph rooted in the assay’s count matrix (Fig. 3a). Scientific parameters include random seeds and the execution flags that change floating-point summation or graph construction order, such as `ann_parallel` and parallel UMAP optimization, whereas thread counts, block sizes that do not change results, creation times and package versions are recorded separately, in `execution_options` and group attributes, so that a change of hardware never invalidates reuse.

Scientific inputs that are not JSON values enter *π*(*a*) as BLAKE2b digests: arrays, such as boolean cell selections, as 256-bit digests of their canonical little-endian dtype, shape and bytes; user-supplied Python functions by a declared identity or, failing that, by their module and qualified name with a 128-bit digest of their compiled bytecode, constants, variable names and defaults; and the root count matrix as a layout-independent dataset fingerprint of its shape, dtype and per-row nonzero-entry digests. Provenance records are serialized canonically to UTF-8 JSON with sorted keys, explicit type tags and big-endian IEEE-754 hexadecimal strings for finite float64 scalars. When an operation is requested, CyteArc compares the canonical bytes of (*o, θ, I*) directly with those of the complete artifacts in scope, with no digest serving as a lookup key, and reuses the newest complete artifact whose bytes match at the current revision *r*.

An artifact’s complete attribute is set to true in a single write, only after its payload is written and validated: the required arrays are present with their expected shapes and dtypes, the required attributes are set and any operation-specific validator passes, so an interrupted write is never reused. When a CyteArc release changes an algorithm’s numerical output, incrementing that operation’s revision *r* retires the superseded artifacts from automatic reuse while keeping them readable for audit and reproduction. Pipeline runs record whether each stage was reused or computed, its wall-clock duration and the process tree’s resident memory, sampled every 0.1 s.

### Overlay mounts and virtual composites

Mounting a remote store (an overlay mount, `mount_datastore`) creates a writable user store, on local disk or in object storage, that copies only the source’s cell and feature metadata tables and records the source URI with its dataset and count fingerprints (Fig. 1a). Count chunks are streamed on demand from the read-only source in either orientation, and artifact resolution queries the union of the source and user stores, so matching reductions, graphs or clusterings already in the remote store are reused in place. All new artifacts and metadata columns are written to the user store, and writes to the source raise an error. Each reopening compares the dataset and count fingerprints stored with the source against those recorded at mount time, and cloud credentials are supplied when a session opens and never written to the store.

A virtual composite store (`create_composite`), created on local disk, assembles selected cells of several read-only stores of one assay type into one writable store without copying count chunks (Fig. 1a). Features are matched by exact identifier, through intersection or union, and features a source does not measure read as zero. To mix studies across cell batches while keeping each block read within one source shard, each source’s selected cells are split at its cell-major shard boundaries and permuted within each block, and the non-final blocks of all sources are shuffled together before each source’s final block is appended (seed 42); each cell identifier is prefixed with its source’s name and checked for uniqueness. Both counts and countsT are virtual arrays: for each two-dimensional read, each source is read in whichever stored orientation needs fewer estimated decoded bytes, and each touched source chunk is decoded at most once per batch. Encoded shards of remote sources are kept in an optional local disk cache (2 GiB and 4,096 entries by default; 0 turns it off). At creation, the selected counts are read batchwise once to compute cell quality-control totals and the composite’s own root count fingerprint, which roots the provenance of its artifacts, and reopening stops if any source fingerprint has changed. Composites can also be exported batchwise to H5AD on disk.

### CyteBase resource construction

CyteBase release 2610 was built from a snapshot of CZ CELLxGENE Discover [2] taken on 10 October 2026. From 2,170 datasets in 376 registered collections, we excluded 792 datasets whose observations are all secondary (`is_primary_data` false, which CELLxGENE assigns to cells that are not the canonical instance of an observation, such as cells reused in meta-analyses or secondary views, and to all cells of spatial datasets with more than one section) and 6 non-RNA datasets, yielding 1,372 primary-count datasets; 29 datasets in 11 further collections were held back until their assays are reviewed. Each source H5AD was imported into a cloud-profile CyteArc store (Zstandard level 3, with counts and countsT) that keeps the authors’ cell annotations and embeddings beside the CELLxGENE collection and dataset identifiers, the SHA-256 checksum of the source H5AD and a build receipt of checks on the imported counts.

A DuckDB catalog, published with its SHA-256 checksum, indexes all 1,372 stores by organism, tissue, organ, assay, disease, cell type, sex, developmental stage and suspension type, and before opening or mounting a store, the CyteBase client reads its own `dataset.json` record to verify its CELLxGENE dataset version and source SHA-256 checksum, then confirms the opened cell, feature and count-array shapes against the committed build receipt. Each release is a Hugging Face bucket named by year and month, such as Nygen/cytebase-2610. Storage totals come from listings of the bucket’s objects on 14 October 2026, with a store’s nonzero count entries estimated as its cells times its mean detected genes per cell, and disease coverage counts human non-spatial stores with at least 10,000 primary cells.

### Single-cell RNA-seq analysis workflow

When a store is first opened for writing, cells with at most 10 detected features are masked unless that would remove at least half of the active cells, which protects raw unfiltered droplet matrices from silent truncation. CyteArc’s default quality-control filter, pooled or per sample, keeps cells within *m* ±3×1.4826 MAD on the ln(1 + *x*) scale for total counts and detected genes, and below *m* + 3 ×1.4826 MAD of the raw mitochondrial (MT-) and ribosomal (RPS, RPL, MRPS and MRPL) percentages, where *m* is the median and MAD the median absolute deviation. The toolkit, embryo scaling and Tahoe-100M benchmarks used the Gaussian filter, which keeps cells strictly between the 1st and 99th percentiles, 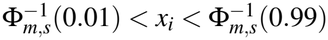,of a normal distribution with each metric’s median *m* and population standard deviation *s* on its raw scale; the Kang reference used the default MAD filter, while the cross-study mapping, reference-ladder and multi-study composite runs used the curated source cells without additional cell filtering.

Highly variable gene (HVG) selection (flavor cytearc; `seurat_v3` and `pearson_residuals` flavors are also available) processes countsT batchwise over the *n* retained cells with full-transcriptome totals *t*_*i*_ (a zero total counted as 1) to compute library-scaled values *y*_*ig*_, population means *µ*_*g*_ and population variances 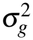. Across every gene with *µ*_*g*_ *>* 0, a baseline 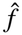 of log variance against ln *µ*_*g*_ is fitted by local 0.25-quantile regressions of degree at most 2, anchored at the median ln *µ*_*g*_ of each of 200 equal-width bins of a coordinate that averages each gene’s normalized rank and normalized ln *µ*_*g*_, and interpolated linearly between anchors. Ranking then keeps the *G*_HVG_ candidates with the largest variance ratios *v*_*g*_ (1,000 by default) among genes detected in at least 20 retained cells (scaled down proportionally to max(3, ⌈0.01*n*⌉) and capped at ⌊*n/*5⌋ for datasets below 2,000 cells) and in at most 99.5% of them, excluding case-insensitive matches to mitochondrial, ribosomal, cyclin (CCN[A-Z]), HLA-, H2-, replication-dependent histone and ten sex-linked genes:

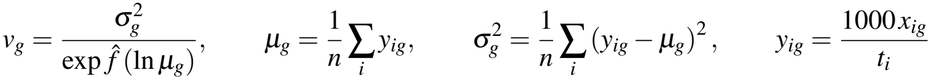

By default, expression over the selected genes *S* is normalized in float64 with each cell’s subset total 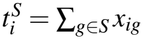 (a zero total counted as 1), stored in float32 and standardized per gene to zero mean and unit population standard deviation without clipping:

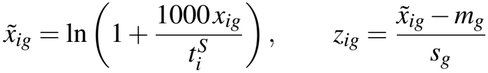

where *m*_*g*_ and *s*_*g*_ are gene *g*’s mean and population standard deviation, with *s*_*g*_ = 1 when it is zero. For up to 4,096 genes, PCA accumulates the exact float64 sample covariance 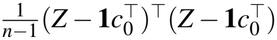 of the scaled values *Z* in one batchwise pass, where *c*_0_ is their mean over the cells that fit the PCA, and takes the leading *d* eigenvectors by symmetric eigendecomposition (*d* = 21 by default). For more genes, CyteArc uses a batchwise randomized block-Krylov solver [57, 58] with block width *d* + max(10, *d*), which stops when the largest principal angle between the leading *d*-dimensional subspaces of consecutive passes is at most 0.1°, after at most 24 passes.

Approximate nearest-neighbor search indexes the reduced coordinates in an hnswlib HNSW graph [59] under Euclidean distance and retrieves each cell’s *k* nearest non-self neighbors (*k* = 11 by default). By default, the index uses *M* = 24 (up to 48 links per node in the base layer), a construction depth of 200 and a query depth of max(200, 2*k*), built serially for deterministic reproducibility or in parallel when requested (benchmark parameter configurations in Supplementary Note 9). Directed fuzzy membership strengths [24] over the neighbors *j ∈ N*_*k*_(*i*) are computed with umap-learn’s kernels:

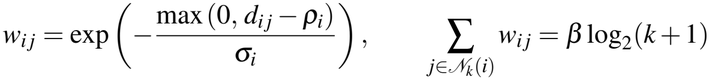

where *ρ*_*i*_ is the distance from cell *i* to its nearest neighbor at positive distance, *β* = 1.5 is the default bandwidth multiplier and *σ*_*i*_ is found by umap-learn’s bisection search of up to 64 iterations. To initialize two-dimensional embeddings deterministically in bounded memory, CyteArc fits 1,000 mini-batch k-means centroids on the graph’s input coordinates (PCA coordinates, or Harmony2-corrected coordinates in integrated runs; seed 4466), projects the centroids onto their two principal axes, clips each axis to the 10th and 90th percentiles of a normal distribution fitted to its median and population standard deviation, and places each cell at its centroid’s position; UMAP then optimizes the fuzzy union *W* + *W* ^*⊤*^ −*W* ° *W* ^*⊤*^ with umap-learn’s optimizer for 300 epochs (spread 2.0, minimum distance 1.0, five negative samples per edge, seed 4444).

Community detection uses CyteArc’s native multi-threaded Leiden implementation [25, 47], which maximizes the configuration-model modularity *Q* at resolution *γ* (default 1.0, seed 4444) on the symmetrized adjacency *A* = *W* + *W* ^*⊤*^ in three outer iterations of parallel local moving, community refinement and graph aggregation:

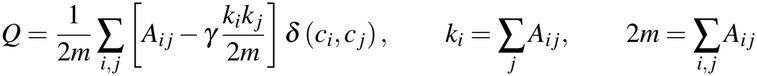

where *c*_*i*_ is the community of cell *i*. Counter-based random streams, snapshot-isolated parallel moves, exact integer community totals and fixed-order block reductions are designed to give identical cluster assignments on any number of CPU threads; graph levels with fewer than 2^21^ stored entries move nodes on one thread, so the 1- to 16-thread check on 18,597 Kang PBMCs tested parallel refinement but not parallel moves. One-versus-rest marker discovery processes countsT batchwise gene by gene across the *n* = *n*_1_ + *n*_2_ cells of a cluster (*n*_1_) and the rest (*n*_2_), ranking float32-rounded library-normalized values 1000 *x*_*ig*_*/t*_*i*_ with mid-ranks by a zero-aware three-pass radix sort of the nonzero entries. From the cluster’s rank sum *R*_1_, CyteArc computes the Mann-Whitney statistic *U*_1_ = *R*_1_ − *n*_1_(*n*_1_ + 1)*/*2, the area under the receiver operating characteristic curve AUROC = *U*_1_*/*(*n*_1_*n*_2_), and two-sided P values, evaluated exactly by dynamic programming over doubled mid-ranks when 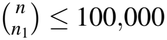 and otherwise from the standard continuity- and tie-corrected normal approximation *P* = 2Φ(−|*z*|) using the tie-adjusted variance 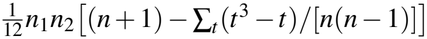 over tied-value group sizes *t*, followed by Benjamini-Hochberg adjustment within each cluster [60].

### Harmony2 batch integration and Symphony reference mapping

Batch correction uses CyteArc’s implementation of Harmony2 [26, 27], with at most 40 outer rounds of at most 200 clustering iterations each (seed 4466) over *K* = min(100, max(1, round(*n/*30))) clusters. Each round alternates soft clustering of the current corrected coordinates with a mixture-of-experts ridge correction applied to the original PCA coordinates. The clustering is maximum-diversity soft spherical k-means of the unit-normalized corrected coordinates 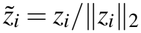 against unit centroids 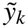 (width *σ*_*k*_ = 0.1, diversity penalty *θ*_*v*_ = 2 for each batch variable *v*), which updates the assignments of blocks of 5% of the cells, taken in one random order per fit, by

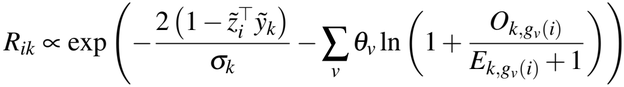

where *g*_*v*_(*i*) is cell *i*’s level of batch variable *v, O*_*kg*_ is the observed assignment mass of batch level *g* in cluster *k*, and *E*_*kg*_ = (*N*_*g*_*/n*) *n*_*k*_ is the independence-expected mass from the full-dataset batch proportion *N*_*g*_*/n* and cluster mass *n*_*k*_, with *O*_*kg*_ and *n*_*k*_ computed on the cells outside the update block. In the correction, using *O*_*kg*_ and *E*_*kg*_ over all *n* cells, the batch levels *g* with *O*_*kg*_ *>* 0 and *O*_*kg*_ ≥ 10^−5^*N*_*g*_ form the active set *A*_*k*_ of cluster *k*, and the coefficients for the original PCA coordinates *Z*_*i*_ minimize

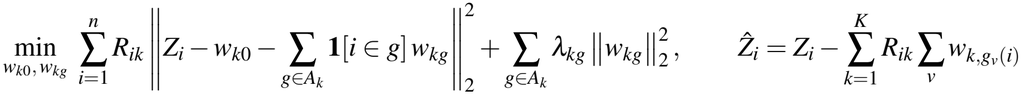

with ridge penalty *λ*_*kg*_ = 0.2 *E*_*kg*_ and *w*_*kg*_ = 0 for *g* ∉ *A*_*k*_, solved in closed form by arrowhead elimination for one batch variable or by Schur-complement Cholesky factorization for several. The centroids, cluster masses and widths are stored for query correction.

Registering a completed CyteArc analysis as a mapping reference writes a compact descriptor without reading reference count arrays. The descriptor stores the selected feature identifiers with their scaling means *m* and standard deviations *s*, the PCA center *c*_0_ and loadings *P*, 1,001 quantiles of up to 100,000 reference first-neighbor distances and, for integrated references, the Harmony2 centroids, corrected centroids, cluster masses 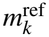 and widths *σ*_*k*_, while referencing the stored reduction and HNSW index artifacts in place. Query cells are library-normalized with the reference’s normalization settings using each query cell’s full-assay count total (or the shared-feature total when `renormalize_subset` is enabled), unmeasured reference features taking the reference mean, and projected into reference PCA space as *z*_*i*_ = *P*^*⊤*^((*x*_*i*_ − *m*) *⊘ s* − *c*_0_), where *⊘* denotes elementwise division; query cells with zero raw counts across all measured reference features are flagged as uninformative. For integrated references, Symphony query correction [35] assigns the unit-normalized projected query vectors 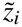 softly to the unit Harmony2 centroids 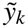 and subtracts per-cluster offsets *O*_*bk*_ of the cell’s query batch *b*(*i*):

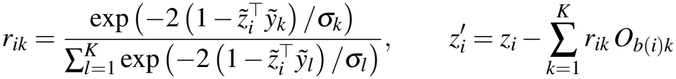

where the offsets are estimated per cluster by ridge regression with unit penalty on the batch terms, anchored to the Harmony2-corrected reference centroid with prior mass 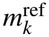; the stored reference HNSW index is then queried at 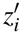.

Label transfer freezes the chosen reference label column or cluster artifact into the query store and scores each informative query cell over its *k* nearest reference neighbors (`save_k`, 3 by default). Each class *ℓ* receives vote 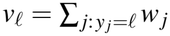 from inverse-distance neighbor weights 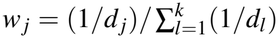 normalized over all *k* neighbors (including unlabeled reference cells, which add to the denominator without voting for any class), or equal weights 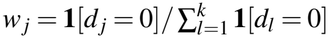 shared across zero-distance neighbors when min_*l*_ *d*_*l*_ = 0. A query cell is left unassigned, with a typed abstention code, if it is uninformative, has no labeled neighbors, ties for the top vote *v*_*ℓ*_ within `np.isclose` tolerance (`rtol` 10^−5^, `atol` 10^−8^), has a top vote below `threshold_fraction` (0.5 by default; 0, a plurality vote, in the mapping benchmarks, the in-place annotation, and the T-cell subtype transfer) or has its nearest reference neighbor beyond an optional `max_distance`.

### Multimodal assay normalization and joint graph fusion

A single cell table links RNA, antibody-derived tags (ADT), sample hashtags (HTO) and chromatin accessibility (ATAC) within one store, so each modality executes batchwise under the shared memory planner into assay-scoped artifacts. CyteArc normalizes ADT counts by a feature-wise centered log ratio (CLR) across the *n* selected cells [41], 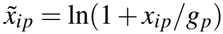 with geometric mean 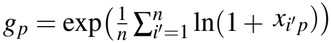, and also accepts a user-supplied function; the COMBAT T-cell subtype reference used a cell-wise CLR over the non-control proteins, supplied this way by the experiment scripts (`cell_clr`). ATAC peak counts *x* are normalized by term frequency-inverse document frequency (TF-IDF), 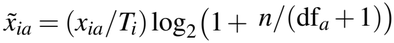, where *T*_*i*_ is cell *i*’s total count across peaks and df_*a*_ the number of selected cells with *x*_*ia*_ *>* 0, followed by batchwise randomized latent semantic indexing (LSI) that omits the depth-correlated first component [61] (five power iterations and ten oversamples by default).

Weighted nearest-neighbor (WNN) integration [7] fuses two or more modalities from their stored neighbor indices and L2-normalized reductions 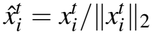, restricting each cell’s candidates 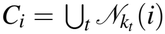 to the union of its stored modality neighbors, where Seurat searches a wider candidate set by default. Within modality *t*, the exponential kernel *θ*_*t*_(*i, δ*) = exp −max(*δ* − *ρ*_*t*_(*i*), 0)*/σ*_*t*_(*i*)) uses the distance *ρ*_*t*_(*i*) to the cell’s nearest neighbor and the bandwidth 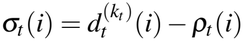, the distance to its *k*_*t*_-th neighbor less *ρ*_*t*_(*i*); Seurat’s default bandwidth is computed differently (Supplementary Note 19). With 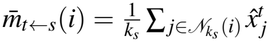 denoting the mean in modality *t* of the cell’s modality-*s* neighbors, the cross-modality prediction ratios *S*_*ts*_(*i*) and softmax modality weights *w*_*t*_(*i*) are

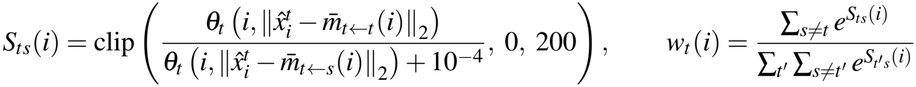

and the *k* = min_*t*_ *k*_*t*_ candidates *j ∈ C*_*i*_ with the largest joint affinity 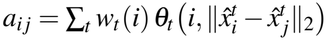 form the directed WNN graph. Shared nearest-neighbor (SNN) fusion, which needs only the connectivity graphs, instead ranks the edges of each modality graph *g* by *r*_*g*_(*i, j*) = *w*_*g*_(*i, j*) + |*N*_*g*_(*i*) ∩ *N*_*g*_(*j*)|*/*(*k* − 1), where *k* − 1 is the largest possible overlap of two self-free *k*-neighbor sets, and keeps the *k* highest-ranked distinct neighbors of each cell with their original weights *w*_*g*_(*i, j*).

### Doublet detection and matched-input method verification

Doublet scoring adapts Scrublet [62] to reuse the stored uncorrected PCA model instead of fitting a second reduction. Within each sample of *n*_*s*_ cells, *s* = max(1, round(*rn*_*s*_)) simulated doublets (*r* = 2 by default) are made by summing the raw counts of random cell pairs, normalized with the summed parent totals and the reference’s variable-gene settings, and projected onto the stored PCA loadings beside the observed cells. A joint HNSW index (seed 4444) over the *n*_*s*_ + *s* points, queried for *K* = round(*k*(1 + *s/n*_*s*_)) neighbors (*k* = min(30, *n*_*s*_ − 1) by default), gives each point *d*_*i*_ simulated neighbors and the Laplace-smoothed simulated fraction *q*_*i*_ = (*d*_*i*_ + 1)*/*(*K* + 2). Reading *q*_*i*_*/*((1 − *q*_*i*_) *s/n*_*s*_) as the local ratio of doublet to singlet density, with a prior doublet rate *ρ* = 0.05, gives the Bayes posterior score *p*_*i*_ = [*q*_*i*_ *ρ/*(*s/n*_*s*_)]*/ q*_*i*_ *ρ/*(*s/n*_*s*_) + (1 − *q*_*i*_)(1 − *ρ*), and cells are called doublets when *p*_*i*_ exceeds the minimum between the two modes of a 256-bin histogram of the simulated doublets’ scores, smoothed by a three-bin running mean until two maxima remain (at most 10,000 steps), or Otsu’s threshold when the histogram does not become bimodal.

Table 2 evaluates numerical agreement under matched inputs and convergence settings, against reference implementations or direct computations and, where one was set, a predefined tolerance: Gram and block-Krylov PCA against a dense float64 SVD on 3,696 mouse pancreas cells [63] (tolerance, largest principal angle 0.05°); native Leiden at 1 to 16 threads against igraph on 18,597 Kang PBMCs; three fresh runs of the RNA workflow with artifact reuse turned off; batchwise LSI (80 power iterations, 100 oversamples) against a dense SVD on 6,194 TEA-seq PBMCs (0.05°); the WNN kernel on shared candidate pools and bandwidths against Seurat (10^−6^); Symphony query correction under shared, unclipped scaling against R Symphony 0.1.3 (10^−5^); and matched-input checks of EmptyDrops, SoupX, HTODemux, GeneScores, chromVAR, GRNBoost2, cisTarget, AUCell, WAGGR, cell-cycle scoring, LISI, MAGIC imputation, Population Balance Analysis and CellRank absorption probabilities [43, 64–71] (Supplementary Notes 19 to 25).

### Software, benchmark execution and scaling metrics

This paper describes CyteArc 1.3.0, available on PyPI, and the runs that used this release ran it under Python 3.14.4 with CyteBase 0.1.0 (Python 3.14.2 inside the R images). External comparators were Scanpy 1.12.4 with anndata 0.13.4 and Dask 2026.8.0, CELLxGENE Census 1.18.0, CellTypist 1.7.1, and R 4.6.1 with Seurat 5.5.1, BPCells 0.3.1, uwot 0.2.5 and Symphony 0.1.3 (the Seurat WNN golden fixtures were generated under R 4.4.3).

Each benchmark job ran in its own Linux container. Input preparation (building a CyteArc store, writing BPCells on-disk matrices or staging uncompressed H5AD inputs for R) was timed separately from analysis, which runs from reopening the prepared input to the end of marker testing and excludes the initial dataset download and container start-up; multi-job Tahoe-100M runs sum their stage times. Across the 1- to 10-million-cell toolkit benchmarks, embryo scaling series and Tahoe-100M runs, workflows targeted *G*_HVG_ = 1,000 variable genes, *d* = 21 principal components, *k* = 11 neighbors, 300 UMAP epochs and Leiden resolution *γ* = 1.0 (using parallel HNSW construction with the default *M* = 24 and depth 200 in the embryo scaling series, storage-backend comparisons and Tahoe-100M runs, and *M* = 48 with construction and query depths of 50 in the toolkit comparisons), and the Kang perturbation mapping likewise used the defaults (*G*_HVG_ = 1,000, *d* = 21, *k* = 11, serial HNSW, and `save_k` of 11), whereas the multi-study composite, tissue and corpus reference-mapping and multimodal RNA analyses used *G*_HVG_ = 2,000, *d* = 30, serial HNSW construction, and `save_k` of 30 reference neighbors for blood, kidney and lung mapping (15, with a 0.6 vote threshold, in the lung reference revision) or 20 for CITE-seq T-cell subtype transfer (full comparator configurations in Supplementary Note 9). Memory was sampled every 0.05 to 1 s, depending on the experiment, from both the container’s cgroup counter (memory.current, or memory.usage_in_bytes under cgroup version 1, which includes page cache and runtime overhead) and the process tree’s resident memory, and a reported peak is the largest sample over the measured span. Remote query benchmarks record, for CyteBase, both the bytes returned to Zarr and the encoded HTTP response-body bytes counted at the httpx2 client transport, and for CELLxGENE Census all bytes received on the container’s network interfaces (`/proc/net/dev`, loopback excluded), which include protocol overhead.

Toolkit runtime comparisons evaluate each method in the smallest container tier in which it completed. Overall speedups compare the geometric-mean runtimes of CyteArc (*A*) and a comparator (*B*) as 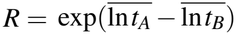, reported as a time reduction 1 − *R* or a fold speedup 1*/R*; per-stage speedups are ratios of arithmetic means. Analysis core-time multiplies analysis runtime by the container’s allocated CPUs (16 times the summed stage times for Tahoe-100M), and the scaling exponent *b* is the ordinary least-squares slope of log_10_ *T*_*m*_ = *a* + *b* log_10_ *N*_*m*_, where *T*_*m*_ is the core-time and *N*_*m*_ the cell count of run *m*, over 18 runs (three replicates at each of five mouse embryo sizes from 100,000 to 10,000,000 cells and three Tahoe-100M replicates), alongside the exponent fitted over the 12 runs from 1,000,000 to 95,624,334 cells. Gene-pass layout scaling exponents between 5,000,000 and 10,000,000 cells are 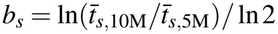, from three-run arithmetic means. Decode amplification *α*, the ratio of uncompressed bytes decoded to bytes used when an aligned gene band is read from the cell-major counts array alone, is *α* = max(1, *g*_*c*_*/w*), against *α* = 1 from countsT.

### Integration, reference-mapping and clustering metrics

In multi-study composites, the within-cell-type study batch entropy and the kNN cell-type purity use each cell’s 30 stored neighbors. Let *T* be the harmonized cell types with at least 20 cells in every study and, for *c ∈ T*, let *E*_*c*_ be the cells of type *c* with at least five same-type neighbors, *M*_*i*_ denoting cell *i*’s same-type neighbors. With *p*_*is*_ the fraction of *M*_*i*_ from study *s, H*_*i*_ = −∑_*s*_ *p*_*is*_ ln *p*_*is*_ its Shannon study entropy, and *H*_*c*_ the Shannon study entropy of all cells of type *c*, normalized batch mixing is 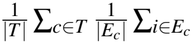 min(*H*_*i*_*/H*_*c*_, 1), and kNN cell-type purity is 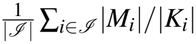, where *K*_*i*_ is the set of cell *i*’s annotated neighbors and *ℐ* the annotated cells with at least one annotated neighbor. In the CITE-seq series, purity counts each cell’s 19 nearest neighbors, unannotated ones as mismatches, and the purity of a Tahoe-100M cluster is the share of its cells from its most common cell line.

Reference-mapping evaluations tabulate a confusion matrix *C* whose rows *ℓ ∈ ℒ* are the harmonized author labels present in the query and whose columns *ℓ*^′^ *∈ ℒ* ^′^*∪*{*∅*} are every label of the reference or the query (*ℒ* ^′^ *⊇ ℒ*) and unassigned cells, so that abstentions and reference-only predictions count as errors in overall accuracy 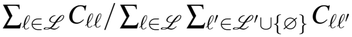 (with disagreement 1 −accuracy) and per-class 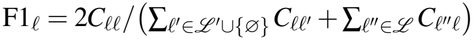. Macro-F1 averages F1_*ℓ*_ equally over the labels present in the query, and rare-type recall averages 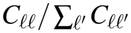 over types below 1% of the reference, with at least 20 reference cells in the held-out tissue cohorts; in-place annotation of the blood corpus evaluates the cells whose author label maps to one of the nine reference classes. Partition concordance uses the adjusted Rand index (ARI) and normalized mutual information (NMI) of scikit-learn [72] over annotated cells. In the CITE-seq scaling series, every graph was clustered with CyteArc’s Leiden at the resolution, found by bisection, that gave a target cluster count with seed 4466: 19 clusters, which seeds 1 and 2 reran at the same resolution (16 to 22 clusters), and the cluster counts each tool recorded, such as Seurat’s 105 and 112; cells in connected components of fewer than 10 cells joined their nearest neighbor’s cluster. Protein-defined subtype recovery gives each cluster its majority author label and averages F1_*ℓ*_ over the ten types the authors defined by surface protein.

### Corpus-wide interferon scoring and circulating progenitor meta-analysis

Before donor-level scoring in both the blood interferon screen and the circulating progenitor models, uniform blood sample rules defined from each store’s cell metadata exclude 11 sorted, enriched, stimulated, or cell-product stores whole, drop sorted, enriched, or stimulated sample subsets within mixed stores using store-native labels, and retain one sample per donor (the earliest by the store’s recorded time columns, or the largest when no time is recorded; stores without a sample column use their first recorded time column to define samples when present, and otherwise treat all of a donor’s cells as one unit). For the CyteBase blood interferon screen, each scoring unit *u* is one study-scoped donor’s retained sample within one coarse immune lineage and store, with at least 20 cells. Summing the raw counts *x*_*ug*_ of each measured gene *g* of the 18-gene type I interferon panel and the full-transcriptome counts *t*_*u*_ over the unit’s cells gives the pseudobulk log-CPM expression *v*_*ug*_ = log_2_(1 + 10^6^ *x*_*ug*_*/t*_*u*_), which is standardized against the store’s healthy-control units *R* of the same lineage (from at least three healthy donors, with sample mean 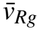 and sample standard deviation *s*_*Rg*_ on |*R*| − 1 degrees of freedom) as 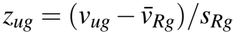 and averaged into score 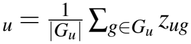 over the measured panel genes *G*_*u*_ with *s*_*Rg*_ *>* 0. Within each study and lineage with at least five disease and five healthy donors, disease elevation is quantified by the AUROC of donor scores, computed from mid-ranks as for markers; acute viral infections (COVID-19 and influenza) and type-I-interferon-associated autoimmune diseases (systemic lupus erythematosus and juvenile dermatomyositis) formed the prespecified interferon-driven group, and all other qualifying diseases served as non-interferon controls. Before the contrasts, the 23 donors of the COMBAT store’s sepsis cohort, which the store labels COVID-19, were relabeled as sepsis, two COVID-19 low-density-neutrophil samples were dropped, and the colorectal cancer atlas was split by its source studies.

For the circulating hematopoietic progenitor models, after applying the blood sample rules described above and excluding two leukemia cohorts (Chen 2022 infant B-ALL and Lim 2025 childhood T-ALL) and one 460-gene panel store (Triana 2021), each study-scoped donor *d* is represented by its earliest (or largest undated) peripheral-blood sample, kept only if that sample has *m*_*d*_ ≥ 2,000 cells, with *y*_*d*_ candidate progenitors among them. Within each eligible study *s*, with at least five case and five healthy control donors and progenitors in both groups (where healthy controls in multi-disease cohorts such as COMBAT and van der Wijst serve each contrast from that cohort and are modeled as separate disease contrasts), a binomial generalized linear model with logit link of (*y*_*d*_, *m*_*d*_ − *y*_*d*_) on disease status gives the log odds ratio 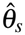; sensitivity models for COVID-19 additionally include each donor’s mean-centered log lymphocyte sequencing depth ln(*U*_*d*_*/L*_*d*_) (total UMIs *U*_*d*_ divided by cells *L*_*d*_ across T, B and NK/ILC cells in the same earliest sample, avoiding myeloid cell-size shifts) and, in studies with at least five cases and five controls of recorded male or female sex and nonzero progenitors in each group, an indicator for donor sex. Severity trends instead use an ordinal step, 0 for healthy, 1 for asymptomatic, mild or moderate and 2 for severe or critical COVID-19 across the multi-severity studies, and 0 to 3 from healthy volunteers to critical disease in COMBAT. The GLM standard error of 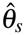 is scaled by 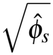 to 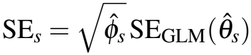 using the quasi-binomial Pearson dispersion 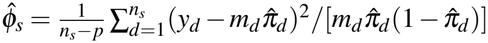 over the study’s *n*_*s*_ donors, *p* model parameters and fitted progenitor fractions 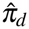. Across the *K* ≥ 2 studies of a disease or severity contrast, study log odds ratios 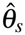 with inverse-variance weights 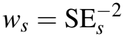 and fixed-effect mean 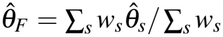 are pooled by the DerSimonian-Laird random-effects estimator [73], with between-study variance 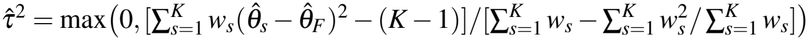, random-effects weights 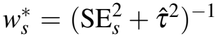, pooled estimate 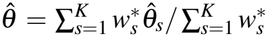, standard error 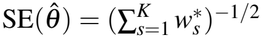, Wald 95% confidence intervals exp 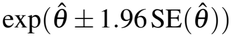 and two-sided normal P values; a disease with one eligible study keeps that study’s quasi-binomial estimate. Across the nine evaluated diseases, P values are adjusted for multiple testing by both the Benjamini-Hochberg false discovery rate [60] and the Holm-Bonferroni procedure. Specificity of mobilization to COVID-19 relative to chronic conditions is tested by a two-sided Wald contrast 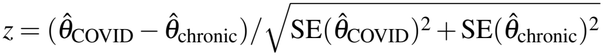 comparing the pooled COVID-19 log odds ratio against the DerSimonian-Laird random-effects pool of the nine study-level contrasts across the six chronic viral, autoimmune, post-infectious and clonal conditions, leaving out sepsis and the COVID-negative patients with moderate to critical illness recorded as respiratory system disorder.

### Statistics and reproducibility

For benchmark arms with *n*_*A*_ ≥ 2 and *n*_*B*_ ≥ 2 runs, two-sided P values and 95% confidence intervals exp 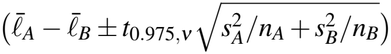 for the geometric-mean time ratio *R* come from Welch’s two-sample t-test on the log runtimes *ℓ*_*Ar*_ = ln*t*_*Ar*_ of runs *r*, with sample variances 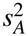 and 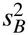 and Welch-Satterthwaite degrees of freedom *ν*, while arms with a single completed run, stage fold ratios, scaling exponents and total-time savings including input preparation are reported descriptively; times beside ratios are arithmetic means, and ranges span the stated runs.

Repeated benchmark executions measure computational and cloud I/O variability, not biological replication. Uncertainty in reference-mapping disagreement is quantified by a nonparametric donor bootstrap: 1,000 resamples of query donors with replacement, within each held-out study or across the pooled ladder query donors, summing the donors’ confusion matrices in each resample and taking the 2.5th and 97.5th percentiles (seed 4444), omitted when fewer than three donors exist. Aside from these donor-bootstrap intervals on mapping accuracy and disagreement, point estimates of mapping accuracy, adjusted Rand indices, normalized mutual information and AUROCs are reported descriptively, and mapping methods, run once each, were not formally tested against one another. Nested mouse embryo subsamples keep the rows with the smallest SplitMix64 hash of their row index offset by a fixed 64-bit sampling key (11789696437827111060) plus seed 0 [74], so each smaller subsample is contained in the larger ones. No statistical method was used to predetermine sample sizes, which are all the cells, donors and studies in public repositories that met the stated inclusion rules; randomization and blinding do not apply to computational analyses of public datasets.

## Data availability

All analyzed datasets are public, and no new sequencing data were generated; Supplementary Notes 1 to 25 describe the datasets and cite their sources. CZ CELLxGENE Discover accessions are dcfd4feb-18a3-4b30-81d7-1b0c544a8ab3 (version 1bc30289-9565-4099-abf9-3326328c11ac; mouse embryo atlas), 53d208b0-2cfd-4366-9866-c3c6114081bc (version 1c88f927-bcbb-4bb1-9881-281842945a2d; Tabula Sapiens), 8e10f1c4-8e98-41e5-b65f-8cd89a887122 (version 937f8304-300e-46ff-a84c-28df40b59d2b; human brain cell atlas neurons) and 37a17b78-4864-4a42-b67b-31c00962795a (version 0e853475-e298-4b09-881a-ed0b60d5a8c9; PsychAD Mount Sinai prefrontal cortex), and the Census benchmarks used CELLxGENE Census release 2025-11-08. Tahoe-100M was read from Hugging Face dataset tahoebio/Tahoe-100M at revision 2dc57900b7981cfcf5e211527169a0b006546a95. Prepared stores of the 10x Genomics 1.3 million mouse brain cells and 5,000 PBMCs, the Tabula Sapiens copy used for remote queries, the Kang PBMCs (GEO GSE96583), the Bastidas-Ponce mouse pancreas cells (GEO GSE132188), the 10x PBMC CITE-seq data and the TEA-seq PBMCs (Zenodo 6360802) are in the cytearc_docs folder of the public Nygen/cytebase bucket on Hugging Face.

Lung reference-revision and mapping analyses used CELLxGENE datasets 066943a2-fdac-4b29-b348-40cede398e4e (version 688185ad-11c2-4172-a53a-f4f1f4076860; Human Lung Cell Atlas core), 9f222629-9e39-47d0-b83f-e08d610c7479 (version 5f863718-02a6-46cb-83e7-52be988b915b; Human Lung Cell Atlas full) and f14bc322-1322-4184-8d16-409557525ea5 (version c3d9262e-0dc5-4eca-bf20-56e6d96d0306; Natri cohort), and held-out tissue mapping read CELLxGENE dataset versions 4078abd1-063d-4b35-aa04-324dddf8244e (Hao), 4554d5bf-9182-47af-8caf-6b2c93513829 (Yoshida), 8d7fad46-2f37-4aa0-ac9d-babbc36efb6f (Wilk), 5e491a70-a1ea-4469-9ffa-9ddff0ec3910 (Lee), 255ce81a-eb4e-48ff-9f18-e50455ccb24d (Lake), c171ca0b-5e46-4260-9dc2-c03486499308 (Stewart) and 0fe5eee4-380d-4bd9-8735-ada5e03021d9 (Reck). Multimodal benchmarks used COMBAT CITE-seq (Zenodo 6120249), Stephenson CITE-seq (processed H5AD `haniffa21.processed.h5ad` from the COVID-19 Cell Atlas; raw data under ArrayExpress E-MTAB-10026) and the Hao PBMC atlas (GEO GSE164378; samples GSM5008737 and GSM5008738).

CyteBase release 2610, with its catalog (`catalog/cytebase.duckdb`), is available at https://huggingface.co/buckets/Nygen/cytebase-2610 and through the CyteBase client, and each experiment’s records name the stores it read. CyteBase is maintained by Nygen Analytics AB on Hugging Face’s free public-dataset infrastructure with open, unauthenticated access, and because each store records the identifiers and SHA-256 checksum of its source H5AD, any store can also be rebuilt independently from CZ CELLxGENE Discover with CyteArc’s open-source importer. The datasets, annotations and motif collections that the experiments read, among them GENCODE release 32 and JASPAR 2024, are listed with their sources and licenses in `THIRD_PARTY_NOTICES.md` of the manuscript repository. Result records, analysis scripts and experiment configurations are available at https://github.com/NygenAnalytics/cytearc-manuscript.

## Code availability

CyteArc (Python 3.13 to 3.15) is released under the BSD 3-Clause license, with bundled components under the MIT and Biopython licenses, at https://github.com/NygenAnalytics/CyteArc and on PyPI (https://pypi.org/project/cytearc/), with documentation at https://docs.nygen.io/CyteArc/. The CyteBase client is available at https://github.com/NygenAnalytics/CyteBase and on PyPI (https://pypi.org/project/cytebase/), with documentation at https://docs.nygen.io/CyteBase/. Benchmark runners, analysis scripts and the code that builds the figures and tables are available at https://github.com/NygenAnalytics/cytearc-manuscript.

## Acknowledgements

We thank Göran Karlsson for comments on the manuscript. This work was supported by Vinnova (grant 2025-03766) to Nygen Analytics AB.

## Author contributions

PD conceived and led the project, acquired funding and wrote the first draft. PD and GA designed the methods, analyzed the results and made the figures. PD, GA, YS and SV developed the software. PD, GA and YS ran the experiments. GA curated the data and provided the compute and CyteBase infrastructure. All authors tested CyteArc, wrote its documentation, and reviewed, edited and approved the manuscript.

## Additional information

Supplementary Information (Supplementary Notes 1 to 25 and Supplementary Figs. 1 to 5) is available for this paper. Correspondence and requests for materials should be addressed to Parashar Dhapola.

## Extended Data

**Extended Data Table 1.**
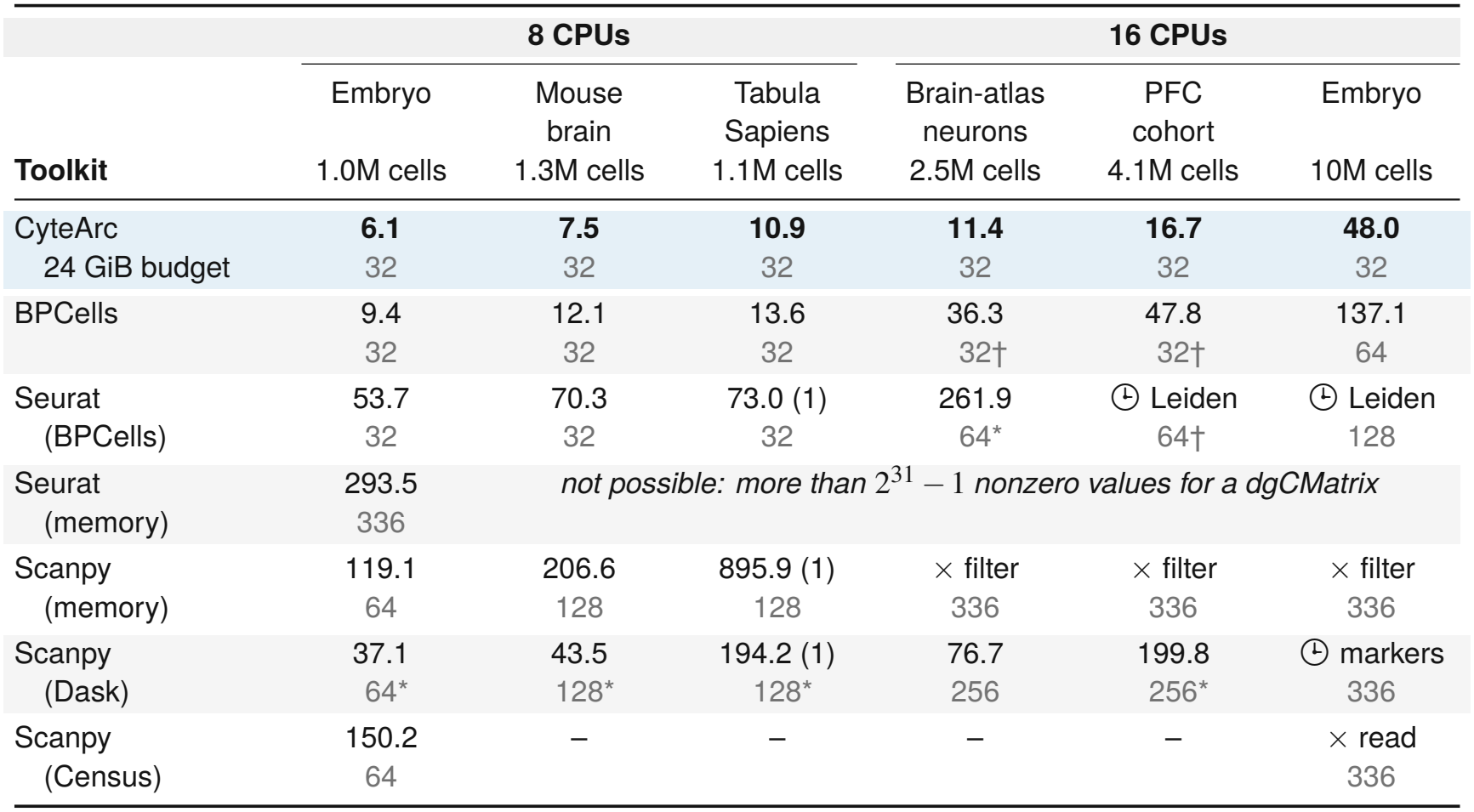
Analysis time and container memory across six 1- to 10-million-cell benchmarks. Mean analysis minutes (reopening prepared input through marker testing; bold, fastest per dataset; *n* = 3 unless noted in parentheses) and allocated container GiB (grey; observed peaks in Fig. 2c). CyteArc used a 24 GiB budget in 32 GiB containers (4 to 24 GiB sweep in Fig. 2i); runs used 8 CPUs up to 1.3M cells and 16 CPUs above (Methods; Supplementary Note 9). ×, out of memory; 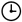, time limit; –, not run; *, initial tier tried; †, UMAP initialized from scaled PCA; PFC, prefrontal cortex.

| Toolkit | 8 CPUs |  |  | 16 CPUs |  |  |
|---|---|---|---|---|---|---|
|  | Embryo<br>1.0M cells | Mouse brain<br>1.3M cells | Tabula Sapiens<br>1.1M cells | Brain-atlas neurons<br>2.5M cells | PFC cohort<br>4.1M cells | Embryo<br>10M cells |
| CyteArc<br>24 GiB budget | **6.1**<br>32 | **7.5**<br>32 | **10.9**<br>32 | **11.4**<br>32 | **16.7**<br>32 | **48.0**<br>32 |
| BPCells | 9.4<br>32 | 12.1<br>32 | 13.6<br>32 | 36.3<br>32† | 47.8<br>32† | 137.1<br>64 |
| Seurat<br>(BPCells) | 53.7<br>32 | 70.3<br>32 | 73.0 (1)<br>32 | 261.9<br>64* | 🕒 Leiden<br>64† | 🕒 Leiden<br>128 |
| Seurat<br>(memory) | 293.5<br>336 | *not possible: more than* $2^{31}-1$ *nonzero values for a dgCMatrix* |  |  |  |  |
| Scanpy<br>(memory) | 119.1<br>64 | 206.6<br>128 | 895.9 (1)<br>128 | × filter<br>336 | × filter<br>336 | × filter<br>336 |
| Scanpy<br>(Dask) | 37.1<br>64* | 43.5<br>128* | 194.2 (1)<br>128* | 76.7<br>256 | 199.8<br>256* | 🕒 markers<br>336 |
| Scanpy<br>(Census) | 150.2<br>64 | – | – | – | – | × read<br>336 |

