## Supplementary Information for "CyteArc: in-place analysis and reuse of remote atlases at 100-million-cell scale"

---

**Contents**


- Supplementary Note 1. Overlay mounts and remote benchmarks
- Supplementary Note 2. CyteBase release and remote queries
- Supplementary Note 3. Blood archive interferon screen
- Supplementary Note 4. Virtual composites and baselines
- Supplementary Note 5. Blood composite and copying baselines
- Supplementary Note 6. Circulating progenitor screen and integration
- Supplementary Note 7. Progenitor disease and severity models
- Supplementary Note 8. Count layouts and read requests
- Supplementary Note 9. Benchmark datasets and comparator runs
- Supplementary Note 10. Comparator workflow parameters
- Supplementary Note 11. Stage profiling and cloud resource trace
- Supplementary Note 12. Tahoe-100M runs
- Supplementary Note 13. Memory budget compliance
- Supplementary Note 14. Artifact reuse and reference revision
- Supplementary Note 15. Reference mapping benchmarks
- Supplementary Note 16. Tissue, corpus, and perturbation mapping
- Supplementary Note 17. Multimodal WNN benchmarks
- Supplementary Note 18. Multimodal T-cell reference
- Supplementary Note 19. Documented non-equivalences
- Supplementary Note 20. Reduction, clustering and marker checks
- Supplementary Note 21. Cluster agreement across toolkits
- Supplementary Note 22. Evaluation against experimental data
- Supplementary Note 23. Integration and mapping agreement
- Supplementary Note 24. Droplet, hashtag and regulatory agreement
- Supplementary Note 25. Scoring, imputation and fate agreement
- Supplementary Figs. 1 to 5


### Supplementary notes

#### Supplementary Note 1. Overlay mounts and remote benchmarks

Mounting a remote store copies its cell and gene metadata tables locally while streaming count chunks and stored artifacts from the remote source on demand. Mounting the 10,000,000-cell mouse embryo store from object storage created a 771.1 MB local user store (769.8 MB for cell tables, or 1.2% of the 64.8 GB source store), leaving the 5.93 GB of stored results in object storage; querying the stored clustering returned it in 19.28 seconds with 0 bytes written locally, and object listings before and after all operations verified that the source store remained unchanged. On the 1,000,000-cell embryo workflow (8 CPUs), CyteArc ran 2.3-fold faster from object storage than Scanpy on Dask over remote Zarr (10.3 versus 24.6 minutes; 95% CI 1.4-4.0, P = 0.020), came within a minute of local-only BPCells (9.4 minutes), and, on local disk, ran 1.3-fold faster than BPCells (7.1 versus 9.4 minutes; 95% CI 1.1-1.6, P = 0.017) and 3.9-fold faster than Scanpy on Dask (28.2 minutes; 95% CI 2.0-7.4, P = 0.010; Fig. 1b). Fold changes are ratios of geometric means over three runs per arm, and the minutes are arithmetic means. The CyteArc and Scanpy on Dask times come from three dedicated full-workflow runs per backend in 8-CPU, 32 GiB containers (CyteArc with a 24 GiB budget and 8 workers), separate from the toolkit comparison's runs, in which CyteArc averaged 6.1 minutes (using the 48-link, depth-50 HNSW settings of Supplementary Note 9, where index build and neighbor query took 17 seconds versus 47 seconds under the default 24-link, depth-200 settings used here, accounting for roughly half of the 1-minute local-disk difference) and Scanpy on Dask averaged 37.1 minutes in a 64 GiB container (Extended Data Table 1); the BPCells times are the toolkit comparison's. Scanpy on Dask took similar times from object storage and from local disk (24.6 versus 28.2 minutes; P = 0.57). Mounting the Wilk COVID-19 blood store [1] over the network created a 5.9 MB local store and completed a six-gene interferon comparison across donor CD14-positive monocytes in 8.56 to 10.37 seconds across three runs (mount included), streaming 32.8 MB of counts.

#### Supplementary Note 2. CyteBase release and remote queries

CyteBase release 2610 comprises 1,372 primary-count stores (including 639 spatial stores) from 376 CZ CELLxGENE Discover collections [2] (178.0 million total observations, 163.6 million primary, across nine species in 1.09 TB of cloud objects; 1,009 human and 329 mouse stores), spanning 715 tissue, 324 disease, and 1,124 cell-type annotation values; among human non-spatial stores with at least 10,000 primary cells, 21 disease annotations other than normal span at least three collections, led by COVID-19 (28 stores from 20 collections). Across the release, cell-major count arrays occupy 634.9 GB and gene-major copies 434.3 GB (a median addition of 75% per store), an estimated 3.04 combined bytes per nonzero count. The Tabula Sapiens benchmark queried the 1,136,218-cell CyteArc store (12.2 GB of count chunks) in the `cytearc_docs` folder of the public Nygen/cytebase bucket [3, 4]: across three runs, opening took 3.90 to 4.45 seconds and reading the UMAP coordinates, cell-type labels, and MS4A1 counts of the 114,495 B cells took 4.49 to 5.66 seconds without a local store, with first open plus query transferring 68.9 MB of HTTP response bodies in a mean of 9.0 seconds (0.20 MB open and 68.7 MB query; 68.6 MB returned to Zarr) and stateless repeat queries on the open store taking 2.45 to 2.78 seconds (mean 2.6 seconds; 66.4 MB over HTTP; 66.3 MB returned to Zarr). The first open plus query ran 6.7-fold faster with 67-fold less network traffic (comparing HTTP response bodies against container network-interface bytes) than the same B-cell MS4A1 query in CELLxGENE Census on Amazon S3 (60.6 seconds and 4.65 GB received on container network interfaces, without UMAP) and 25-fold faster with 653-fold less data than downloading the 45.0 GB H5AD (222 seconds; Fig. 1c).

#### Supplementary Note 3. Blood archive interferon screen

The 10-gene blood screen summed panel counts per donor and coarse cell type across all 82 blood-containing stores (33.5 million primary blood cells among 45.7 million total cells; 3,890 scored donors; 257.6 GB of count arrays) in 19.6 minutes of summed job wall time across two jobs while transferring 12.1 GB, 4.7% of the stores' count-array bytes: 67 stores in 14.9 minutes (8.29 GB transferred, 6.5% of their count arrays, 3.6 GiB peak RAM) and 15 stores in 4.7 minutes (3.81 GB transferred, 2.9% of theirs, 4.1 GiB peak RAM; Supplementary Fig. 1b). The 18-gene interferon screen excluded 11 stores of sorted, enriched, stimulated, or cell-product samples, removed matching samples from the remaining 71 blood stores (31.8 million primary blood cells; 3,586 donors; 18.1 minutes), and scored each donor's earliest sample within each study and lineage when both disease and healthy groups had at least five donors (Methods), spanning

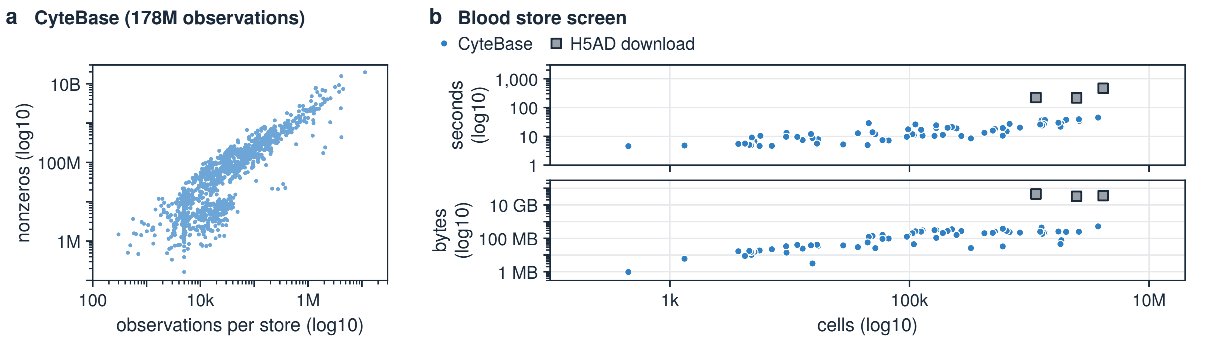


**Supplementary Fig. 1. CyteBase store scale and in-place screening cost** (**a**) Observations and estimated nonzero counts across all 1,372 stores of CyteBase release 2610 (log10 axes). (**b**) Seconds and bytes streamed per human blood store in the 10-gene screen (66 of the 82 stores; 16 read while a job waited on the host's request quota are left out) versus three full H5AD downloads (squares; log10 axes).

13 interferon-driven contrasts (acute viral infections and type-I-interferon-associated autoimmune diseases: 10 for COVID-19 and one each for systemic lupus erythematosus, juvenile dermatomyositis, and influenza) and 10 non-interferon control contrasts (all other qualifying diseases: common variable immunodeficiency, cytomegalovirus infection, respiratory system disorder, rheumatoid arthritis, sepsis, post-COVID-19 disorder, and clonal hematopoiesis). Donor AUROC exceeded 0.50 in every evaluated lineage of all 13 interferon-driven contrasts (0.55 to 1.00; the AUROCs are descriptive). In T cells, every interferon contrast scored above all 10 control contrasts (0.64 to 1.00 versus 0.12 to 0.54), and in dendritic cells, the nine interferon contrasts scored there exceeded the eight control contrasts scored there (0.74 to 0.94 versus 0.29 to 0.63; Fig. 1d).

### Supplementary Note 4. Virtual composites and baselines

The four-study COVID-19 composite combined 233,870 primary COVID-19 and healthy cells from the Wilk, Lee, Arunachalam, and Schulte-Schrepping stores [1, 5–7] (the latter two as harmonized by Jin et al. [8]) across 19,405 shared genes, running without cell filtering using 2,000 variable genes, 30 principal components, 30 neighbors, and Leiden clustering (resolution 1.0) on 8 CPUs under a 16 GiB budget with the source chunk cache disabled. Across three runs, every entry of the virtual composite (zeros included) equaled its source count across all 4.54 billion matrix values, and Leiden clusters and PCA subspaces matched a materialized local store (ARI = 1.0; largest subspace angle 6.5e-5 degrees, within the 0.05-degree tolerance). Streaming the virtual composite (49.1 MB at creation, 1.23 GB after analysis) directly over HTTP completed the analysis in 133 to 161 seconds, faster end to end than writing a materialized local store (150 to 160 seconds; 2.46 GB after its own analysis) and analyzing it locally (64 to 69 seconds; 214 to 230 seconds total).

### Supplementary Note 5. Blood composite and copying baselines

The ten-study blood composite combined all 4,714,857 primary cells of the COMBAT, Yazar, AIDA, Stephenson, Hao, Rabadam, Binvignat, Lee, Schulte-Schrepping, and Ivanova stores [5, 7–16] across 18,446 shared genes in a 1.09 GB (1.02 GiB) local index with zero copied count chunks. Across the composite, Scanpy on Dask, and cached BPCells runs, all three workflows ran without cell filtering using 2,000 variable genes (excluding mitochondrial, ribosomal, HLA, histone, and sex-linked genes), 30 principal components, 30 neighbors, and Leiden resolution 1.0. With the source cache disabled on 16 CPUs under a 24 GiB budget (three runs), building the composite index took 71 to 81 minutes (reading 16.3

GB) and streaming the unintegrated workflow through Leiden clustering over HTTP took 58 to 60 minutes (streaming 26.3 GiB at 12.1 to 12.3 GiB peak RAM, mean 12.2 GiB); in a separate 128 GiB container run with the default source cache enabled, composite creation took 24.4 minutes and unintegrated analysis 49.9 minutes (15.7 GiB peak RAM), followed by Harmony2 integration [17] (which raised normalized within-cell-type study batch entropy from 0.18 to 0.53 at 0.965 kNN cell-type purity; Fig. 1e) and a full-transcriptome donor pseudobulk over 14,040 donor-by-cell-type groups (streaming 1.16 TiB in 221 minutes). All three copying baselines started from the ten pre-subsetted, gene-aligned local H5AD files exported by the composite, totaling 29.1 GB (27.1 GiB), excluding remote download, cell selection, and gene alignment from their timers. From those local files, in-memory AnnData concatenation [18] took 286 to 344 seconds and peaked at 166.7 to 166.8 GiB RAM (14-fold higher memory and 27-fold higher local disk from the 29.1 GB inputs than the 1.09 GB composite) in a 336 GiB container before any analysis began. AnnData's on-disk concatenation failed on the exported files until cell metadata types were harmonized (378 to 398 seconds), then took 395 to 483 seconds at 21.3 to 22.2 GiB RAM and wrote 89.3 GB (83.2 GiB) for the merged H5AD, bringing total local disk to 118.4 GB (110.3 GiB), 109-fold larger than the composite's 1.09 GB index. Scanpy on Dask [19, 20] analyzed that merged H5AD from local disk in 21 to 24 minutes at 67.4 to 69.2 GiB RAM (mean 68.2 GiB, 5.6-fold higher than the composite), running in Asian and European regions while the composite and BPCells runs ran in US ones. BPCells [21] ingested the ten local files and reached Leiden clusters in 40.5 to 46.0 minutes, peaking at 29.7 to 29.8 GiB RAM (2.4-fold higher) and 134.2 GB (125.0 GiB) of local disk, 123-fold larger than the index (Fig. 1e), against 15.2 GB for the composite store after its own analysis through Leiden.

#### Supplementary Note 6. Circulating progenitor screen and integration

Across the 32,896,001 peripheral-blood cells retained across the 82 CyteBase human blood stores after removing sorted, enriched, or stimulated sample subsets within mixed studies (31.9 million peripheral-blood cells across 80 stores and 1.01 million cells across 2 CD34+-enriched reference stores; Methods), the in-place screen finished in 8.1 minutes (transferring 5.96 GB): it read author cell-type labels, library sizes, and counts of four HSPC markers (CD34, SPINK2, PROM1, and CRHBP), retaining non-malignant cells either annotated as progenitors by the study authors or expressing at least two markers. Outside two leukemia cohorts and one 460-gene targeted panel, that selection identified 33,138 candidate cells in unselected blood (0.10%) and 790,157 in CD34+-enriched stores (78.1%; Fig. 1f). We combined the 33,138 peripheral-blood candidates with 25,000 randomly sampled author-labeled progenitors from each of the two CD34+-enriched reference stores (50,000 reference cells total) [22]. Candidates were extracted per store through single-store composites in eight parallel jobs, the longest taking 34.6 minutes (seven stores had zero candidate cells, and eight stores totaling 52 cells, four sorted or stimulated datasets and four lacking state-panel genes, were excluded), leaving 83,086 cells across 64 stores (41 studies and 2,376 donors) aligned on 12,677 shared genes and analyzed in 7.5 minutes (7.0 GiB peak memory) without additional cell filtering using 2,000 variable genes, 30 principal components, Harmony2 integration by collection [17], 30 neighbors, and Leiden clustering at resolution 1.0. Each of the 21 clusters was assigned to whichever of 12 marker-defined states maximized standardized mean marker expression, separating three mature leukocyte clusters (4,612 mature T/NK, B, and monocyte cells) from 18 progenitor clusters across nine states (78,474 cells: 29,758 from unselected blood and 48,716 from CD34+-enriched stores): multipotent HSC/MPP (39,616 cells), erythroid (11,866), lymphoid-primed (11,332), cycling (3,778), myeloid (3,752), megakaryocytic (2,810), basophil/mast (2,801), dendritic (2,326), and B-lymphoid (193) (Fig. 1g). Across all nine states, unselected blood progenitors matched CD34+-enriched reference state proportions within 3.5 percentage points; the three largest states were HSC/MPP (48.3% versus 51.8%), erythroid (15.3% versus 15.0%), and lymphoid-primed progenitors (13.7% versus 14.9%; Fig. 1h; Fig. 1i).

#### Supplementary Note 7. Progenitor disease and severity models

Donor-level models evaluated 1,234 donors across nine diseases with internal healthy controls (15 studies), counting each donor once within a study by their earliest sample if it had at least 2,000 cells, treating CELLxGENE collections as studies and separating the two independent cohorts republished by Jin et al. [8]; within multi-disease cohorts, the same healthy controls serve each contrast from that cohort and are modeled as separate disease contrasts (COMBAT's 10 healthy volunteers serve both COVID-19 and sepsis, and van der Wijst's 10 serve both COVID-19 and respiratory system disorder), and five donors with matching identifiers, ages, and sexes appear in both the Stephenson COVID-19 and the Yoshida post-COVID-19 cohorts and are counted in each. Within each study with at least five disease and five healthy donors, quasi-binomial logit models (standard errors scaled by Pearson dispersion) estimated disease-versus-healthy log odds ratios, pooled across studies by DerSimonian-Laird random effects [23] (Fig. 1j). In COVID-19 across eight studies (349 COVID-19 and 110 healthy donors), circulating progenitor odds were elevated 1.69-fold (95% CI 1.17 to 2.44, P = 0.0054, rising in seven of eight studies with zero estimated heterogeneity $\hat{\tau}^2 = 0$; Fig. 1k). Two fewer COVID-19 cohorts qualify here than in the interferon screen (Wilk et al. and Ivanova et al. drop out) because the progenitor models require at least 2,000 blood cells in each donor's earliest sample and at least one progenitor cell in both disease and healthy groups. Sepsis was represented by one COMBAT cohort of 22 sepsis and 10 healthy donors (odds ratio 5.12, 95% CI 1.13 to 23.2, P = 0.034). Across all nine diseases, COVID-19's adjusted P value stayed below 0.05 under Benjamini-Hochberg and Holm-Bonferroni correction (0.049 for both, from Wald tests that treat study standard errors as known; versus Benjamini-Hochberg q = 0.15 for single-study sepsis). COVID-19's odds ratio exceeded by 1.95-fold (95% CI 1.33 to 2.87, P = 0.00068) the pooled estimate across the nine study-level contrasts of six chronic viral, autoimmune, post-infectious, and clonal conditions (pooled odds ratio 0.86, 95% CI 0.77 to 0.97, P = 0.013, with three cytomegalovirus studies carrying 77% of the inverse-variance weight; one study's 48 healthy controls are shared between cytomegalovirus and rheumatoid arthritis, carrying 32% of pool weight), a pool that leaves out sepsis and the COVID-negative patients with moderate to critical illness recorded as respiratory system disorder. Systemic lupus erythematosus, which was interferon-driven in the screen (Fig. 1d), showed no elevation on its own (odds ratio 0.81, 95% CI 0.60 to 1.09). Adjusting within each study for log lymphocyte sequencing depth across all eight COVID-19 studies (459 donors) yielded an odds ratio of 1.66 (95% CI 1.14 to 2.42, P = 0.0088), and adjusting jointly for lymphocyte sequencing depth and donor sex across the six studies recording sex in at least five cases and five controls (395 donors) yielded 1.87 (95% CI 1.25 to 2.79, P = 0.0022). COVID-19 progenitor elevation held across five sensitivity definitions: odds ratios were 1.63 (95% CI 1.12 to 2.36; eight studies) for marker-defined cells alone, 2.09 (1.29 to 3.39; four studies) for author labels alone, 1.69 (1.15 to 2.49; seven studies) restricting to cells in progenitor states, 1.70 (1.15 to 2.50; seven studies) excluding megakaryocytic progenitors, and 1.79 (1.21 to 2.65; seven studies) restricting to stem and multipotent progenitors. Across the five COVID-19 studies with at least five donors per class (healthy = 0; mild = 1, including asymptomatic and moderate donors; severe = 2, including critical donors), progenitor odds rose 1.85-fold per step (95% CI 1.37 to 2.50, P = 7.1e-5); among patients alone, severe-versus-mild contrasts varied across studies (pooled odds ratio 1.36, 95% CI 0.55 to 3.36; between-study variance 0.81), largely from one study, Yoshida's, whose mild class held four newborns and one donor recorded only at postnatal stage (between-study variance 0.17 without that study), whereas within the uniformly graded COMBAT cohort (healthy, mild, severe, and critical COVID-19; 2.05-fold per step, 95% CI 1.69 to 2.48, P = 1.8e-13), pooled progenitor rates rose monotonically at every consecutive grade from 14.2 progenitors per 10,000 blood cells in healthy volunteers to 16.6 in mild, 37.4 in severe, and 87.0 in critical COVID-19 (Fig. 1l).

#### Supplementary Note 8. Count layouts and read requests

A gene-wise pass (variable-gene selection, gene-wise normalization, or marker testing) reads bands of genes across all cells. Reading a gene band from the cell-major array alone decodes every touched chunk across thousands of unused genes. Across the 1,372 primary-count stores in CyteBase, storing a paired gene-major count array alongside the cell-major array adds a median of 75% to cell-major storage, and under the planned default layout of the 10,000,000-cell mouse embryo sample [24] (computed from planned chunk geometry), decode amplification reaches 91 bytes decoded per byte used from the cell-major array alone, compared with 1.0 from the gene-major copy (a 98.9% reduction in decoded bytes; the gene-major copy compresses to 23.2 GB, adding 66% to the 35.2 GB cell-major array).

Eliminating decode amplification alone does not bound the Zarr range reads issued per pass, which we counted with a store wrapper (each read also fetches its shard's index, so object-store GET requests are roughly twice these counts, for example 21,390 versus 167,624 to 167,625 count GET requests at 10,000,000 cells). Under fixed byte targets, increasing cell count both narrows each memory-bounded gene band (producing more bands) and splits each band across more cell chunks. With the default 1 GB read units and 100 MB chunks, the variable-gene pass issued 20,976 range reads at 5,000,000 embryo cells and 83,812 at 10,000,000 cells. Widening the cell span of gene-major shards at import (2 GB read units with 200 MB chunks at 5,000,000 cells and 100 MB chunks at 10,000,000 cells) reduced variable-gene range reads to 5,336 and 10,695, a 7.8-fold reduction at 10,000,000 cells (Supplementary Fig. 2a). Using fixed cell selections, 1,000 variable genes, and Leiden labels across three runs per layout under a 24 GiB budget (embryo layout jobs were not pinned to one region), both layouts produced identical normalized values and marker statistics, and the wider layout reduced 5,000,000-to-10,000,000-cell runtime scaling exponents (fitted to mean pass seconds across three runs per size) from 1.44 to 0.95 for variable-gene selection, 1.41 to 0.95 for normalization, and 1.10 to 1.02 for marker testing (within run-to-run variation; Supplementary Fig. 2b). At 5,000,000 and 10,000,000 cells, the wider layout gave longer single-pass wall times on marker testing (and on variable-gene selection at 5,000,000 cells), whereas on all 95,624,334 Tahoe-100M cells [25], widening gene-major cell spans by lowering the chunk target from 200 MB to 25.5 MB under 2 GB read units (which narrows gene-major shards in genes so each shard spans nearly eightfold more cells) cut variable-gene range reads 7.9-fold from 3,762,600 to 476,596 and pass runtime 4.5-fold from 196 to 43 minutes (both variable-gene passes ran in us-central, comparing the single run on the 200 MB layout against the first of three on the 25.5 MB layout; the 200 MB marker pass reused the scaled run's Leiden labels). One run on the 200 MB layout matched the first of three runs on the 25.5 MB layout with identical retained cells, variable genes, and marker genes (maximum AUROC difference 0).

#### Supplementary Note 9. Benchmark datasets and comparator runs

The six 1- to 10-million-cell benchmarks comprised the 1,000,000-cell mouse embryo sample [24], Tabula Sapiens (1,136,218 cells, omitted from Fig. 2b and Fig. 2c to avoid overlap near 1 million cells) [3, 4], 1,306,127 embryonic mouse brain cells [26], 2,480,956 human brain neurons [27], 4,140,453 prefrontal cortex cells [28], and the 10,000,000-cell mouse embryo sample [24] (Extended Data Table 1). Analysis time spans reopening each tool's prepared input through marker testing; one-time input preparation is timed separately and builds the CyteArc store, the on-disk BPCells matrices [21], or the uncompressed

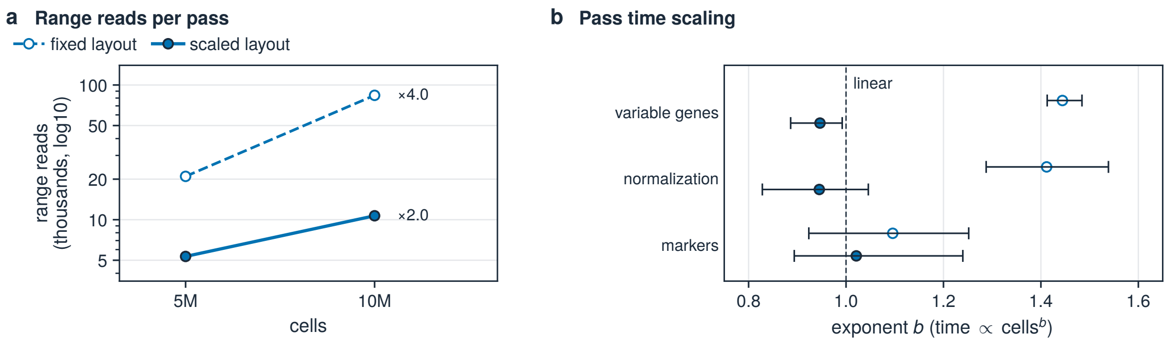


**Supplementary Fig. 2. Gene-major layout scaling of range reads and pass runtimes** (**a**) Variable-gene range reads at 5M and 10M embryo cells on fixed versus scaled layouts (n = 3 per point). (**b**) Runtime scaling exponent b (time ∝ $cells^b$) from 5M to 10M cells across three passes (points, mean-time exponents; bars, run-pair ranges; dashed, b = 1).

H5AD copy read by the R workflows. CyteArc ran three replicates per dataset in 32 GiB containers under a 24 GiB memory budget (8 CPUs up to 1,306,127 cells and 16 CPUs above, peaking at 10.6 to 22.7 GiB). Comparator configurations were in-memory Scanpy [19], Scanpy on Dask arrays [20], Scanpy reading the first embryo cells from CELLxGENE Census [2], in-memory Seurat with presto Wilcoxon testing [29, 30], Seurat on BPCells counts, and native BPCells (using Seurat's Leiden implementation because BPCells' default igraph Leiden ran out of memory at 1,000,000 cells). Any comparator that ran out of memory was retried across 64, 128, 256, and 336 GiB containers, with failures reported at the largest container attempted, except BPCells' igraph Leiden variants, which were stopped manually after failing at 64 GiB (geodesic graph) and 128 GiB (shared-nearest-neighbor graph). CyteArc, native BPCells (with multi-threaded svds PCA; US regions recorded), and Scanpy on Dask ran on non-preemptible containers, whereas all compared 1,000,000-cell embryo runs of in-memory Scanpy, in-memory Seurat, and Seurat on BPCells (plus earlier replicate-1 runs on larger datasets and the Scanpy on Census runs) ran on preemptible containers, while replicates 2 and 3 ran non-preemptible on mouse brain (in-memory Scanpy and Seurat on BPCells) and neurons (Seurat on BPCells); container regions are recorded for native BPCells and omitted from the other public toolkit records.

Dataset-level speedups, percentage time reductions, Welch 95% confidence intervals, and P values compare ratios of geometric means on log-transformed analysis times across three runs per arm, whereas reported minutes and total wall-time savings use arithmetic means. Native BPCells was the only comparator to complete all six datasets; it used scaled-PCA UMAP initialization on the 2,480,956-cell neurons and 4,140,453-cell prefrontal cohort where spectral initialization segfaulted, and peaked at 4.2 GiB on 1,000,000 cells, 15.3 GiB on the prefrontal cohort (all three compared replicates in 32 GiB containers, after an initial 32 GiB attempt was interrupted and completed at 64 GiB), and 41.6 GiB in a 64 GiB container on 10,000,000 cells. Against BPCells, CyteArc ran 1.5- to 3.2-fold faster across the five datasets in Fig. 2b and 1.3-fold faster on Tabula Sapiens, reducing analysis time by 35% at 1,000,000 embryo cells (6.1 versus 9.4 minutes; 95% CI 21-47%, P = 0.0085), 20% on Tabula Sapiens (10.9 versus 13.6 minutes; 95% CI 3-34%, P = 0.038), 38% on 1,306,127 mouse brain cells (7.5 versus 12.1 minutes; 95% CI 29-46%, P = 0.0018), 68% on 2,480,956 human brain neurons (11.4 versus 36.3 minutes; 95% CI 53-79%, P = 0.0063), 65% on 4,140,453 prefrontal cortex cells (16.7 versus 47.8 minutes; 95% CI 64-66%, P = 4.3e-7), and 65% on 10,000,000 embryo cells (48.0 versus 137.1 minutes; 95% CI 51-75%, P = 0.0021), saving 10% to 57% of total wall time across the six atlases when one-time preparation is included (31% to 57% across the five atlases in Fig. 2b, and 10% on Tabula Sapiens).

Comparator failures fell into three categories: 1) full-matrix memory limits, where R's dgCMatrix limit of 2,147,483,647 nonzero entries restricted in-memory Seurat to the 1,000,000-cell embryo sample (1,706,570,607 entries; 238.6 GiB mean peak) and blocked Tabula Sapiens (3,627,322,911 entries), the mouse brain (2,624,828,308 entries), and all larger atlases, while in-memory Scanpy peaked at a mean of 43.7 GiB on 1,000,000 embryo cells and 116.9 GiB in its one Tabula Sapiens run before exhausting 336 GiB during cell filtering from 2,480,956 cells upward; 2) in-memory stages in lazy workflows, where Scanpy on Dask ran batchwise from filtering through PCA yet took up to 212.2 GiB materializing its gene-major marker matrix and 218 to 241 GiB in marker testing at 2,480,956 and 4,140,453 cells, ran out of memory at 247.6 GiB in a 256 GiB container on 10,000,000 cells, and hit the 24-hour call limit at 266.5 GiB in a 336 GiB container, while Scanpy on Census ran out of memory loading the 10,000,000-cell query in both 256 GiB and 336 GiB containers; and 3) R embedding and clustering limits, where uwot's default spectral UMAP initialization segfaulted on the 2,480,956-cell neurons and 4,140,453-cell prefrontal cohort (completed BPCells runs there initialized UMAP from scaled principal components), and Seurat on BPCells counts crashed in default UMAP on the prefrontal cohort, hit the per-attempt time limit in Leiden when initialized from scaled principal components, and timed out at 24 hours in Leiden at 10,000,000 cells.

#### Supplementary Note 10. Comparator workflow parameters

Across the six 1- to 10-million-cell RNA benchmarks (Extended Data Table 1), every toolkit ran to the same targets: 1,000 highly variable genes among genes detected in at least 20 cells, 21 principal components, 11 nearest neighbors (CyteArc excludes the cell itself, whereas Scanpy, Seurat and BPCells count it), 300 UMAP epochs, Leiden clustering at resolution 1.0, and two-sided Wilcoxon rank-sum marker testing across all clusters, each with its documented algorithms for those targets; UMAP and Leiden were seeded with 4444, and native BPCells' multi-threaded UMAP took R's global seed of 4444. By default, CyteArc builds HNSW neighbor indices serially with $M = 24$ (up to 48 links per node in the base layer), construction depth 200, and query depth $\max(200, 2k)$ (used in the multimodal, composite, and mapping runs); the embryo scaling series, storage-backend comparisons, and Tahoe-100M runs kept these depths with parallel HNSW construction, whereas the 1,000,000- to 10,000,000-cell toolkit benchmarks built the index in parallel with $M = 48$ (up to 96 links per node in the base layer) and construction and query depths of 50. On the 1,000,000-cell mouse embryo sample, these toolkit settings maintained high replicate cluster consistency (ARI 0.84 to 0.93 across standard CyteArc runs; 0.87 to 0.95 on the 1,306,127-cell mouse brain) and comparable agreement with published author annotations (major cell clusters: CyteArc NMI 0.70, ARI 0.35 to 0.36 versus BPCells NMI 0.72, ARI 0.39 and Scanpy NMI 0.62, ARI 0.32; fine cell types: CyteArc NMI 0.62 to 0.63, ARI 0.27 to 0.29 across standard runs (0.30 matched) versus BPCells NMI 0.62, ARI 0.30 and Scanpy NMI 0.57, ARI 0.27). The comparators kept cells with at least 10 detected genes within the empirical 1st to 99th percentiles of total counts, detected genes, mitochondrial percentage, and ribosomal percentage, and tested markers on genes detected in at least 20 retained cells. CyteArc's standard arm instead kept cells with more than 10 detected genes within Gaussian-fit 1st to 99th percentiles (counting mitochondrial ribosomal genes as ribosomal), left its default gene families (mitochondrial, ribosomal, cyclin, HLA, histone, and sex-linked genes) out of the variable-gene candidates, and tested markers on all genes.

In-memory Scanpy and Scanpy on CELLxGENE Census [2, 19] applied total-count normalization, log1p transformation, Seurat-flavor variable-gene selection, feature scaling clipped at 10, ARPACK PCA (seed 4466), PyNNDescent neighbor search, spectral-initialized UMAP (spread 1.0, minimum distance 0.5), two-iteration igraph Leiden, and `rank_genes_groups` Wilcoxon testing on the CSR matrix (CSC in the `scanpy_csc` variants at 1,000,000 cells); Scanpy on Dask [20] ran out-of-core on 50,000-cell row chunks with covariance-eigendecomposition PCA and materialized a preallocated CSC matrix in two passes for Wilcoxon testing. In-memory Seurat and Seurat on BPCells counts [21, 29] applied LogNormalize (scale factor 10,000), vst variable-gene selection, ScaleData (clipped at 10), irlba PCA (seed 4466), RcppAnnoy neighbor search, uwot UMAP, leidenbase clustering (algorithm 4), and Wilcoxon testing via presto [30] in memory or BPCells `marker_features` on feature-major bitpacked counts on disk. Native BPCells followed the BPCells tutorial's steps, writing uint32 bitpacked cell-major and feature-major matrices, selecting the 1,000 highest-variance log-normalized genes (scale factor 10,000), caching the normalized 1,000-gene matrix to disk before unclipped z-scoring and multi-threaded BPCells svds PCA, building an RcppHNSW geodesic graph (ef = 100), running multi-threaded uwot UMAP (spectral initialization at 1,000,000, 1,136,218, 1,306,127, and 10,000,000 cells; scaled-PCA initialization at 2,480,956 and 4,140,453 cells, where RSpectra spectral initialization segfaulted), and batchwise tie-corrected Wilcoxon tests from the feature-major matrix; it clustered the geodesic graph through `cluster_graph_seurat` (algorithm 4, leidenbase) because BPCells' igraph Leiden was killed for memory at 1,000,000 cells in containers of up to 64 GiB on the geodesic graph and 128 GiB on the shared-nearest-neighbor graph. A matched-filter CyteArc arm (`cytearc_matched`) that applied the comparators' cell filter and marker genes in the toolkit runs took a mean of 7.1 rather than 6.1 minutes at 1,000,000 embryo cells (distinct from the dedicated local-disk runs (mean 7.1 minutes) under default HNSW settings in Supplementary Note 1), 11.1 rather than 10.9 minutes on Tabula Sapiens, 7.8 rather than 7.5 minutes on the 1,306,127-cell mouse brain, and 49.8 rather than 48.0 minutes at 10,000,000 embryo cells, and remained 1.2- to 2.7-fold faster than BPCells on the four atlases where it ran. CyteArc's standard Gaussian-fit filter kept 5% to 10% fewer cells than the comparators' filter, for example 889,150 against 954,823 of the 1,000,000 embryo cells.

#### Supplementary Note 11. Stage profiling and cloud resource trace

By stage on 10,000,000 embryo cells (three runs per arm; stage fold ratios compare arithmetic mean stage minutes), CyteArc ran Leiden clustering in 1.3 versus 34.1 minutes (26.4-fold), Wilcoxon marker testing in 5.9 versus 25.4 minutes (4.3-fold), neighbor search in 3.5 versus 8.4 minutes (2.4-fold; summing index build, neighbor query, and connectivity graph for CyteArc against queryNeighbors for BPCells), and UMAP in 21.2 versus 57.4 minutes (2.7-fold; summing embedding initialization and UMAP for CyteArc against runUmap for BPCells), compressing the four downstream stages from 125.4 minutes to 31.8 minutes, while BPCells ran cell filtering through PCA in 11.7 versus 16.1 minutes (Fig. 2d).

In a separate non-preemptible US cloud run with default HNSW settings streaming the 10,000,000-cell embryo store from object storage on 16 CPUs in a 64 GiB container under a 48 GiB CyteArc memory budget (distinct from the 32 GiB, 24 GiB-budget local-disk runs above), one-second container telemetry (6,713 samples over 6,843 seconds; the figure panel begins at cell filtering and omits store import) recorded network receive rates up to 327 MB/s during neighbor search and 152 MB/s during the count passes from filtering through PCA (144 MB/s during marker search), up to 15.9 busy cores (from `/proc/stat` busy time during analysis), and container memory peaking at 27.2 GiB during marker search, within the 48 GiB budget (Fig. 2e).

#### Supplementary Note 12. Tahoe-100M runs

Tahoe-100M [25] comprises 95,624,334 cells (139.6 billion nonzero counts, equivalent to 1.52 TiB in uncompressed sparse CSR format with 32-bit values and 64-bit indices) from 50 cancer cell lines across 62,710 genes in 3,388 Parquet files, verified against their published SHA-256 checksums and staged in object storage. Three independent replicates each imported the Parquet files into a fresh CyteArc store in object storage (2 GB read units and 25.5 MB chunks; peaking at 128.5 to 129.6 GiB during import) and executed every workflow stage as a separate job on 16 CPUs and 256 GiB of RAM under a 192 GiB memory budget, reading inputs from object storage and writing output artifacts back to object storage. Store preparation (cell-major and gene-major import plus store initialization) took 3.82, 4.24, and 3.92 hours (mean 4.0 hours), and end-to-end analysis (reopening the store through marker testing) took 8.10, 9.58, and 10.65 hours (mean 9.4 hours); both sum stage execution times and exclude initial Parquet staging to the bucket, copying the Parquet files to local container disk before import, container start-up, and each job's first store opening (Fig. 2f). UMAP accounted for the largest share of runtime and inter-run spread at 2.5, 4.4, and 4.9 hours (replicates 2 and 3 ran UMAP on Azure hosts whereas replicate 1's host provider was not recorded) and set the analysis memory peak at 77.9 to 78.5 GiB (mean 78.3 GiB), whereas multi-threaded Leiden clustering finished in 14.5 to 16.8 minutes (mean 15.3 minutes). Among count-matrix stages during analysis, marker testing peaked at 54.7 to 56.3 GiB and cell filtering through PCA peaked at 16.4 to 16.5 GiB. Quality control in the first replicate retained 83,881,526 cells, which Leiden clustering partitioned into 43 clusters; 35 clusters drew at least 90% of their cells from a single cell line, with a median single-line purity of 96% (Fig. 2g).

For comparison, rapids-singlecell with Dask reported running the atlas from log-normalization through UMAP and Leiden clustering, without marker testing, in under 20 minutes on an 8-GPU NVIDIA B300 node (described by the authors as eight NVIDIA DGX B300 GPUs), and its single-GPU benchmark timings exclude I/O [31]; the analysis hours here include reading counts from object storage and all-cluster marker testing, which took 54, 46, and 75 minutes (mean 58 minutes). Native BPCells, run once on 16 CPUs and 336 GiB of RAM (2,560 GiB local disk, non-preemptible in us-central, scaled-PCA UMAP initialization, 16 threads for svds PCA) with the settings of Supplementary Note 9, converted the same Parquet files to H5AD shards (10.2 minutes) and on-disk bitpacked matrices (76.5 minutes cell-major and 88.1 minutes feature-major), spent 10.1 hours in PCA, 70 minutes in neighbor search, and 86 minutes in UMAP (peaking at 253.6 GiB), and was closed as a memory kill in Leiden clustering 17.4 hours after job start at a last recorded container peak of 322.2 GiB (Fig. 2c). The companion 200 MB chunk-layout run is detailed in Supplementary Note 8.

#### Supplementary Note 13. Memory budget compliance

CyteArc's memory budget bounds the buffers each batchwise operation reserves to read, decode, compute, and write, excluding heap memory retained in a Python process by earlier steps. Across the 1,000,000-cell (19.1 GiB uncompressed CSR) and 10,000,000-cell (190.6 GiB CSR) mouse embryo samples under a 24 GiB budget and Tahoe-100M (1.52 TiB CSR) under a 192 GiB budget, the batchwise variable-gene, normalization, and PCA passes peaked at 4.6, 6.9, and 16.4 GiB while whole-atlas graph and embedding stages grew with cell count at 3.8, 15.8, and 77.9 GiB (means of three runs at 1,000,000 and 10,000,000 cells; first Tahoe-100M run; Fig. 2h), and marker search, which reserves up to the budget, peaked at 6.5, 22.0, and 55.7 GiB. Across all three Tahoe-100M replicates, batchwise passes peaked at up to 16.5 GiB, UMAP averaged 78.3 GiB, and marker search peaked at 54.7 to 56.3 GiB.

First, a dry-run admission check across both count orientations of stores with the 1,000,000- and 10,000,000-cell embryo dimensions, eight budgets (4 to 64 GiB), and 4, 8, or 16 cores verified that all 96 read plans met the budget, read-width, and per-core constraints, reserving at most 99.8% of the budget (under an 8 GiB budget on 8 cores, cell-major plans reserved 7.73 GiB at 1,000,000 cells and 7.82 GiB at 10,000,000 cells). Second, during the variable-gene pass of the standard 10,000,000-cell local-disk runs, 128 concurrent read workers running alongside 16 compute workers waited 0.5 to 0.7 seconds in total for read slots or memory reservations across 4.3- to 6.0-minute passes, with 0 seconds of compute wait.

Third, sweeping the budget across 4, 8, 16, and 24 GiB from input conversion through PCA on 1,000,000 embryo cells (read-unit and chunk targets of 250/25 MB at 4 GiB, 500/50 MB at 8 GiB, and 1,000/100 MB at 16 and 24 GiB; three runs per budget on 8 CPUs in 32 GiB containers) kept container memory strictly below every limit, with mean peaks of 3.8, 5.6, 8.9, and 11.0 GiB, replicate maxima of 3.85, 5.8, 9.4, and 11.5 GiB, and mean runtimes of 4.0 to 4.5 minutes (Fig. 2i); in a single-run screen at 4 GiB, 250/25 MB targets finished in 4.8 minutes versus 7.8 minutes with halved 125/12.5 MB targets.

Fourth, running the full workflow under an 8 GiB budget (500/50 MB targets, three replicates at 1,000,000 cells and one at 10,000,000 cells) compared isolated fresh processes per stage against a single long-lived process. At 1,000,000 cells, every fresh-process stage stayed below 8 GiB (peaking in marker search at 7.9, 7.2, and 7.4 GiB), whereas in a single process retained memory from upstream stages raised marker search to 8.8, 9.6, and 9.1 GiB while UMAP (run in the single-process arm) peaked at 3.5 to 4.1 GiB. At 10,000,000 cells in fresh processes, PCA peaked at 2.6 GiB, neighbor indexing, querying, and graph construction peaked at 4.4 to 5.4 GiB, and Leiden clustering estimated 8.4 GiB and exited before allocating; in a companion fresh-process run seeded with precomputed 10,000,000-cell Leiden labels (loaded outside CyteArc by the test driver at 9.5 GiB), store creation through normalization peaked at 1.6 to 5.8 GiB and marker search at 6.4 GiB. In the single-process arm at 10,000,000 cells, retained upstream allocations pushed neighbor querying and graph construction to 9.6 and 8.9 GiB, and UMAP estimated 10.9 GiB and exited before allocating.

Fifth, across the embryo scaling series (100,000 to 10,000,000 cells in single-container non-preemptible US object-storage runs: 4 CPUs and 16 GiB at 100,000 cells, 8 CPUs and 32 GiB at 500,000 and 1,000,000 cells, and 16 CPUs and 64 GiB at 5,000,000 and 10,000,000 cells) and Tahoe-100M (95,624,334 cells; 16 CPUs and 256 GiB per stage; three replicates per size), mean analysis core-time progressed from 24.7 core-minutes at 100,000 cells through 73.3 at 500,000, 101.8 at 1,000,000, 380.3 at 5,000,000, and 879.8 at 10,000,000 to 9,065 core-minutes on Tahoe-100M, scaling with a power-law exponent of 0.87 from 100,000 to 95,624,334 cells (18 runs) and 1.00 across the 1,000,000-cell embryo to 95,624,334-cell Tahoe-100M range (12 runs; Fig. 2j); between the 1,000,000- and 10,000,000-cell embryo means alone, the two-point exponent was 0.94 in this object-storage series and 1.20 in the local-disk toolkit runs (48.6 to 768 core-minutes).

#### Supplementary Note 14. Artifact reuse and reference revision

The remote 10,000,000-cell mouse embryo store [24] (8.89 million filtered cells; initial 16-CPU run: 29.7 minutes for the stages Leiden clustering depends on, 58.3 minutes through marker discovery, and 83.9 minutes from the 40.2 GB source H5AD) was mounted read-only into a local user store on 8 CPUs under a 24 GiB budget, reading the authors' major cell-cluster and experiment metadata columns by range requests from the source H5AD. A store wrapper counted remote reads, local writes were measured by user-store size changes, and object listings before and after verified that the remote source remained unchanged (Supplementary Fig. 3a). Requesting the stored clustering returned the remote artifact in 19.28 seconds with 0 bytes written locally, and repeating the initial neighbor query returned the stored artifact in 34 seconds. Branching the stored 644 MB neighbor graph at new Leiden resolutions (r = 0.5 and 2.0; each branch read 646 MB, with the cell selection and feature tables) on 8 CPUs finished in 3.3 and 2.3 minutes while writing only 6.0 and 7.1 MB locally, whereas branching earlier from the stored neighbor index to k = 21 computed neighbor search, graph construction, and Leiden in 11.7 minutes (Fig. 3a) and wrote 2.79 GB locally for the new neighbor table, graph, and labels (1.11 GB, 1.67 GB, and 6.6 MB; Supplementary Fig. 3a). Isolating and reanalyzing the 272,930 endothelial cells branched six clusterings (11, 15, and 21 neighbors at resolutions 0.5 and 1.0) from one shared chain of variable genes, normalized counts, PCA, Harmony2 correction across experiments, and neighbor index [17, 32]; across all requests, including those on the initial whole-atlas analysis, total reads from the remote source were 89.6 GB, and the local user store occupied 1.72 GB after the endothelial reanalysis (23-fold smaller than the 40.2 GB source H5AD) and 4.76 GB after all requests (Supplementary Fig. 3c). On another replicate's stored 10,000,000-cell analysis (48.4 minutes originally across the eleven stages), requesting normalization, PCA, and the neighbor index with matching parameters returned the stored artifacts from object storage in 19.0 to 40.5 seconds (each lookup reading 151 to 448 MB of stored artifacts, compared with 0.06 to 0.07 seconds on a local store with 684-fold fewer cells (14,619 Kang PBMCs [33])), and forking the neighbor count from k = 11 to k = 15 on 16 CPUs created 5 new downstream artifacts from neighbor search through UMAP and all-gene marker testing, cutting runtime by 27% from 48.4 minutes to 35.4 minutes of recomputed stages (by 24%, to 36.7 minutes, including the three stored-artifact lookups; Supplementary Fig. 3b).

On the four-study COVID-19 composite, running Harmony2 integration after the unintegrated workflow reused all 4 upstream preprocessing and PCA artifacts, avoiding 2.56 GB of remote count reads (0 count reads during the Harmony2 branch) and cutting total runtime by 32% across three runs from 3.4 to 3.5 minutes (mean 3.46 minutes) on a fresh composite to 2.3 to 2.4 minutes (mean 2.36 minutes; Supplementary Fig. 3d; Supplementary Fig. 3e).

In the lung reference-revision experiment, five query stores mapped onto the core Human Lung Cell Atlas [34] were updated to a revised reference adding 239,707 `Kaminski_2020` interstitial lung disease and control cells (excluding its 67,943 COPD cells) with the authors' aberrant basaloid annotation [35]. Updating the Natri query store [36] in place took 12.3 minutes (10.3 minutes for the revised mapping plus 2.0 minutes re-requesting stored first-version results, with 6 seconds to open the store) and added 61 MB, compared with 20.0 minutes and 157 MB for a complete rerun of both reference versions in a fresh store, or 10.6 minutes and 99 MB for mounting and mapping the revised version alone in a fresh store; in place, the updated store recovered the first-version results exactly from stored label transfers in 31 seconds with 0 reference bytes read (Supplementary Fig. 3f). A screen of the Natri cells against the core reference flagged the 5,077 cells beyond the 99th percentile of control-cell distances within their lineage and clustered them. Among 10 single-parameter variants of the screen, the cheapest, reclustering the flagged cells at resolution 0.5 or 2.0, took 114 to 156 seconds and added 37 to 42 kB, and the costliest, mapping onto a core reference extended with the healthy cells of the atlas extension (excluding the Natri study and query donors), took 25.1 minutes and added 1.45 GB.

#### Supplementary Note 15. Reference mapping benchmarks

The blood reference scaling benchmark drew nested subsets of 500,003; 1,499,999; 2,999,999; and 4,292,870 primary cells (nine harmonized coarse types across seven studies [9–15], ranked within study and cell type by one permutation) from the H5AD exports of the ten-study composite, and mapped 100,002 held-out query cells (33,334 each from Lee, Schulte-Schrepping, and Ivanova [5, 7, 8, 16]). CyteArc ran

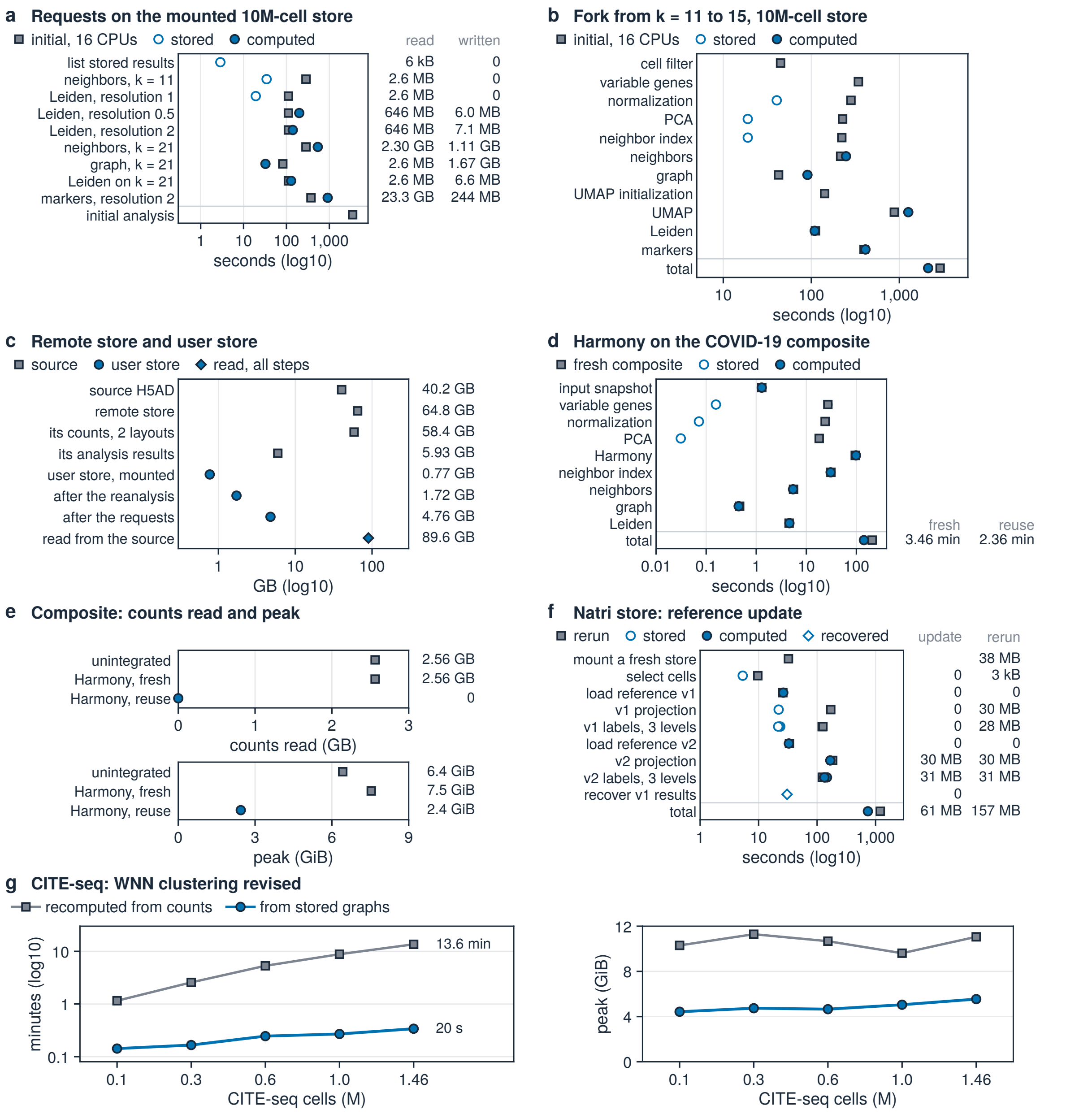


**Supplementary Fig. 3. Stage runtimes and I/O of artifact reuse, forks, and revisions** (**a**) Requests on a mounted 10M-cell embryo store (open, stored; filled, computed) beside original 16-CPU stage times (squares; log10 s); right, bytes read and written. (**b**) Forking k = 11 to 15 on the 10M-cell store: original stage times (squares) beside stored lookups (open) and recomputed stages (filled; log10 s). (**c**) Sizes of the source H5AD, remote store, and mounted user store before and after endothelial reanalysis, plus total remote bytes read (log10 GB). (**d**) Harmony2 stages on the four-study COVID-19 composite run fresh (squares) versus reusing unintegrated artifacts (open, stored; filled, computed; log10 s, means and ranges of n = 3); right, total minutes. (**e**) Remote count bytes read and peak container memory per composite run in d (means and ranges of n = 3). (**f**) In-place Natri lung reference update (circles) versus a fresh two-version rerun (squares), plus v1 recovery (diamond; log10 s); right, bytes added. (**g**) Reclustering CITE-seq WNN graphs at r = 1.2 from stored graphs (circles) versus raw counts (squares): minutes (left) and peak memory (right).

in 64 GiB containers with 2,000 variable genes, 30 components, and 30 neighbors (`save_k` = 30; 20 in the CITE-seq T-cell subtype transfer), both without batch correction and with Harmony2 integration over study plus Symphony query projection [17, 37]. Comparator runs executed once per reference size in 256 GiB containers: Seurat [38] used 2,000 variable genes with FindTransferAnchors and TransferData on 30 dimensions and cached L2-normalized Annoy reference indices; R Symphony [37] used buildReference (2,000 vst genes, 20 principal components, K = 100, Harmony2 over study), mapQuery, and knnPredict with k = 5; and CellTypist [39] trained mini-batch stochastic gradient descent L2-regularized classifiers (batch size 1,000, feature selection enabled, no majority voting) on all genes or, in this scaling benchmark, also on 2,000 batch-aware `seurat_v3` variable genes. Across the four blood scaling references and the held-out blood, kidney, and lung references, marking an existing CyteArc analysis as a mapping reference finished in 2 to 23 seconds and wrote a 0.5 to 0.6 MB manifest pointing to stored artifacts (on the held-out references, where read volume was logged, this step read 0 count bytes and 0.57 to 0.86 GB of stored artifacts). By contrast, Seurat saved its whole uncompressed reference object and Annoy index at 25.5 and 76.5 GB at 0.50 and 1.50 million blood cells (22.4 to 35.8 GB on the held-out tissues), and R Symphony wrote 0.70 and 2.10 GB at those two blood sizes (0.56 to 0.78 GB on the held-out tissues; Supplementary Fig. 4a; Supplementary Fig. 4b). A request log on a 14,619-cell Kang control store [33] confirmed the same separation during query mapping: projecting 12,358 query cells issued 66 requests totaling 3.9 MB in 4.00 seconds without reading any count chunks from the 64 MB reference count arrays.

Across the four blood reference sizes, building the reference from raw counts and annotating the 100,002 query cells in CyteArc took 3.4 to 29.3 minutes without batch correction (9.1 minutes at 1.50 million cells) and 6.5 minutes to 1.7 hours with Harmony2 batch correction (22.7 minutes at 1.50 million cells), compared with 15.2 and 35.6 minutes for Seurat and 18.6 and 56.1 minutes for R Symphony at 0.50 and 1.50 million cells and 7.4 minutes to 1.9 hours for 2,000-gene CellTypist across all four sizes (Fig. 3c). Peak memory (measured as peak container memory) stayed at 13.5 to 17.2 GiB for CyteArc without batch correction (15.1 GiB at 1.50 million cells) and 12.4 to 14.5 GiB with Harmony2 (12.4 to 17.2 GiB overall). By comparison, Seurat peaked at 71.4 and 212.7 GiB and R Symphony at 29.4 and 90.5 GiB at 0.50 and 1.50 million cells before stopping at pre-checks guarding in-memory dgCMatrix loading at 3.00 and 4.29 million cells (4.14 and 5.92 billion nonzero counts, above R's 32-bit dgCMatrix index limit of 2,147,483,647 entries [21]); CellTypist ran out of memory at 246 GiB at 1.50 million cells on all genes and rose from 21.4 to 178.6 GiB on 2,000 variable genes (Fig. 3d). From stored references, CyteArc annotated the 100,002 query cells in 31 to 68 seconds without batch correction (31 and 35 seconds at 0.50 and 1.50 million cells; 68 and 66 seconds at 3.00 and 4.29 million cells) and 20 to 74 seconds with Harmony2 and Symphony projection (20 to 37 seconds at 0.50 and 1.50 million cells and 63 to 74 seconds at 3.00 and 4.29 million cells across both modes). At matched sizes of 0.50 and 1.50 million cells, stored-reference CyteArc mapping ran 13- to 22-fold faster than Seurat's 6.8 and 12.9 minutes and 29- to 79-fold faster than R Symphony's 15.2 and 45.8 minutes, versus 1.7 to 4.3 seconds for a trained 2,000-gene CellTypist model after its 7.4 minutes to 1.9 hours of training and initial annotation (Supplementary Fig. 4c). Across the four reference sizes, label disagreement was 4.7% to 5.0% (accuracy 95.0% to 95.3%) for CyteArc without batch correction, within 0.5 to 1.0 percentage points of Seurat (3.9% to 4.0%; accuracy 96.0% to 96.1%) and R Symphony (4.4% to 4.5%; accuracy 95.5% to 95.6%) and 2.9 to 3.4 percentage points below 2,000-gene CellTypist (7.9% to 8.2%; accuracy 91.8% to 92.1%), and 5.2% to 5.7% (accuracy 94.3% to 94.8%) for Harmony2-integrated CyteArc (Fig. 3e). Across the four rare cell types below 1% of the reference pool, CyteArc without batch correction achieved the highest mean recall at 60.8% to 61.5%

(a mean that includes progenitors, of which no method recovered more than 1 of 142), compared with 58.4% to 58.8% for Seurat and 45.8% to 46.2% for R Symphony (Supplementary Fig. 4d).

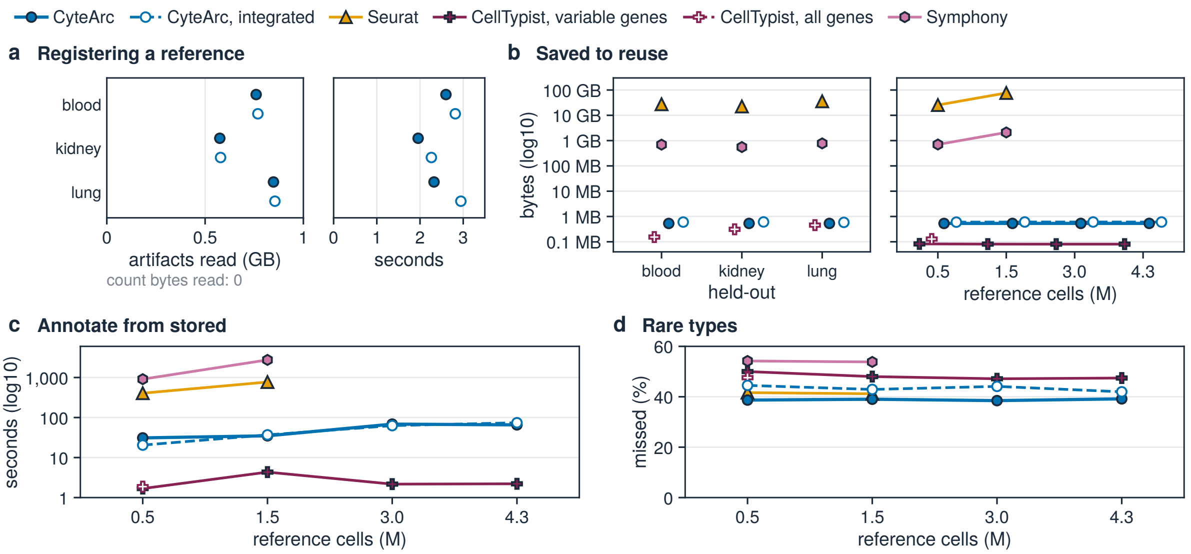


**Supplementary Fig. 4. Reference registration size, query speed, and rare-type recall** (**a**) Artifact bytes read and time to register unintegrated (filled) and Harmony2-integrated (open) references. (**b**) Saved reference size on disk (log10 bytes) across held-out tissues (left) and blood ladder sizes (right). (**c**) Seconds to annotate 100,002 query cells from each stored blood-ladder reference (log10). (**d**) Rare-type cells missed (100 minus mean recall across the four types below 1% of the reference pool).

### Supplementary Note 16. Tissue, corpus, and perturbation mapping

Held-out tissue benchmarks mapped Wilk [1] and Lee [5] blood queries onto Hao and Yoshida [13, 40], 46,957 Reck kidney cells onto a 388,878-cell Lake and Stewart reference [41–43], and 35,300 `Meyer_2019` (Madissoon et al.) lung cells from three donors onto the remaining 546,358 labeled cells of the Human Lung Cell Atlas core [34, 44]. Unintegrated and Harmony2-integrated CyteArc reached 95.8% and 95.5% accuracy on `Lee_2020` and 86.6% and 87.0% on `Wilk_2020` (versus 87.7%, 87.6%, and 83.9% for Seurat, R Symphony, and CellTypist on `Wilk_2020`). On held-out solid tissues, unintegrated CyteArc reached 3.7% disagreement (96.3% accuracy; 1,000-resample donor-bootstrap 95% CI 95.4% to 97.0%; Harmony2-integrated 4.3%) on kidney versus 3.4% for Seurat, 5.6% for R Symphony, and 7.1% for CellTypist (Fig. 3f), and 7.1% disagreement (92.9% accuracy; 95% CI 88.1% to 94.6%; Harmony2-integrated 8.4%) on lung versus 7.4% for Seurat, 7.9% for CellTypist, and 9.9% for R Symphony (Fig. 3g).

For in-place corpus annotation across 14 CyteBase blood stores (21.3 million primary cells across six studies, in seven CELLxGENE collections, from Gong (including Sound Life), Perez, Ren, van der Wijst, Wells, and Yoshida [40, 45–49]), eight jobs each opened the remote 4.29-million-cell reference once in 105 to 137 seconds (1.8 to 2.3 minutes; reading 2.65 GB of artifacts and 0 reference count bytes, imputing missing query genes to the reference mean), annotating each store in 6.9 to 20.7 minutes at 4.7 to 6.5 GiB mapping peak memory per store (8.7 to 11.8 GiB across whole jobs, which also included a post-mapping pseudobulk pass over query counts). Across all 21.1 million typed cells, overall disagreement was 1.8% (98.2% agreement); the two four-store Sound Life jobs accounted for 64.9% of typed cells at 0.4% (CMV-negative) and 0.8% (CMV-positive) disagreement (99.6% and 99.2% agreement), while per-job disagreement across all eight jobs ranged from 0.4% to 7.5% (Ren) and pooled to 95.9% agreement across the six single-store jobs (Fig. 3h; Fig. 3i; Fig. 3j). Finally, in an unintegrated perturbation test mapping 12,358 interferon-beta-stimulated Kang PBMCs onto 11,944 unstimulated control cells [33], raising the neighbor-vote threshold from 0.5 to 0.9 abstained on 2.9% to 59.6% of query cells (at threshold 0.6, abstention was highest in B cells at 36.7% and FCGR3A+ monocytes at 33.5%, while CD14+ monocytes retained 99.3% agreement among accepted cells and dendritic cells abstained at 18.3% with 10.2% agreement among accepted cells, projecting mainly to CD14+ monocytes) and increased label agreement among accepted cells from 85.1% to 98.1%.

**Supplementary Note 17. Multimodal WNN benchmarks**

The CITE-seq scaling series analyzed nested subsets of 100,000; 300,000; 600,000; 1,000,000; and 1,457,527 cells from one permutation of the COMBAT and Stephenson PBMC cohorts (188 shared surface proteins, their 192-antibody panel without its four isotype controls; 228 antibodies on Hao) [9, 12], once per tool and size. Both CyteArc WNN (in 64 GiB containers) and Seurat v5 FindMultiModalNeighbors used 2,000 variable genes and 30 components for RNA, across-cell centered log-ratio (CLR) normalization and 15 components for antibody-derived tags (ADT), L2-normalized reductions, 20 neighbors per modality (deterministic hnswlib in CyteArc; Annoy with Seurat's default 50 trees and knn.range = 200 in Seurat), and Leiden clustering at resolutions 1.0 and 1.2 (native multi-threaded Leiden in CyteArc; FindClusters algorithm 4 with leidenbase in Seurat), with CyteArc completing the initial two-assay workflow in 1.1 to 13.3 minutes across the five sizes (5.4 minutes at 600,000 cells, 41-fold faster than Seurat v5 WNN's 3.8 hours; Fig. 3k). Reclustering the stored WNN graph at resolution 1.2 took 8.5 to 20.4 seconds across all five sizes in CyteArc, running 748-fold faster at 600,000 cells (14.7 seconds) than Seurat reclustering its stored WNN graph with leidenbase (3.1 hours; Fig. 3l; Supplementary Fig. 3g). CyteArc initial-analysis memory peaked at 8.6 to 10.7 GiB and whole-run peak memory stayed at 9.9 to 11.3 GiB across all sizes (timings exclude a one-time Numba compilation warm-up of 69 to 79 seconds, and recomputing the resolution-1.2 clustering from scratch took 1.2 to 13.6 minutes with identical cluster labels to stored-graph reclustering). Seurat v5 [29] ran matched steps with FindMultiModalNeighbors in 128 GiB containers (120 GiB memory stop; 2- to 10-hour time limits by size; input loading excluded), rising in whole-run peak memory from 16.7 GiB at 100,000 cells and 87.6 GiB at 600,000 cells (where it finished the initial workflow and stored-graph reclustering before hitting its 8-hour limit during the fresh recompute) to 118.8 GiB at 1.0 million cells (where it timed out after 10 hours inside WNN clustering) and exhausting the 120 GiB limit during RNA normalization at 1.46 million cells (Fig. 3m). On the 161,764-cell Hao CITE-seq atlas [13] (three replicates), CyteArc finished the initial workflow in 1.9 to 2.6 minutes (compilation included) at 6.7 to 6.9 GiB initial peak memory (6.9 to 7.5 GiB whole-run peak), versus 13.9 to 14.4 minutes at 26.1 to 27.6 GiB whole-run peak for Seurat. On 6,194 TEA-seq PBMCs [50] and 7,299 10x CITE-seq PBMCs [51], replaying Seurat's weighted-neighbor kernel in R on CyteArc's candidate neighbors returned identical neighbors and modality weights within 3e-8.

To compare graph quality without resolution bias, every graph (including Seurat's) was clustered with CyteArc's Leiden across three seeds using the resolution tuned on seed 4466 to match 19 harmonized author cell types (yielding 18 to 20 clusters across seeds for CyteArc WNN and 16 to 21 for Seurat WNN; Fig. 3n; Supplementary Fig. 5a). At that matched resolution, CyteArc WNN exceeded Seurat WNN at every completed size in cluster agreement (mean ARI 0.43 to 0.44 versus 0.35 to 0.36, a +0.07 to +0.08 ARI gain) and raised mean F1 recovery of the ten surface-protein-defined types to 0.44 to 0.45 (versus 0.30 to 0.33 for Seurat WNN or RNA alone). Pooled unintegrated ARI (0.46 for RNA alone) reflected a split between the two cohorts: within COMBAT, CyteArc WNN reached mean ARI 0.77 to 0.78 across all sizes (0.78 at 600,000 cells, versus 0.61 for Seurat WNN and 0.60 for RNA alone), whereas within Stephenson the unintegrated protein graph alone scored ARI 0.19 to 0.21 (versus 0.61 to 0.66 within

COMBAT), pulling unintegrated WNN to 0.33 to 0.37 (versus 0.58 to 0.62 for RNA alone and 0.23 to 0.25 for Seurat WNN). Once Harmony2 corrected for study across all 1,457,527 cells (Fig. 3o), WNN achieved higher overall author-label agreement than RNA or ADT alone (ARI 0.443 across 24 clusters versus 0.414 across 35 RNA clusters and 0.197 across 63 ADT clusters). At the workflow's default resolution of 1.0 on 600,000 cells, CyteArc WNN produced 22 clusters (ARI 0.43), whereas Seurat WNN subdivided the cells into 105 and 112 clusters at resolutions 1.0 and 1.2 (ARI 0.25 and 0.24; forcing CyteArc WNN or RNA alone to 105 and 112 clusters subdivided the 19 coarse author classes to ARI 0.13 and 0.12 or 0.14 and 0.13, respectively).

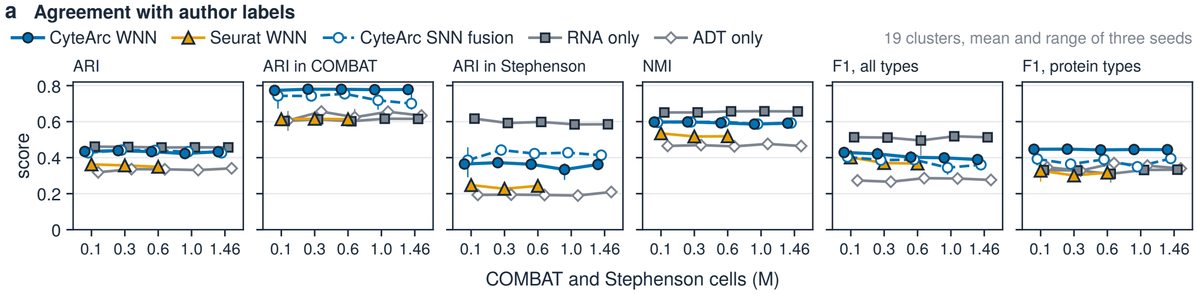


**Supplementary Fig. 5. Multimodal graph agreement with harmonized author cell types** (**a**) Agreement with 19 harmonized author types at 19 clusters across 0.10M-1.46M CITE-seq cells: overall and within-study ARI, NMI, and all-type and protein-type mean F1 (means and ranges, 3 seeds).

#### Supplementary Note 18. Multimodal T-cell reference

For multimodal-to-RNA T-cell transfer, clustering COMBAT's 371,747 T cells [9] at resolution 0.5 on their WNN graph (per-cell CLR-normalized ADT, chosen because across-cell CLR split these T cells partly by per-cell antibody capture) yielded ten clusters, seven named from author labels and surface markers and three small clusters left unnamed (Fig. 3p); RNA-only projection of Hao's 73,060 T cells [13] against that named reference assigned a subtype to 73,034 cells. In a controlled comparison naming every resolution-0.5 WNN and RNA cluster by majority author label across 68,085 Hao T cells in the seven subtypes (with 4,975 Hao T cells of other subtypes not scored), the WNN-defined reference improved RNA-only transfer over the RNA-defined reference overall (83.4% versus 68.3%) and in five of seven subtypes, including gains from 47.4% to 87.5% in memory CD4 T cells and from 79.5% to 92.5% in effector CD8 T cells, held it in mucosal-associated invariant T (MAIT) cells (92.2% versus 92.0%), and lowered it in regulatory T cells (30.8% versus 33.6%), whose WNN cluster was half regulatory T cells (regulatory T, naive CD8, and $\gamma\delta$ T cells showed smaller mRNA-only separation under both references; Fig. 3q). Across nine canonical shared surface markers in the 73,034 Hao T cells assigned by RNA against the named reference, transferred labels reproduced COMBAT's subtype profiles, including TCR V$\delta$2 in $\gamma\delta$ T cells (median CLR 3.006), TCR V$\alpha$7.2 in MAIT cells (1.515), CD57 in CD8 effector cells, and CD25 in regulatory T cells (1.080) (Fig. 3r).

#### Supplementary Note 19. Documented non-equivalences

Table 2 reports numerical agreement under matched inputs and convergence settings. Under other settings the outputs differ where CyteArc follows a published formula that an upstream package departs from, keeps cells an upstream package drops, leaves scaled features unclipped, or searches a narrower neighbor pool; the experiment records keep all comparator and sensitivity runs. In Harmony2 [17, 32], CyteArc evaluates the diversity penalty $\theta\ln[(E+1)/(O+E+1)]$ in each assignment logit, with $O$ and $E$ denoting the observed and expected batch masses in the cluster, using the exact published numerator $E+1$, whereas harmonypy 2.0.2 [52] and R harmony 2.0.5 compute $2E+1$, a discrepancy confirmed as a bug by an R harmony developer (`github.com/immunogenomics/harmony/issues/294`); with the penalty enabled, a single assignment step from a shared state differed by up to 0.0087 from harmonypy and 0.0064 from R harmony. During the linear correction step, CyteArc retains all cells in each cluster's regression (omitting only the batch offset term for batches pruned below the cluster mass threshold), whereas harmonypy drops cells of pruned batches from the regression entirely, producing a maximum single-step difference of 0.022 across 100 clusters. Across complete fits with each tool's default stopping criterion, CyteArc agreed with harmonypy within harmonypy's own seed-to-seed variation (Supplementary Note 23); with a fixed number of rounds instead, the overlap with harmonypy fell just below the lowest overlap between two harmonypy seeds on Kang PBMCs (0.954 versus 0.959 with the penalty disabled) and on both datasets with the penalty enabled. R harmony 2.0.5, by contrast, halts as soon as its objective increases, stopping after two rounds on every seed of the Binvignat PBMCs [15] and yielding a median neighbor overlap of 0.68 with CyteArc. Similarly, Symphony query mapping matches R Symphony [37] within $10^{-5}$ under shared unclipped PCA scaling, whereas R Symphony's default feature clipping shifts projected coordinates by up to 9.45 and corrected coordinates by up to 4.72. In weighted nearest-neighbor (WNN) fusion [13], CyteArc restricts each cell's candidate pool to the union of its stored $k$-nearest modality neighbors, so fusion reuses the stored indices, whereas Seurat searches a wider candidate pool (knn.range = 200) with its own bandwidth; on 7,299 CITE-seq PBMCs and 6,194 TEA-seq PBMCs, the two candidate strategies shared a mean of 0.64 and 0.48 of final edges while producing concordant Leiden partitions at ARI 0.90 and 0.82 [13, 50]. For chromatin accessibility, batchwise randomized LSI with the multimodal workflow's default 5 power iterations and 10 oversamples missed the 0.05-degree tolerance on all 6,194 TEA-seq cells (largest subspace angle 88.4 degrees), as did 20 iterations with 100 oversamples (5.6 degrees), and came within 0.000935 degrees of dense SVD at 80 power iterations and 100 oversamples [53, 54]. Finally, agreement is lower where tools use distinct stopping rules or draw and preprocess inputs differently: GRNBoost2 reproduces arboreto [55] exactly under arboreto's early-stopping rule, whereas under CyteArc's convergence rule the importances correlate at $r = 0.63$ and the 1,000 strongest links overlap at a Jaccard index of 0.16; cell-cycle scores with independent random control genes correlated at 0.995 and 0.987 on pancreas cells [56], and through CyteArc's public scoring function, which also matches PIMREG and JPT1 by their previous symbols FAM64A and HN1 where Scanpy's exact matching finds neither, pancreas G2M scores differed from Scanpy's by up to 0.028 with the same controls, while S scores and PBMC scores matched to about $10^{-16}$; and with each tool's default parameters on shared inputs, median Pearson correlations were 0.95 per gene for MAGIC on CyteArc's normalized values [57] and 0.99 per gene set for AUCell [58], and CellRank alpha fate probabilities on the pancreas data correlated at 0.985 [59].

#### Supplementary Note 20. Reduction, clustering and marker checks

PCA in CyteArc uses an exact Gram solver up to 4,096 features and a batchwise randomized block-Krylov solver above [54, 60]. Across 10 configurations spanning 2,000 to 5,000 features and two block sizes on 3,696 mouse pancreas cells, the largest loading-subspace angles against a dense double-precision SVD were $6.4 \times 10^{-13}$ degrees for the Gram solver and 0.0143 degrees for the randomized solver, well within the 0.05-degree equivalence tolerance [56]. On a fixed input of 1,024 cells and 2,000 genes, the Gram solver stayed within $1.26 \times 10^{-12}$ degrees of the dense solution at every block size, whereas an incremental fit with scikit-learn's IncrementalPCA drifted by 85.1 degrees with 4 blocks and 89.3 degrees with 8 blocks [61]. Batchwise LSI of 25,000 TEA-seq peaks matched dense SVD within 0.002

degrees on a 400-cell sample at 20 power iterations and within 0.000935 degrees on all 6,194 cells at 80, both with 100 oversamples [50, 53] (Supplementary Note 19). Native multi-threaded Leiden produced identical partitions across 1, 4, 8, and 16 threads on a fixed graph of 18,597 Kang PBMCs, achieving modularity 0.8629 compared with 0.8628 for single-threaded igraph [33, 62, 63]. At this graph size, local moves run on one thread and refinement in parallel, so the check did not exercise parallel moves. With artifact reuse disabled, three fresh runs of the RNA workflow in default deterministic mode returned identical intermediate arrays and cluster assignments (ARI 1.0), and parallel mode reached ARI 0.94 between some runs. For differential marker discovery, comparing 6,000 sampled gene-by-cluster pairs on a fixed clustering of the 1,000,000-cell mouse embryo, 1,306,127-cell mouse brain, and 1,136,218-cell Tabula Sapiens datasets [3, 4, 24] confirmed that CyteArc's Mann-Whitney AUROC matched presto and Scanpy within $1.2 \times 10^{-6}$ and its tie-corrected P values agreed with BPCells across all pairs above the double-precision underflow floor [19, 21, 30] (Table 2).

#### Supplementary Note 21. Cluster agreement across toolkits

To separate solver stability from upstream preprocessing differences, we compared cluster membership across the shared retained cells of the 1,000,000-cell mouse embryo sample (877,450 cells) and the 1,306,127-cell mouse brain dataset (1,152,328 cells) across three standard CyteArc runs, three CyteArc runs configured with the comparator cell and gene filters, one BPCells run, and one Scanpy run [19, 21, 24]. Replicate standard CyteArc runs shared an identical principal subspace (smallest canonical correlation 1.0) and achieved high internal partition stability, with pairwise adjusted Rand index (ARI) of 0.84 to 0.93 on the embryo atlas and 0.87 to 0.95 on the mouse brain; these runs build their neighbor indices in parallel, which varies the graph between runs. In contrast, switching only the upstream cell and variable-gene filters within CyteArc shifted the embryo partition to ARI 0.59 to 0.64, and the PCA subspaces constructed by BPCells and Scanpy from their own normalization and variable-gene selections each contained directions nearly orthogonal to CyteArc's subspace (smallest canonical correlations 0.0007 and 0.0025 on the embryo atlas and 0.0014 and 0.0022 on the mouse brain). Cross-toolkit partitions agreed at ARI 0.38 to 0.39 between CyteArc and BPCells, 0.32 to 0.33 between CyteArc and Scanpy, and 0.30 between BPCells and Scanpy on the embryo atlas (0.38 to 0.39, 0.48 to 0.49, and 0.36 on the mouse brain), while the three toolkits reached similar concordance with the coarse author annotations (ARI 0.36 for the first standard CyteArc run, 0.39 for BPCells, and 0.32 for Scanpy). Cross-toolkit differences therefore exceed run-to-run variation and follow the upstream filters, normalization, and variable genes; these comparisons measure consistency between complete workflows, not which partition is biologically more accurate.

#### Supplementary Note 22. Evaluation against experimental data

Where ground-truth experimental labels exist, we evaluated CyteArc's biological recovery directly against experimental benchmarks and standalone tools. Projecting simulated doublets into the store's existing PCA artifact without fitting a second reduction, CyteArc's default doublet detector reached a median AUPRC of 0.48 across the 16 experimentally annotated doublet benchmark datasets of Xi and Li [64], above Scrublet (0.41) and DoubletFinder (0.36) and below scDblFinder (0.55), which it exceeded on 4 of the 16 matrices, and a median AUROC of 0.787 against scDblFinder's 0.795 [65–67]. On 27,718 Kang PBMC droplets with 3,156 genetically demultiplexed doublets [33], CyteArc achieved an AUPRC of 0.59 under package defaults and 0.64 under documentation settings, improving over both Scanpy's Scrublet implementation (0.54) and library-size ranking (0.30). In a clonal lineage-tracing benchmark across 1,110 early hematopoietic progenitors from 725 barcoded clones in the LARRY dataset [68], CyteArc's

Markov absorption fate probabilities predicted neutrophil versus monocyte clonal fate at AUROC 0.76, matching CellRank [59]. Similarly, coupling CyteArc's GRNBoost2 importance links with cisTarget motif pruning found 2,437 consensus SPI1 target genes in at least two of three seeds, 266 of which overlapped experimentally measured PU.1 binding peaks in an independent CUT&RUN assay (10.9% against 6.5% of the 14,207 genes tested), a 1.67-fold enrichment over background ($P = 1.6 \times 10^{-19}$, prespecified one-sided test) [58, 69].

#### Supplementary Note 23. Integration and mapping agreement

Harmony2 [17, 32] in CyteArc was compared with harmonypy 2.0.2 [52] and R harmony 2.0.5 on identical principal components of Binvignat and Kang PBMCs [15, 33]. With the diversity penalty disabled, a single assignment step from a shared state, on 200 cells and six centroids drawn from 6,000-cell subsets of each dataset, differed from harmonypy by up to $4.5 \times 10^{-7}$ and from R harmony by up to $1.4 \times 10^{-6}$. Across complete multi-iteration fits (Supplementary Note 19), CyteArc matched harmonypy within harmonypy's intrinsic seed-to-seed variation: on 9,000 Binvignat PBMCs, root-mean-square coordinate differences were 0.06 of the reference standard deviation, and across ten random seeds the median nearest-neighbor overlap between CyteArc and harmonypy was 0.883 on Binvignat PBMCs and 0.978 on Kang PBMCs, against 0.884 and 0.982 among harmonypy's own seeds. On shared neighbor graphs, integration and cell-type local inverse Simpson's indices (iLISI and cLISI) differed from scib-metrics by up to $1.4 \times 10^{-15}$ in dataset medians and $1.5 \times 10^{-14}$ per cell, on three embeddings of two datasets [70]. For Symphony reference mapping of 500 held-out Krasnow lung cells onto a three-study HLCA reference under shared unclipped PCA scaling, maximum absolute differences against R Symphony were $4.3 \times 10^{-14}$ for projected coordinates, $7.0 \times 10^{-9}$ for soft cluster assignments, and $8.0 \times 10^{-8}$ for corrected coordinates, well within the $10^{-5}$ equivalence tolerance [34, 37, 71]. Replaying Seurat's weighted nearest-neighbor kernel [13] on shared candidate pools and bandwidths (Supplementary Note 17) reproduced modality weights and affinities within $3 \times 10^{-8}$ in three repeated runs on identical inputs on both TEA-seq and 10x CITE-seq PBMCs as well as on Seurat's 300-cell CITE-seq and 48-cell synthetic three-modality reference fixtures, generated with Seurat 5.5.1 under R 4.4.3.

#### Supplementary Note 24. Droplet, hashtag and regulatory agreement

EmptyDrops ambient-RNA cell calling was compared with Bioconductor DropletUtils on the raw 10x PBMC 1k and 10k unfiltered droplet matrices using independent Monte Carlo streams, matching cell calls at a Jaccard index of 1.0 on PBMC 1k and 0.983 on PBMC 10k (6,819 cells called by CyteArc versus 6,756 by DropletUtils) [72]. SoupX ambient contamination fractions matched the R reference within $4 \times 10^{-14}$, with CyteArc's integer-corrected counts differing from SoupX's continuous subtraction by less than one count [73]. Hashtag oligo demultiplexing matched Seurat's HTODemux on the negative, singlet, and doublet classification of all 21,522 cells in the Stoeckius dataset and all 8,438 cells in GSE245108, differing on the sample identity of only one Stoeckius singlet where CyteArc assigns the sole positive hashtag above threshold rather than the largest CLR value [74, 75]. Chromatin accessibility GeneScores across 400 TEA-seq cells, 240,122 peaks, and 256 genes reproduced an independent interval-overlap and TF-IDF reference implementation to machine precision [50]. Given identical motif matches and background peak sets, chromVAR bias-corrected deviations matched the R chromVAR package within $1.7 \times 10^{-14}$ and z-scores within $2.8 \times 10^{-12}$ [76]. Under arboreto's stopping rule, CyteArc's GRNBoost2 reproduced all 1,179,424 importance-weighted regulatory links identically on 2,612 multiome PBMCs [55], and cisTarget motif recovery ranks and AUC scores matched pySCENIC across all 54 tested regulator-by-seed pairs [58, 77].

#### Supplementary Note 25. Scoring, imputation and fate agreement

Cell-cycle S and G2M phase scores matched Scanpy within $3.9 \times 10^{-16}$ on mouse pancreas cells and human PBMCs when evaluated with identical control gene sets and gene lists [19, 56]. AUCell enrichment and weighted gene-set average (WAGGR) scores matched decoupler across 18 gene sets on 4,331 PBMCs within $3.0 \times 10^{-8}$ and $9.2 \times 10^{-7}$ under matched tie-breaking and transformations, with zero difference after rounding decoupler's double-precision AUCell outputs to CyteArc's stored float32 precision [58, 78]. On a shared Markov diffusion operator and step count, MAGIC expression imputation matched magic-impute within $6.7 \times 10^{-6}$ across 4,331 PBMCs and 200 variable genes, matching the single-precision representation of the stored graph weights [57]. Population Balance Analysis (PBA) pseudotime potentials on 3,546 mouse pancreas cells matched the exact dense pseudoinverse solution ($r = 0.999995$ to 0.99999995 as truncated singular components increased) and agreed with a dense truncated SVD of the same graph Laplacian operator within $3.19 \times 10^{-7}$, a check of the solver against its own definition rather than an independent implementation [79]. Finally, Markov absorption fate probabilities matched CellRank on the mouse pancreas trajectory with matched settings within $4.31 \times 10^{-5}$ for the alpha lineage and $3.65 \times 10^{-5}$ for the beta lineage [59, 80], and CyteArc's sparse linear solver matched exact dense linear system solutions within $2.9 \times 10^{-7}$ across four graphs, the 3,546-cell pancreas and 11,944-cell Kang PBMC graphs and two small test graphs [33].

ll.2025.02.017.

[17] Nikolaos Patikas, Hongcheng Yao, Roopa Madhu, Soumya Raychaudhuri, Martin Hemberg, and Ilya Korsunsky. Integration of large, complex single-cell datasets with Harmony2. bioRxiv, 2026.

24-07069-w.

[35] Taylor S. Adams, Jonas C. Schupp, Sergio Poli, Ehab A. Ayaub, Nir Neumark, Farida Ahangari, Sarah G. Chu, Benjamin A. Raby, Giuseppe DeIuliis, Michael Januszyk et al. Single-cell RNA-seq reveals ectopic and aberrant lung-resident cell populations in idiopathic pulmonary fibrosis. *Science*

*Advances*, 6(28):eaba1983, 2020. doi: 10.1126/sciadv.aba1983.

[52] Kamil Slowikowski, John Arevalo, Florian De Rop, jkh, and Jonathan Manning. slowkow/harmonypy: v2.0.2. Zenodo, 2026.

[53] Darren A. Cusanovich, James P. Reddington, David A. Garfield, Riza M. Daza, Delasa Aghamirzaie, Raquel Marco-Ferreres, Hannah A. Pliner, Lena Christiansen, Xiaojie Qiu, Frank J. Steemers et al. The cis-regulatory dynamics of embryonic development at single-cell resolution. *Nature*, 555(7697): 538–542, 2018. doi: 10.1038/nature25981.

[54] N. Halko, P. G. Martinsson, and J. A. Tropp. Finding Structure with Randomness: Probabilistic Algorithms for Constructing Approximate Matrix Decompositions. *SIAM Review*, 53(2):217–288, 2011. doi: 10.1137/090771806.

[55] Thomas Moerman, Sara Aibar Santos, Carmen Bravo González-Blas, Jaak Simm, Yves Moreau, Jan Aerts, and Stein Aerts. GRNBoost2 and Arboreto: efficient and scalable inference of gene regulatory networks. *Bioinformatics*, 35(12):2159–2161, 2018. doi: 10.1093/bioinformatics/bty916.

[56] Aimée Bastidas-Ponce, Sophie Tritschler, Leander Dony, Katharina Scheibner, Marta Tarquis-Medina, Ciro Salinno, Silvia Schirge, Ingo Burtscher, Anika Böttcher, Fabian J. Theis et al. Comprehensive single cell mRNA profiling reveals a detailed roadmap for pancreatic endocrinogenesis. *Development*, 146(12):dev173849, 2019. doi: 10.1242/dev.173849.

[57] David van Dijk, Roshan Sharma, Juozas Nainys, Kristina Yim, Pooja Kathail, Ambrose J. Carr, Cassandra Burdziak, Kevin R. Moon, Christine L. Chaffer, Diwakar Pattabiraman et al. Recovering

gene interactions from single-cell data using data diffusion. *Cell*, 174(3):716–729.e27, 2018. doi: 10.1016/j.cell.2018.05.061.

[58] Sara Aibar, Carmen Bravo González-Blas, Thomas Moerman, Vân Anh Huynh-Thu, Hana Imrichova, Gert Hulselmans, Florian Rambow, Jean-Christophe Marine, Pierre Geurts, Jan Aerts et al. SCENIC: single-cell regulatory network inference and clustering. *Nature Methods*, 14(11):1083–1086, 2017. doi: 10.1038/nmeth.4463.

[59] Philipp Weiler, Marius Lange, Michal Klein, Dana Pe'er, and Fabian Theis. CellRank 2: unified fate mapping in multiview single-cell data. *Nature Methods*, 21:1196–1205, 2024. doi: 10.1038/s41592-024-02303-9.

[60] Cameron Musco and Christopher Musco. Randomized Block Krylov Methods for Stronger and Faster Approximate Singular Value Decomposition. arXiv, 2015.

[61] Fabian Pedregosa, Gaël Varoquaux, Alexandre Gramfort, Vincent Michel, Bertrand Thirion, Olivier Grisel, Mathieu Blondel, Andreas Müller, Joel Nothman, Gilles Louppe et al. Scikit-learn: Machine Learning in Python. arXiv, 2012.

[62] Vincent A. Traag, Ludo Waltman, and Nees Jan van Eck. From Louvain to Leiden: guaranteeing well-connected communities. *Scientific Reports*, 9:5233, 2019. doi: 10.1038/s41598-019-41695-z.

[63] Subhajit Sahu. GVE-Leiden: Fast Leiden Algorithm for Community Detection in Shared Memory Setting. arXiv, 2023.
